# Population-scale inheritance analysis of 858,635 individuals reveals North Sea migration from the Middle Ages to the Industrial Revolution

**DOI:** 10.1101/2025.03.19.643007

**Authors:** Xiaolei Zhang, Poul Holm, Aylwyn Scally, Karina Banasik, Laust Hvas Mortensen, Tomas Fitzgerald, David Westergaard, Piotr Jaroslaw Chmura, Sisse Rye Ostrowski, Erik Sørensen, Ole B.V. Pedersen, Michael Schwinn, Christian Erikstrup, Mie T Bruun, Bitten A Jensen, Henrik Ullum, Christina Mikkelsen, Thomas Folkmann Hansen, DBDS Genetic Consortium, Kari Stefansson, Søren Brunak, Ewan Birney

**Affiliations:** European Molecular Biology Laboratory, European Bioinformatics Institute (EMBL-EBI), Hinxton, UK; Trinity Centre for Environmental Humanities, School of Histories and Humanities, Trinity College, Dublin, Ireland; Department of Genetics, University of Cambridge, Cambridge, UK; Novo Nordisk Foundation Center for Protein Research, Faculty of Health and Medical Sciences, University of Copenhagen, Copenhagen, Denmark; Department of Obstetrics and Gynaecology, Copenhagen University Hospital Hvidovre, Hvidovre, Denmark; Statistics Denmark, Copenhagen, Denmark; Department of Public Health, University of Copenhagen, Copenhagen, Denmark; ROCKWOOL Foundation, Copenhagen, Denmark; Department of Health Technology, Technical University of Denmark, Lyngby, Denmark; Department of Clinical Medicine, Faculty of Health and Medical Sciences, University of Copenhagen, Copenhagen, Denmark; Department of Clinical Immunology, Copenhagen University Hospital, Rigshospitalet, Copenhagen, Denmark; Department of Clinical Immunology, Zealand University Hospital, Køge, Denmark; Department of Clinical Immunology, Aarhus University Hospital, Aarhus, Denmark; Department of Clinical Medicine, Aarhus University, Aarhus, Denmark; Clinical Immunology Research Unit, Department of Clinical Immunology, Odense University Hospital, Odense, Denmark; Department of Clinical Immunology, Aalborg University Hospital, Aalborg, Denmark; Statens Serum Institut, Copenhagen, Denmark; Novo Nordisk Foundation Center for Basic Metabolic Research, Faculty of Health and Medical Sciences, University of Copenhagen, Copenhagen, Denmark; deCODE genetics/Amgen, Inc, University of Iceland, Reykjavik, Iceland; Faculty of Medicine, School of Health Sciences, University of Iceland, Reykjavik, Iceland

## Abstract

The North Sea’s historical migrations have impacted the genetic structure of its neighbouring populations. We analysed haplotype sharing among 858,635 modern individuals from Denmark and Britain to infer migration history from the Middle Ages to the modern day. We estimated the genetic relationship among 370,259 Danes using Danish healthcare registries and validated those with retrievable pedigree relationships from a national family registry. We also compared their haplotype sharing with 488,376 British individuals from the UK Biobank.

Our analysis revealed the fine-grained population genetic history of Denmark and identified distinct coastal and island communities with a history of genetic isolation and bottlenecks. We observed a significant population decline in Jutland compared to Zealand from the late medieval period to the start of early modern period, accompanied by migration from Jutland to Zealand, corresponding to historical evidence.

We identified two major haplotype sharing patterns between Denmark and Great Britain: (1) early coast-to-coast connections between South Jutland and eastern England, likely driven by migration, settlement, continuous trade and the movement of people across the North Sea from the Middle Ages through early modern times, and (2) later city-to-city connections such as those between London and København, likely shaped by the reorientation of economic and trading centres in Britain and Denmark, and the increased shipping from early modern times towards the Industrial Revolution. Further comparisons using other North Sea countries showed both shared and unique histories of genetic exchange with Denmark and Britain.

Our study provides novel genetic evidence of migration across the North Sea from the Middle Ages to the Industrial Revolution. It highlights the power of nationwide biobanks in reconstructing fine-scale historical population movements among closely related populations.

## Introduction

The North Sea area has a fascinating history due to its location and historical interactions. Its rich maritime heritage, including fishing, trade, and wars, have shaped the populations living around and adjacent to its coastline. In this study, we focus on the demographic and ancestral connections between Britain and Denmark, two countries with extensive genetic resources available for responsible research, including historical studies.

Human genetics studies have been previously used to discover and explore historical migrations in the genetic makeup of modern-day populations. The sources of genetic material can either be ancient DNA combined with present-day samples, or just present-day samples alone. Previous studies using present-day samples have investigated the genetic affinity between British and Danish populations (Athanasiadis et al. 2016; Leslie et al. 2015), but being based on only a few thousand samples, had limited spatial and temporal resolution.

From ancient human DNA samples, there is strong genetic evidence supporting the historical migration between Northern Europe, Scandinavia and Britain during the Late Roman and early Medieval periods (Margaryan et al. 2020; Speidel et al. 2025; Gretzinger et al. 2022; Rodríguez-Varela et al. 2023).

However, studies based on autosomal data suggest a limited genetic impact of the Danish Viking settlement (i.e., the ‘Danelaw’) in modern Britain (Leslie et al. 2015), in contrast to studies based on Y-chromosome markers, as well as archaeological and linguistic evidence (Lall et al. 2021; Kershaw and Røyrvik 2016). There are also limited studies of the post-Viking age genetic exchange across the North Sea. It has not been routine to use the extensive genetic resources from modern-day samples to inform the movement and links of groups of people at scale across these timescales. Additionally, Danish populations were previously considered “remarkably homogeneous” (Athanasiadis et al. 2016), and there has not been a large-scale genetic study examining the fine-scale population structure and history of Denmark.

In this study, we assembled 858,635 individuals’ genetic data from two biobanks located in the UK and Denmark, which is at least 100-fold higher than the sample sizes in previous studies. Due to their scale, these data have the potential to provide an unprecedented geographical and time-based view of shared ancestry. At the finest scale that we considered in the study, all NUTS3 (Nomenclature of Territorial Units for Statistics) levels of regions in Britain, at the sub-county level, have more than 40 individuals (median=1,256) born at these locations. Similarly, in Denmark, we can use the municipality level for the Danish sub-structure, with each municipality having at least 73 individuals (median=1,827) (Figure S1). From these data, we identified all shared genomic segments larger than 2cM between all pairs of individuals, each segment inherited from its most recent common ancestor unbroken by recombinations (i.e. haplotype or identity-by-descent (IBD) sharing). Due to recombinations, the length of shared segments reflects the probable age of their common ancestor, with shorter segments more likely to be inherited from older ancestors and longer segments from more recent ones. Thus, these data allow us to identify signals of shared ancestry across different periods.

In total, we analysed a little over 100 billion IBD segments in the 368 billion pairs of individuals to (1) characterise population structure and infer demographic history within Denmark; (2) identify hotspots of IBD sharing between Britain and Denmark across time, stratified by lengths of IBD segments, and (3) analyse the IBD sharing of Britain and Denmark with other neighbouring countries to identify other potential migration patterns. For the non-British or non-Danish European countries, we used proxy populations, comprising people born in and inferred to be from other European countries within our assembled UK and Denmark cohorts. We interpreted our genetic findings in comparison with existing historical evidence, where we both support and challenge aspects of the historical record.

## Results

### Generation of 368.6 billion individual pairs of IBD sharing

We used the large-scale UK BioBank (UKBB) (Bycroft et al. 2018) resource with 488,376 individuals as a sampling of the British population (approximately 0.7% of the total population), and the Danish Blood Donor Study and the Copenhagen Hospital Biobank with a total of 370,259 individuals for a sampling of the Danish population (approximately 6.2% of the total population) (Figure 1 and Methods). We also include individuals from the Faroe Islands in the Danish cohorts to compare their distinct genetic history with that of Denmark (“continental Denmark”), as shown in previous genetic studies (Gislason 2023; Als et al. 2006). For the population of Britain, we include 410,266 samples with self-reported “White British” ethnicity. As there is no ethnicity information available in the Danish administrative datasets, we inferred ancestry using a genetic clustering approach (Methods), and refer to the inferred European samples in the Danish cohorts as “Danes of European ancestries” (n=316,512), representing the population with mainly Danish ancestry (see Methods).

**Figure 1.**
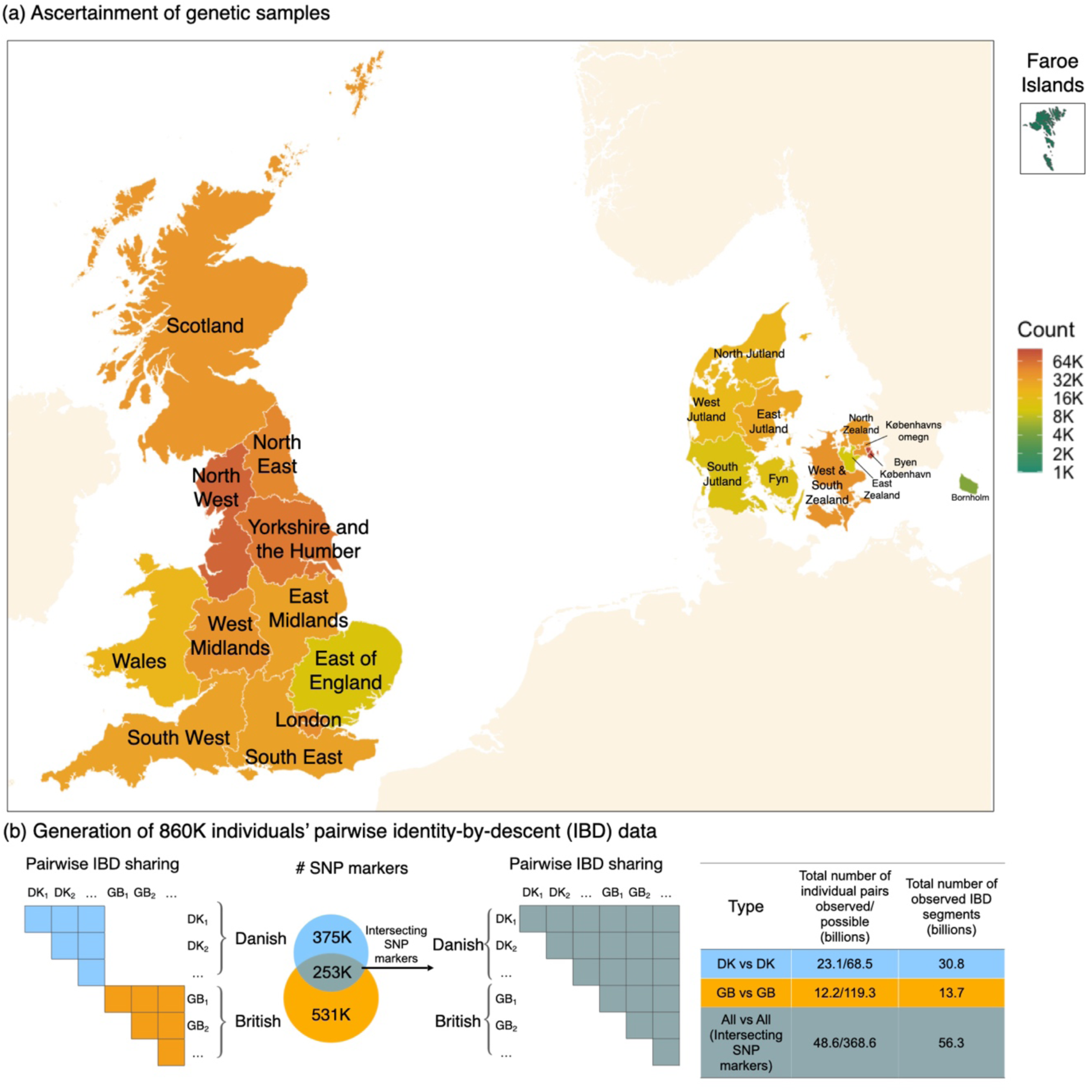
Overview of data collection and study design. (a) Geographic distribution of sample birthplaces and corresponding sample sizes across Britain (including regions of England, Wales and Scotland), Denmark and the Faroe Islands. (b) Diagram for estimating the IBD sharing within the Britain (GB vs GB), within Denmark (DK vs DK), and between the Britain and Denmark (included in the All vs All analysis using the intersecting genotype data), along with a summary of the total pairwise comparisons evaluated and the IBD data generated.

We estimated IBD sharing independently across the samples on three datasets, each defined by a specific set of SNP markers: (1) within Britain, using all SNP markers in the UKBB; (2) within Denmark, using all SNP markers in the Danish samples (DNK); (3) between Britain and Denmark, using the intersecting set of SNP markers shared after combining the UKBB and Danish genotype datasets (UKBB+DNK).

We used FastSMC (Nait Saada et al. 2020) to estimate IBD segments longer than 2cM for all possible pairs of individuals. The total number of pairwise comparisons is substantial, with 368.6 billion pairs of samples evaluated. Across the three datasets, we discovered and processed 100.8 billion IBD segments in total, requiring us to develop a high-performance solution utilising bespoke data structures, parallel computing and indexing to analyse the IBD data.

Here we use the Danish spelling of city place names in Denmark instead of the anglicised names, with the most obvious difference being København (Copenhagen), or the spelling preferred by the local authorities (e.g Aabenraa and Aarhus). For other geographical terms, such as the regional names (e.g. Jutland and Zealand), we use the anglicised spelling for ease of comprehension.

### Overall properties of IBD sharing via 369 billion pairwise comparisons

We focus on IBD segments greater than 2 cM where we are confident that the IBD estimation is sensitive and accurate (see Methods). Figure 2 shows the distribution of IBD sharing among the 860K samples using the intersection SNP marker set. We first considered pairs segregated by their birthplace from the island of Great Britain in the UK (England, Scotland and Wales; Northern Ireland is not considered here due to its low sample representation in the UKBB), Denmark, and the Faroe Islands. We see intra-population IBD sharing is consistently higher than inter-population, with geographic distinct comparisons such as IBD sharing between Britain and Denmark being the lowest (Figure 2 and Table S1).

**Figure 2.**
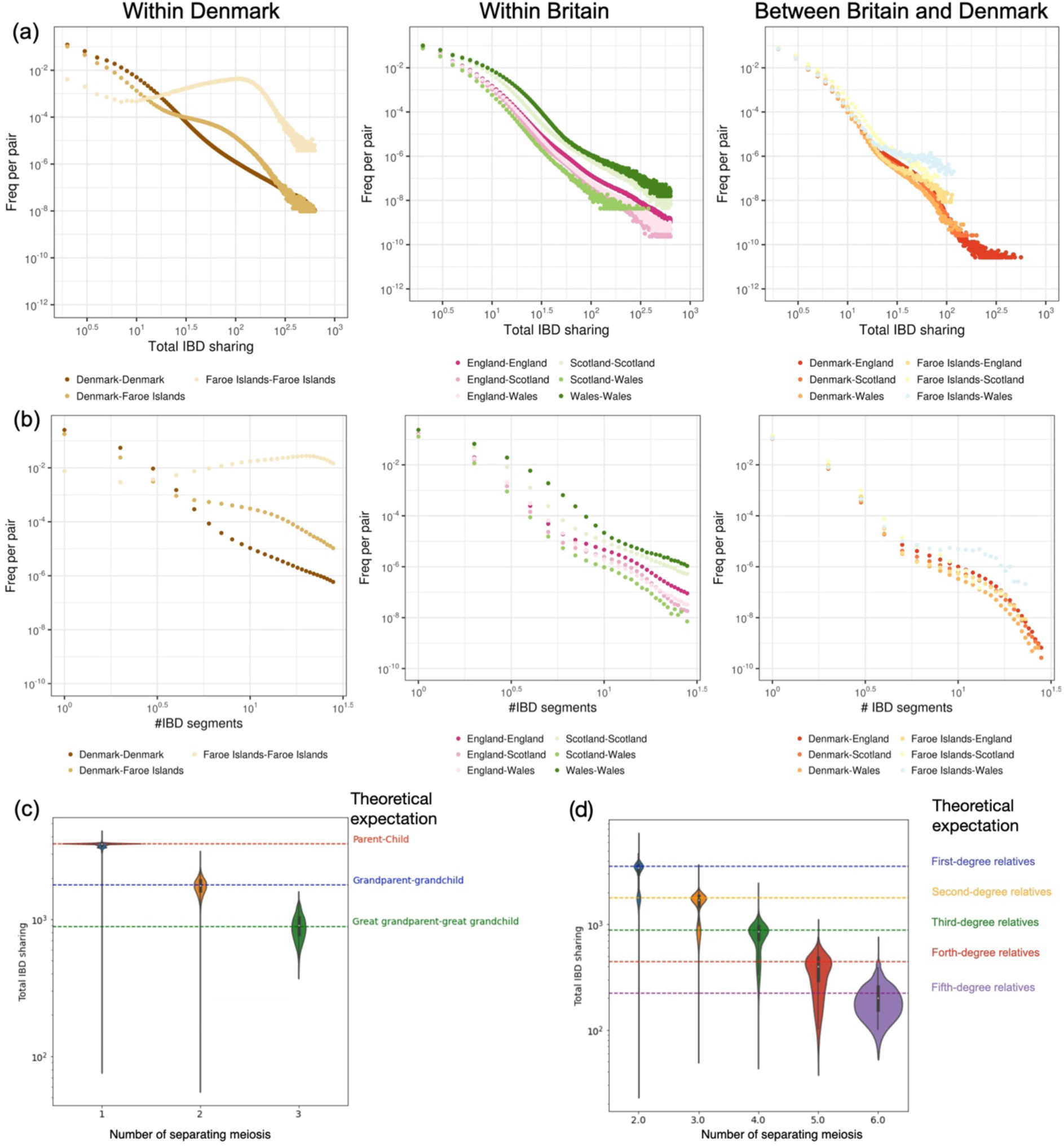
Properties of IBD sharing at population-scale. (a) Frequency distribution of computed pairwise total IBD sharing (>2cM) and (b) total number of IBD segments (>2cM) within Denmark, within Britain, and between Denmark and Britain. (c-d) Relationship between observed and expected pairwise total IBD sharing: (c) among relatives with direct ancestor-offspring relationship, (d) among relatives sharing a common pedigree ancestor, excluding those with direct ancestor-offspring relationship.

Most of the pairs within each of the populations from Britain and Denmark share at most one short IBD segment longer than 2 cM, indicating sharing at most one relatively distant common genetic ancestor. The distributions of the pairwise total IBD sharing and the number of IBD segments are close to exponentially decayed (Figure 2), indicating that the number of detected distant relatives (lower IBD sharing and less IBD segments) increases far more rapidly than that of closer relatives (higher IBD sharing and more IBD segments) in a population. This is consistent with theoretical expectations and with other examples of smaller-scale IBD studies in largely outbred human populations (Huff et al. 2011; Palamara 2014; Henn et al. 2012; Nait Saada et al. 2020; Shemirani et al. 2021).

The situation is different in the Faroe Islands, however, where most pairs of individuals share multiple IBD segments in our length range, with an average of 10.4 segments and 10.7 cM total IBD sharing per pair. These values are 4 times and 41 times higher, respectively, than within Denmark (Table S1). This high level of intra-population sharing likely reflects the fact that this population has been genetically isolated and has gone through genetic bottlenecks, consistent with previous studies on the Faroe Islands (Gislason 2023).

We used the uniquely comprehensive and digitalised Multi-Generation Register Lite (MGR-lite) (Due et al. 2024; Westergaard et al. 2024), a genealogical database of all Danes (n=7,771,981) to reconstruct pedigree relationships among the 370K Danish genetic samples in our cohorts. This family registry documents the relationships between parents (registered as legal guardians) and their children who were alive in 1968 and born from the year 1953, and is partially complete for people born before 1953, after filtering out both reported and likely cases of adoption (Methods). Based on this, we inferred 56,329 pairs of direct ancestor-offspring relatives and 54,089 pairs of relatives with shared common ancestors (excluding direct ancestor-offspring relatives), including both close and distant cousin relationships up to fifth-degree relatives (Table S2 and Table S3). For these 110,418 pairs of individuals, we can infer the total number of meioses separating the two individuals from their pedigree relationship. In Figure 2, we show the distance in separating meiosis compared to the total IBD sharing, which follows closely the expectation from theoretical models (Huff et al. 2011). For parent-child and grandparent-grandchild relatives, we observe long tails on the lower end, suggesting possible unbiological related pairs due to civil registration errors or unrecorded adoptions in the pedigree record, also leading to skewed distributions among shared ancestor pairs. We also observe minor modes in the distributions of IBD in shared-ancestor pairs, indicating cases with single shared ancestors, such as half-siblings or half-cousins. Overall, IBD sharing is exponentially decayed as expected with the increasing number of separating meioses among genetic relatives, thus helping to infer the age of common ancestors in subsequent analyses. For ease of presentation, we assume here an average human historical generation time estimate of 28 years (Moorjani et al. 2016) in subsequent analysis.

### Time-resolved fine-scale population structure in Denmark

There have been several systematic reconstructions of the population structure and ancestry profile of UK populations (Leslie et al. 2015; Nait Saada et al. 2020; Hu et al. 2025), and our datasets of IBD sharing within the UK are consistent with previous work (Figure S6-S7). However, our study is the first to use the large-scale dataset to look at Danish population history from a genetic perspective. As well as using the entire population sample of individuals with European ancestries, we additionally inferred population structure using a refined subset of 28,135 individuals, whom we can identify as having been born in the same provinces as both of their parents using the MGR-lite registry, which resulted in no qualitative difference (Figure S19).

We first considered population sharing across the eleven provinces in Denmark. The geographic division of regions, provinces, and municipalities follows the classification system of Statistics Denmark(“Statistics Denmark” 2007). We find substantial intra-population IBD sharing, and this is most prominent in the rural areas and islands, e.g. Bornholm and Læsø (Figure 3a and Figure S8), with a gradient of decreasing intra-population sharing from Jutland to Zealand (Figure 3a). Within Jutland, North and West Jutland demonstrate higher levels of intra-population IBD sharing compared to South and East Jutland (Figure 3a and Figure S8).

**Figure 3.**
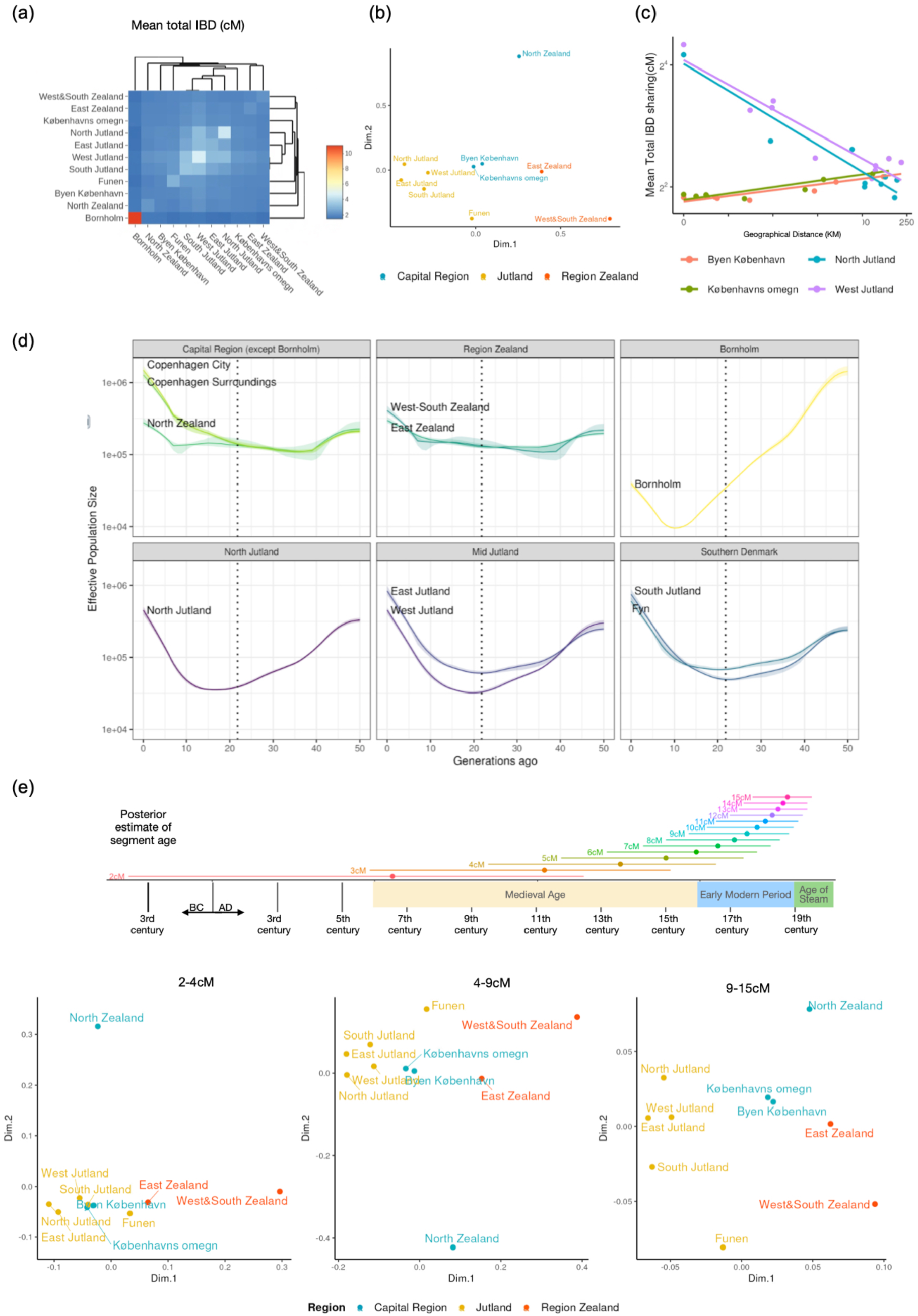
Haplotype-sharing within the Danish population. (a) Pairwise total IBD sharing between provinces of Denmark. (b) Multidimensional scaling on the dissimilarity between provinces of Denmark (except Bornholm) transformed from pairwise total IBD sharing. (c) The relationship between geographic distance (X-axis) from a focal province to all the other provinces (each dot represents one) and their mean total IBD sharing (Y-axis) (except Bornholm). The linear line indicates the fitted linear regression model. We selected Byen København, Københavns omegn, West Jutland and North Jutland as focal provinces here. A full plot including all provinces is shown Figure S16. (d) Effective population size estimate in regions of Denmark across time (up to 50 generations ago) inferred by IBDNe. The dotted line indicates the date of the Black Death (21.6 generations ago) based on a generation time of 28 years. (e) Multidimensional scaling on the total IBD dissimilarity between provinces of Denmark (except Bornholm) stratified by different ranges of IBD lengths. Above, for a range of IBD lengths, we show the age estimate with the maximum posterior probability (dot) and its 50% credible interval (HDI), under a constant population size model.

We also inferred runs of homozygosity (ROHs) within individuals, defined as segments where both parental haplotypes of an individual are identical by descent (>=2cM), thus measuring parental relatedness. Several islands, including Sejerø, Fur, Læsø, and Fanø, have an average amount of total ROHs and number of ROH segments similar to or higher than the Finnish samples (estimated based on Finnish-born individuals from the DNK cohorts) (Figure S10-S11).

We also assessed the ROHs of individuals with other ancestries (predicted European continental component <90%) in the DNK cohorts. We identified municipalities in the Capital Region, especially from Københavns omegn (Copenhagen surroundings), with high levels of mean total ROHs, such as Ishøj, Albertslund, Brøndby, Høje-Taastrup, Vallensbæk and Hvidovre (Table S4). These municipalities are geographically close to each other and are home to large immigrant communities from countries such as Pakistan, Nepal, India, Iraq and Turkey, where some populations have a high rate of consanguinity (“Statistikbank” 2024; Sahoo et al. 2021). Using data from the 1000 Genomes and Human Genome Diversity Project as reference(Bergström et al. 2020; “A Global Reference for Human Genetic Variation, The 1000 Genomes Project Consortium” 2015), we found that individuals with high levels of total ROH (>10cM) in these areas have on average 30%-45% ancestry from Central and South Asia (Figure S13). This is similar to findings in South Asian communities in the UK and elsewhere with high levels of consanguinity (Arciero et al. 2021).

When we consider inter-population IBD sharing, populations from the Jutland Peninsula cluster together and are differentiated from the Islands of Zealand, which is also indicated by the multiple-dimensional scaling analysis using regional pairwise mean total IBD sharing (Figure 3a-b). Focusing on Jutland (i.e., assessing Jutland vs all regions), we find that IBD sharing with other areas of Denmark decreases approximately exponentially with geographic distance (Figure 3c highlights North Jutland and West Jutland; a full figure with other focal provinces is shown in Figure S16). However, Byen København and Københavns Omegn show higher IBD sharing with geographically distant Jutland than with each other or with the neighbouring region of Zealand (i.e. København vs all; shown in Figure 3c). This violates the expectation of an isolation-by-distance model under limited migration (Ralph and Coop 2013), and suggests a history of substantial or extensive migration from Jutland to København, as discussed below.

The relationship between the length of an IBD segment and the time of its most recent common ancestor (TMRCA) is not one-to-one, and is affected by demographic factors such as population size and migration (see Methods). Under certain assumptions the demographic history of a population can therefore be inferred from a distribution of shared segment lengths. We estimated effective population size over time in eleven provinces of Denmark (Figure 3d) using IBDNe (Browning and Browning 2015). Across Jutland, effective population size shows a consistent decline compared to Zealand during the medieval period, particularly in North Jutland and West Jutland, which reached their lowest effective population size 15-20 generations ago (from the end of the 14th century to the mid-16th century). This population decline can also be observed in samples from Aalborg and Aarhus, two of the major cities in Jutland since the Middle Ages (Figure S14). The Black Death in Denmark (1348-1350 CE) occurred within the duration of this estimated population decline or just before the lowest point in inferred population size (“Urban and Rural Survivorship in Pre- and Post-Black Death Denmark” 2021) (21.6 generations ago). Both the Black Death and the emigration associated with it may thus have contributed to the reduction in population in Jutland and Zealand, which we discuss below in relation to historical evidence. This analysis also revealed a distinct demographic history in Bornholm compared to Zealand and Jutland, with a stronger population bottleneck through the medieval and early modern periods, reaching its lowest at 10 generations ago (late 17th century).

To further explore the time dependence of shared ancestry in these data, we stratified IBD sharing by segment length to highlight inter-population sharing across different periods, as visualised by multidimensional scaling (Figure 3e; full time series for each IBD length is shown in Figure S15). Longer shared segments tend to have more recent TMRCAs than shorter ones; however, the uncertainty in these distributions is not well captured by a point estimate such as the mean or mode of the posterior distribution (i.e. maximum a posteriori). Therefore in Figure 3e we illustrate both the maximum a posteriori (MAP) estimate (under a simple demographic model with constant population size, detailed in Methods), and the estimated 50% highest density interval (HDI, i.e. the most credible age range with 50% posterior probability) for a range of IBD lengths, to better illustrate the uncertainty of these estimates. We note also that changes in demographic modelling assumptions can shift these distributions in either direction (see the Discussion on uncertainty in modelling, supported by Supplementary Note).

We observed elevated sharing between København and Jutland over time, compared to København’s sharing with the nearby Region Zealand (Figure 3e). Using København as a focal province, we found a consistent positive correlation between its IBD sharing with other regions and their geographical distance, driven by its high sharing with Jutland (Figure S17). Other areas of Zealand also exhibit higher IBD sharing with Jutland relative to their geographical distance. This is most notable for segments of length 2-9cM (Figure S17-S18), which show a positive correlation between the geographic distance of a focal province to other provinces and their mean total IBD sharing, and have an estimated MAP age ranging from the medieval period to the beginning of early modern period (MAP age = 555AD - 1644AD, 50% credible interval = 257BC - 1773AD). This pattern suggests substantial and continuous migration from Jutland to Zealand Island, mostly evident in København, aligning with the significant population decline in Jutland compared to Zealand we have observed in our effective population size analysis. Meanwhile, North Zealand stands out as an outlier in the Capital Region, showing relatively lower genetic exchange with Jutland (visualised in IBD-length stratified multidimensional scaling plots in Figure 3e) and slower population growth over the last 20 generations (Figure 3c).

Ideally, if we had substantial samples from other European countries, it would help resolve the different possible histories of genetic exchange across Europe with Denmark. In the absence of such data, we constructed proxies of these populations from other European countries by using individuals in the UK and Danish datasets who were born in and inferred to have substantial ancestry from these countries. This inference was made by using a Leiden community detection approach (Traag, Waltman, and van Eck 2019) based on individual pairwise IBD sharing in our combined UKBB and DNK datasets (UKBB+DNK) (see Methods).

Using these data, we estimated the IBD sharing between the Danish populations and the other neighbouring countries (excluding sharing with Britain, which we explore in more detail below). We find the pattern of IBD sharing is strongly associated with geographic proximity and historical evidence of migration (Figure 4). Norway and Sweden have the overall highest IBD sharing with Denmark (Figure 4). Norway is most closely related to the North Jutlandic population, while Sweden and Finland are most related to the eastern side of Zealand, especially North Zealand, and Bornholm. This is in contrast with Germany and the Netherlands, which are most related to the southern border of Denmark at the Jutland Peninsula. Poland shows the highest IBD sharing with the Lolland-Falster islands in the south of Zealand. For Bornholm, compared to Jutland, it has substantially higher sharing with other Baltic countries, including Sweden, Finland and Poland (Figure S21). The results are largely qualitatively the same if we only use individuals born in other countries from either of the cohorts (Figure S22).

**Figure 4.**
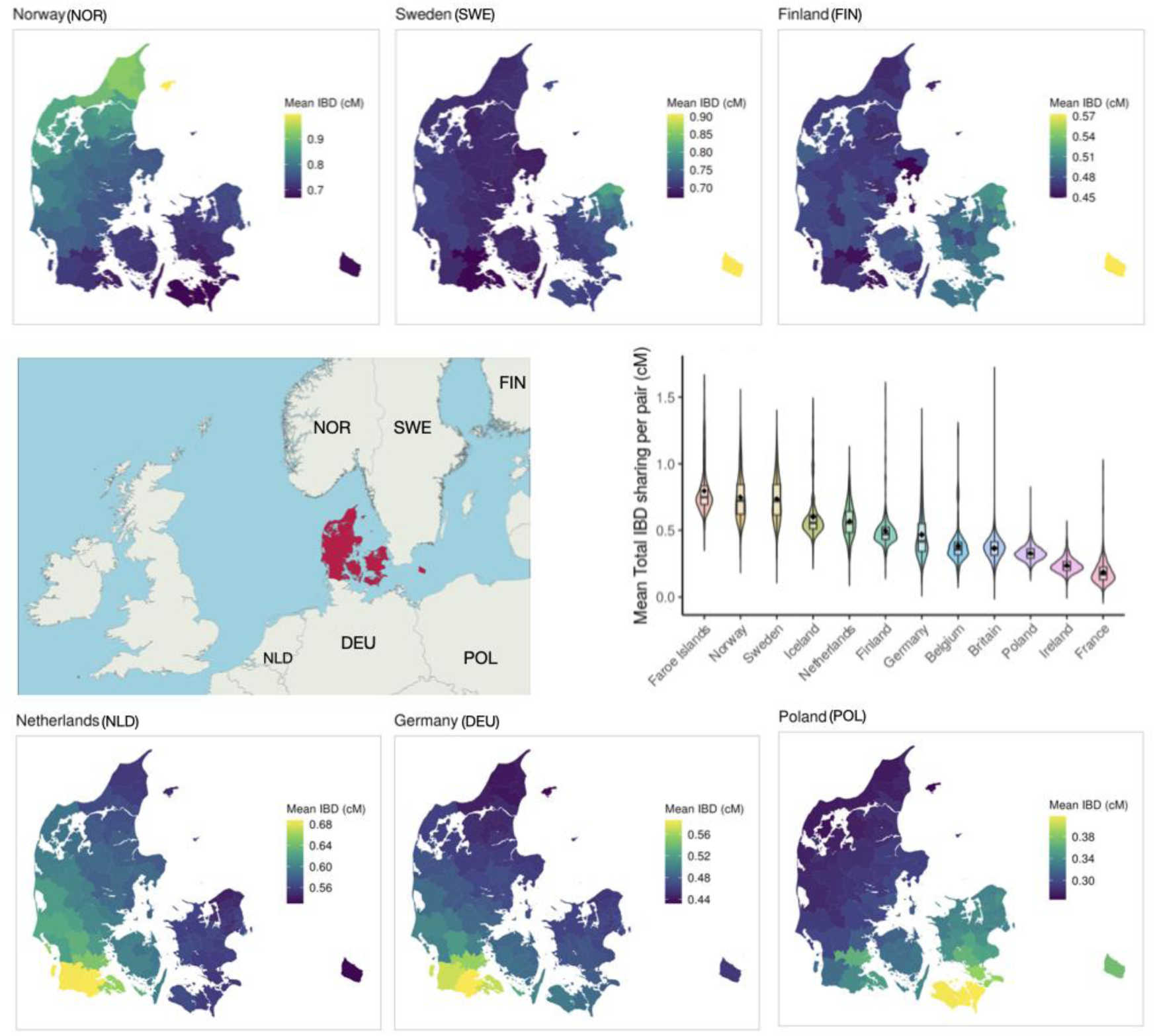
Mean total IBD sharing between Denmark and neighbouring European countries. To capture heterogeneity among samples, we calculated the mean total IBD sharing between each sample from neighbouring European populations and all samples from Denmark (i.e. one vs all comparisons). The violin plot displays the distribution of mean total IBD sharing across individuals (one vs all) in each compared population. The embedded box plot indicates the median, lower (25%), and upper (75%) quartiles, while the dot represents the mean total IBD sharing for each European population with Denmark. Norway, Sweden, Finland, the Netherlands, Germany, and Poland are highlighted to show their sharing with different regions in Denmark. The colour scale of each highlighted panel is normalised independently.

### Haplotype-sharing across the North Sea

A unique aspect of our study is the ability to look at two substantial regions of Europe, which have documented trading and historical links and have accessible genetic resources for research. We analysed 180.8 billion pairs of individuals between Britain and Denmark using the intersection of the two genotyping arrays (as outlined above and in the Methods). Between Britain and Denmark, 9.5% pairs (17.2 billion) share at least one IBD segment larger than 2cM, likely corresponding to shared genetic ancestry within the last two millennia. We find that the high IBD sharing with Britain is significantly clustered at the west coast of Jutland populations (Figure 5, Figure S24, local Morans’ test *P*-value <0.05; for hotspots/statistically significant clusters, where high values are surrounded by other high values, defined with *P*-value <0.05), in particular the southernmost part of South Jutland - Tønder and Aabenraa municipalities (Figure 5b, local Moran’s test *P*-value=0.004). The geographic distribution of total IBD sharing in Denmark is similar when compared separately with the three British populations at the national level: English, Welsh and Scottish (Figure S23). Within Britain, the East Midlands (particularly Derbyshire, Nottinghamshire, Leicestershire, Rutland and Lincolnshire) and South Yorkshire have the most IBD sharing with Denmark (Figure 5, Figure S25, local Moran’s test *P*-value<0.05), in particular South and West Derbyshire with the strongest positive spatial autocorrelation (local Moran’s test *P*-value=0.003).

**Figure 5.**
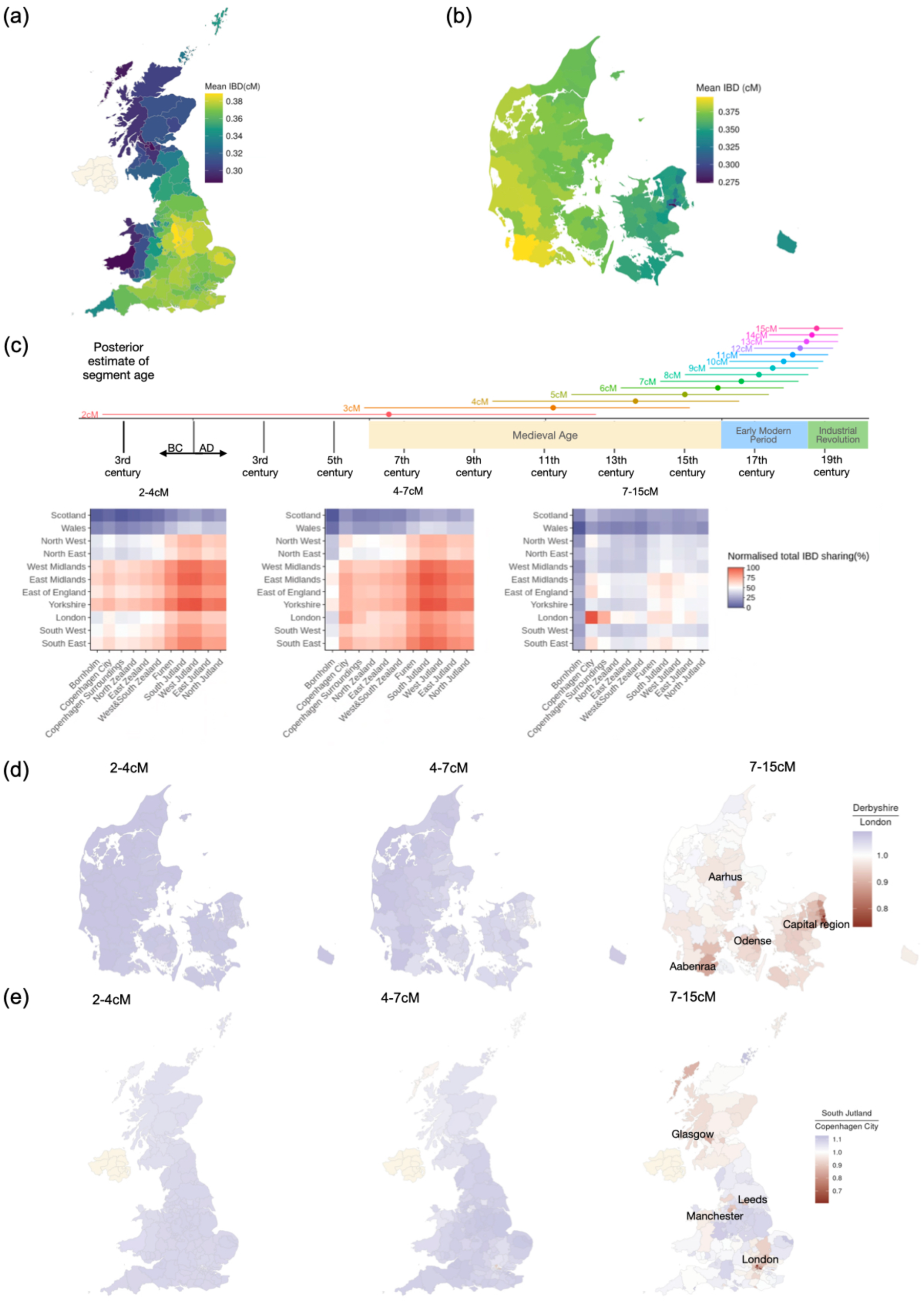
IBD sharing across British and Danish populations. (a).The geographic distribution of mean total IBD sharing between Denmark and regions of Britain (at NUTS3 level). (b) The geographic distribution of mean total IBD sharing between Britain and the regions of Denmark (at the municipality level). (c) The IBD sharing across British and Danish regions across time periods through stratification using the length of IBD segments. For each IBD length, we show the age estimate with maximum posterior probability and its 50% credible interval (HDI). Each heatmap is normalised independently (scaled the range to be between 0 and 1). (d) The contrast, expressed as a ratio, in the geographic distribution of IBD sharing between the Danish population and Derbyshire versus the Danish population and London, stratified by IBD length. (e) The contrast of the geographical distribution of IBD sharing between the British population and South Jutland versus the British population and København, stratified by IBD length.

Similarly to the intra-Danish analysis, we then stratified the analysis into different IBD length intervals, to discern patterns of shared ancestry from different time periods (full series is available in Figure S26-S27; Note that the three length intervals in Figure 5 are to chosen illustrate the patterns we see in this analysis, and are different those used to explore shared ancestry within Denmark.). We see striking differences in IBD sharing between Britain and Denmark across these three length intervals. Short IBD segments (defined as 2-4cM, MAP age = 555 AD - 1255AD, 50% credible interval =257 BC - 1549AD) show the strongest sharing between the East Midlands and Yorkshire in the England, and South Jutland and West Jutland in Denmark (Figure 5c, Figure S24-S25, local Moran’s test *P*-value<0.05). Notably, increased sharing with East Midlands and Yorkshire is also observed in other regions of Denmark. For IBD between 4-7cM (MAP age = 1255AD - 1555AD, 50% credible interval = 849AD - 1717AD), we see a similar pattern, with elevated sharing between South Jutland and the east coastal area of England including East of England and South East, in particular Norfolk (Figure S25 and S28, local Moran’s test *P*-value<0.05).

By contrast, the longer IBD segments, corresponding to more recent ancestry (7-15cM, MAP age=1555AD-1768AD, 50% credible interval = 1325AD-1843AD), show the highest sharing between London and København (Figure 5c and S25, local Moran’s test *P*-value<0.05). Using a fine-grained geographic resolution, we find the two capital cities also both show elevated long IBD sharing relative to their short IBD sharing patterns with other cities across the North Sea, i.e. higher sharing between London and København, Aabenraa, Odense, Aarhus compared to Derbyshire with those Danish cities, and higher sharing between København and London, Greater Manchester, Leeds and Glasgow compared to South Jutland to those British cities (Figure 5d and Figure 5e). For the cities in Jutland Aabenraa, Odense and Aarhus, apart from the link with London, we also observe elevated mid to long IBD sharing between the east coastline of England including Norfolk (East Anglia) and South East (Figure S30).

We also estimated the IBD sharing between Britain and other neighbouring European countries, similar to the Danish analysis (Figure 6). At the country level, we find Ireland has the overall highest sharing with Britain, which is unsurprising given the proximity and entangled history of Ireland and Britain. Western Scotland, particularly Glasgow and the Highlands and Islands, and Liverpool from the North West of England have the most mean total IBD sharing with Ireland in our data. Glasgow and Liverpool have a marked increase in mid and long shared IBD segments (Figure S42-S43). Norway, Iceland and the Faroe Islands all show a similar geographical distribution of IBD sharing with Britain, in which the Western Isles, Orkney and Shetland in Scotland show the highest overall IBD sharing across all IBD lengths (Figure S35-S37). This likely reflects substantial Norwegian influence in these regions during the medieval period (Margaryan et al. 2020; Als et al. 2006; Jorgensen et al. 2004; Helgason et al. 2000; Ebenesersdóttir et al. 2018).

**Figure 6.**
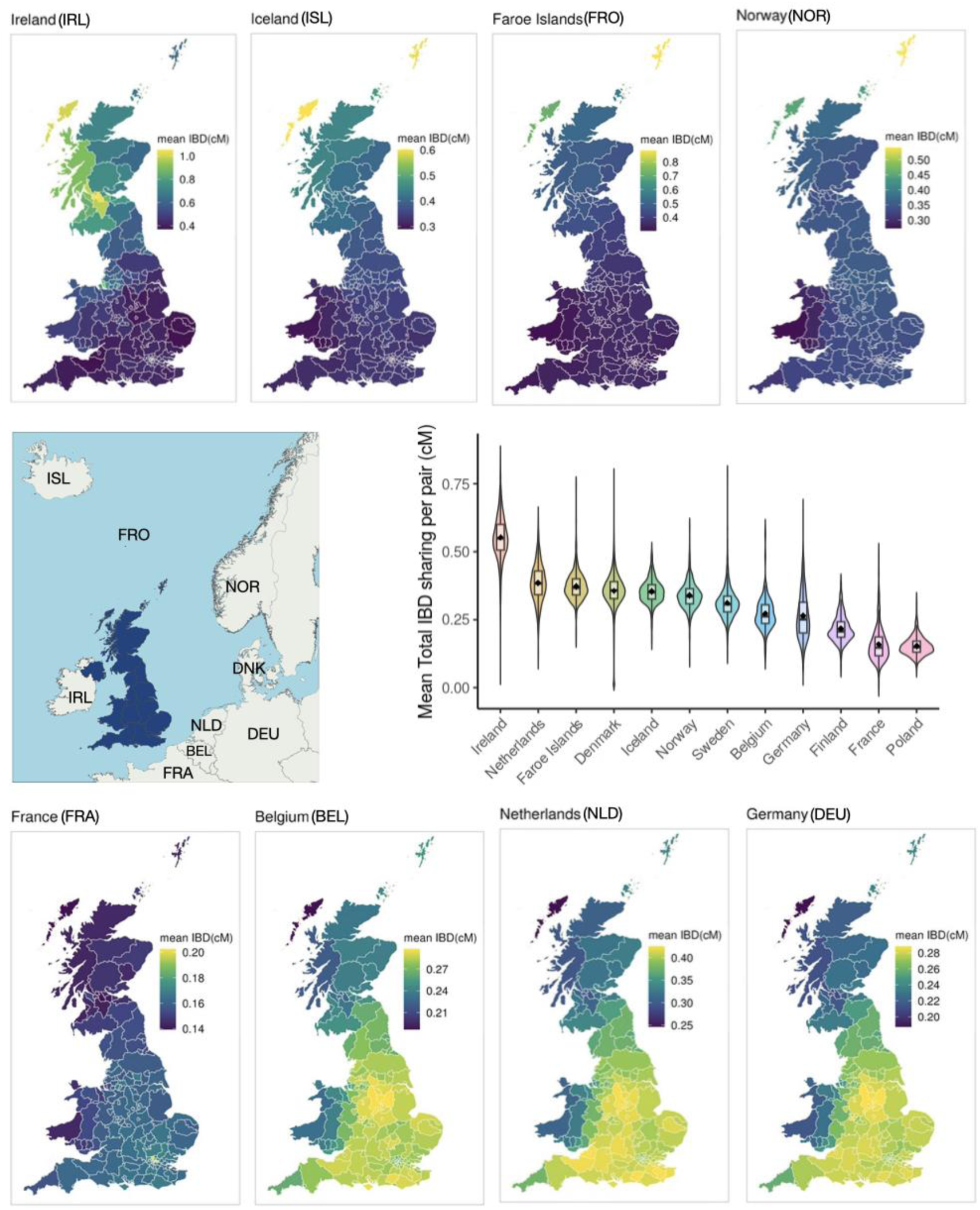
Mean total IBD sharing between British and neighbouring European populations. Here we calculated the mean total IBD sharing between each sample from neighbouring European populations and all samples from Britain (i.e. one vs all comparisons). The violin plot displays the distribution of mean total IBD sharing across individuals (one vs all) in each compared population. The embedded box plot indicates the median, lower, and upper quartiles, while the dot represents the mean total IBD sharing for each European population with Britain. Ireland, Iceland, the Faroe Islands, Norway, France, Belgium, the Netherlands, and Germany are highlighted to show their sharing with different regions in Britain. Each highlighted panel is independently normalised.

After Ireland, and particularly on the eastern and southern side of the North Sea, the highest IBD sharing is with the Netherlands (Figure 6). Derbyshire shows high IBD sharing with the Netherlands, Belgium and Germany for 2-4cM IBD (Figure S38-S40), as well as with Norway across England (Figure S35, showing Derbyshire and the nearby areas have higher IBD sharing with Norway compared to other parts of England), following a geographical dispersal pattern at this IBD length stratification similar to that with Denmark (Figure S28). However, in contrast with Denmark, the Netherlands also has a comparatively high IBD sharing with Kent in the South East of England consistently across the IBD lengths analysed. At longer IBD lengths, sharing with the Netherlands and Germany shifts towards London and the surrounding areas along the South East coast of England (Figure S43). This aligns with what we have observed in the mid to long IBD sharing with Denmark. There are surprisingly low levels of mean total IBD sharing with France relative to other countries given its geographic proximity. The distribution of the mean total IBD sharing between each French sample and all British samples (one vs all), has a long tail on the higher end, so within the French samples there could be one or more subsets with higher sharing (Figure 6). Overall, samples from France predominantly show the high IBD sharing with London and other parts of South East England (Figure S41), with an increasing degree of sharing with London in recent generations (Figure S43). This signal is robust to whether the proxy individuals for France come from either UKBB or the Danish cohorts, and whether we use Leiden clustering of ancestry or not (Figure S33).

### Comparison to historical evidence and synthesis

The fundamental phenomenon underlying our IBD analysis is shared ancestry between the individuals in our cohorts. When two individuals born nearby share IBD segments, the simplest hypothesis in many cases is that their common ancestors were also likely distributed in the same locale. However, alternative hypotheses, such as mass migration from another location, are also plausible (for example in the case of IBD within an immigrant community). Note that here and in what follows we use ‘migration’ to refer to the movement of individuals and their associated genetic ancestry from one location to another. However it is important to consider that many economic, cultural and political factors may have been involved, and in different historical circumstances such movements may have been either voluntary or forced, or a complex mixture of the two. Moreover, when two individuals born in geographically distant locations share IBD segments, multiple hypotheses can explain this sharing. These include direct migration between the two locations, the migration of a related group from a third location to both distant locations, or temporary cohabitation in shared locations where offspring were produced, without long-term settlement. All these scenarios and their combinations over time likely contributed to the patterns we observe, and the IBD data reflect the sum of all these genetic exchanges. Here, we compare IBD data with relevant historical evidence (archival and archaeological, abbreviated as HIS below, derived from a diversity of sources ranging from census to proxy data), bearing in mind that comparisons to known or suspected historical events are complex, particularly demographic events, as these often rely on information about a limited set of high-profile individuals or proxy data of disputed relevance.

Our IBD analysis suggests a significant demographic event occurred in medieval Denmark, altering both population size and migration patterns within the country. IBD-based effective population size estimates indicate a greater population decline in Jutland compared to Zealand during the medieval period, reaching its lowest point 15-20 generations ago. This coincides with the aftermath of the Black Death (1349-50) and subsequent outbreaks of pestilence which continued into the next century (Hybel and Poulsen 2007). While birth and death records from this period are largely absent, other historical evidence points to a population decline in this period. For example, historians have identified a late medieval decline in the number and density of churches in Jutland, with at least 80 churches abandoned in the two centuries before the Protestant Reformation in the 1530s, 75 in Jutland and 5 on the islands of Zealand, Funen, Lolland and Møn (Christensen 1938). In 1200, 63.3% of all Danish churches were located in Jutland, and by 1500, the Jutland share was down to 50.0% (H. Nielsen 2010). Thus, a possible hypothesis is that Jutland was hit much harder by plague and its demographic consequences (discussed below) than the rest of Denmark. While HIS indicates continued migration from Jutland to Zealand after the 15th century, there is limited HIS indication regarding such migration before the Black Death, making earlier migration harder to assess.

The genetic findings indicate potential migration flows within Denmark during the medieval and early modern periods. Modern Zealanders share ancestry with individuals across Denmark, whereas the Jutland population is relatively more homogeneous, with IBD (and thus perhaps most ancestors) being more locally dispersed. Historical evidence shows that degrees of servitude, enforced as a result of the loss of labour after the Black Death, varied considerably. In 1446 at the latest, servitude (vorned) was enforced on adult males in rural areas of Zealand, which tied males to the parish (women could move freely). In Jutland, there was no similar restriction of male mobility, and males are known to have migrated to Zealand in search of better opportunities and better farms (Olsen 1950-2, 203). Farmhands from Jutland (‘jydekarle’), for example, were employed on manors all over Zealand. This may explain why København has higher IBD sharing with Jutland than the nearby regions in Zealand. In the 1699-1705 register of citizens, 19.9% of people in København stated their birthplace as Jutland, and 13.5% originated from Zealand (Olsen 1932, 95).

The IBD and HIS data reveal what appears to be a puzzle: it would seem logical to expect that Jutland, having suffered a severe population decline in the late medieval age, would then have experienced immigration to repopulate deserted farmlands. However there seems little indication of this hypoththesis in the historical or genetic evidence. One alternative hypothesis is that the population decline may have been followed by a shift to agricultural systems that required less labour. For instance, the rise of the north and west Jutland cattle trade, which became a major export trade in the fifteenth century (Enemark, 2003), led to the abandonment of large areas of intensive agriculture in favour of a more extensive pastoral system (Gissel 1981). Pastures yielded at most a third of the food of grain fields (Porsmose 1988), and as many of the cattle were exported, Jutland may have experienced food shortages that drove continued emigration. Another possibility is environmental change. For example, the coastal areas of western and northern Jutland, where we have observed the strongest population decline, have long been characterised by infertile, sandy soils and have periodically suffered from sand drifts between the 6th and 19th centuries, making traditional farming challenging and less productive (Gravesen 2001; Strunge et al., n.d.; Binderup, n.d.; N. K. Nielsen 2023). Another contributing factor may have been repeated military marching and ravaging along the Army Road (Hærvej) in central Jutland (major events are known between the 12th to 17th centuries), which became severely depopulated (Teitsson 1981). Finally, we emphasise that IBD-based effective population size estimates do not account for migration, and thus potentially underestimate Jutland’s historical population size.

The islands around Denmark have varied histories, reflecting different degrees of dependence on trade, agriculture, fishing and international shipping. For Bornholm, the IBD analysis estimates a striking decline from the medieval period until the late 17th century. In Viking times, the island was very wealthy, as evidenced by the archaeological discovery of over 100 silver hoards across its small area (T. Ingvardson 2024). This is much more than nearby regions like Scania and Zealand, despite their larger size. Bornholm’s wealth came from its strategic position on the trade routes between western Scandinavia and the Baltic/Slavic region. Around 900 AD it was an independent kingdom, but it likely came under Danish rule by 1100 AD and may have lost its influence to Gotland. Bornholm’s population decline likely started early and may have been worse than in other Danish regions, due to the loss of its trading role and the effects of the Black Death. However, the population began to grow again in the late 1600s when the island gained strategic value for Denmark after it lost both Gotland and Scania to Sweden. In the 18th century, Bornholm developed a considerable international commercial fleet. This demographic trajectory is largely consistent with the patterns observed in the IBD data.

The islands of Sejerø and Fur relied on a mix of agriculture and local fishing, while the islands of Læsø and Fanø have a strong maritime past. It might therefore be assumed that the strong international orientation of shipping in these islands would show up in present-day genetic ancestry there. In our IBD analyses the majority of the island communities have high amounts of intra-population IBD sharing and ROHs, consistent with a small historical effective population size. Common to all four islands is the fact that the mean total ROH per individual is five or more times higher than the Danish national mean. Thus the cultural and trading differences between these islands appear to have had little impact on the diversity of ancestry on these islands. Pontoppidan (1768, 768) and Kyrre (1912, 30) report the aversion of people on Fur to immigrants from the mainland. On the island of Læsø, there is historical evidence that the surplus of females caused by high male mortality at sea was compensated by migrants from Jutland between the 18th to 19th centuries who were held in low esteem (Stoklund 2018, 241-245) but if so this does not seem to have substantially changed the ROH levels compared to the more agricultural Sejerø or Fur islands.

North Zealand is a consistent outlier with relatively low genetic exchange with other regions of Denmark in our multidimensional scaling analysis of IBD sharing, which otherwise recapitulates much of Danish geography (except København, the capital city and its surrounding area, where immigration seems to be from all other regions, as discussed above). North Zealand’s history is unique in several ways: it was densely forested and depopulated to keep large forests and pastures for the king’s hunt, and soil productivity was at the low midwest Jutland standard (<10 barrels of hartkorn per km^2^) (Dam & Jakobsen 2008, 106). Helsingør (Elsinore) had a large immigrant population, as this was the entry point to the Baltic and the place all ships called to pay the Sound Toll (the Sound or Øresund, the main trade route between the North Sea and the Baltic sea, bordering Denmark and Sweden). Between 1556 and 1575, 50-60 (10-13%) of male citizens of Elsinore were of Scottish or English descent (Tønnesen 1985, 77). In later years, Dutch and German immigrants became more prominent. But Elsinore retained a strong English-Scottish minority, which was reinforced by a new influx of English merchants around 1700. In 1801, the British colony in the city amounted to 100 people, out of a total population of 5,063 inhabitants. The British component stood out with their wealth, and the city was known as ‘Little England’ (Appel 2012). We looked for genetic evidence of this connection with Britain, but did not identify particular signals driven by exchange between Britain and North Zealand. Helsingør also did not show increased IBD sharing with British samples, compared to other municipalities in North Zealand, for long and mid-range IBD segments.

IBD sharing between Denmark and neighbouring countries (Figure 4) follows relatively predictable patterns, at least for longer segments corresponding to periods with more historical evidence for migration and trade. North Zealand has a higher IBD sharing with Swedish and Finnish relative to other parts of Zealand, which may also contribute to it being an outlier in Zealand in IBD sharing patterns. North Jutland has a distinct history of Norwegian immigration, which is well-documented by eighteenth-century records of individual movements and the 1850 census and was strongly related to a grain-for-timber trade, which was a major regional activity between c.1600 and 1870 (Holm 1991, 155-157). Conversely, cultural flows and potential migration from Jutland to southern Norway during the 17th century are also documented (Engelstad 1946). Dutch and German immigration appears to have been more or less continuous from medieval to modern times, originally related to shipping and whaling, and in the 20th century to agricultural investment (Guldberg 2020). Until 1920, South Jutland was part of the wider Schleswig-Holstein region of mixed Danish-German populations. Lübeck, located in this region, and its nearby city Hamburg, have been important trading centres in Northern Europe connecting the North Sea and the Baltic Sea since the 13th century. We would therefore expect people from Northern Germany to share a substantial component of ancestry with Danes from South Jutland; however we lack exact birthplace location for individuals born in other European countries to corroborate this. Poland has the highest IBD sharing with the Lolland and Falster island populations in Denmark, consistent with the history of Polish seasonal workers who settled there to work in sugar beet fields (“Migration Museum of Denmark” 2024). Bornholm shows higher ancestry sharing with other Baltic countries, including Sweden, Finland, and Poland, in comparison with Jutland. IBD with Sweden in particular is consistent with the fact that until 1660, Bornholm was part of the province of Scania, which also includes the southernmost part of Sweden. As well as these documented connections via migration and trade, there is likely a set of earlier migrations from closely related countries, giving rise to shorter IBD segments.

The historical circumstances of migration into and from Denmark changed in 1660, after which Denmark became a protectionist economy under an absolute monarch, with considerable control of migration. In a review of the evidence of international migration after 1645, Johansen (Johansen 2002) concluded: “The total number of such immigrants was probably only a few thousand over a long period, and by 1775 the foreign element in the Danish population was, no doubt, very modest.” Migration in the 18th century involved mostly Norway, Germany, and the Netherlands. For the medieval period, when there were looser controls on migration and the economy was essentially open, lists of burghers (urban residents in towns) document a substantial component of German and Dutch citizens in Danish cities, while Scottish and English immigrants are documented mainly in the town of Helsingør/Elsinore as mentioned above. In the early medieval and Viking period, flows between Denmark and England in both directions are well-attested in the historical record, but there is no consensus on quantitative estimates (McLeod 2014).

The genetic data suggest changes in migratory flows across the North Sea. We observe a striking shift in Danes-British IBD sharing over different ranges of IBD length. The highest level of short IBD sharing (2-4cM) is between South Jutland in Denmark and Derbyshire, Leicestershire, Nottinghamshire, Lincolnshire, Rutland, and South Yorkshire in Britain. Other regions in Denmark show higher sharing with the East Midlands and Yorkshire. Although our analyses of the Dutch, Belgian and German proxy populations are less well-powered, they similarly show higher levels of short IBD sharing with the same British regions as well as Norwegian with English, suggesting a potentially ancient shared population structure connecting these North Sea coastal populations.

Both the temporal and geographical distribution of short IBD across the North Sea point to population movements from continental North Europe into Britain between the fifth to eleventh centuries. The posterior age distribution of short IBD segments (2-4cM; MAP age = 555AD - 1255AD; 50% credible = 257BC - 1549AD) spans both the early medieval ‘post-Roman’ centuries, traditionally regarded as the period of English (Anglo-Saxon) settlement in Britain, and also the era of Danish Viking occupation in England (i.e. the “Danelaw”). Anglo-Saxon control eventually extended across southern and eastern England, whereas the Danelaw covered all of eastern England between the River Tees and the Thames, with York as its Capital (Leslie et al. 2015; Hadley 2006). South Jutland in Denmark and northern Germany lie largely within or near the homelands and centres of activity of the Angles, Saxons and Danish Vikings, as evidenced by archaeological records (Kershaw and Røyrvik 2016; Gretzinger et al. 2022). While our short IBD sharing signal is consistent with previous genetic evidence supporting the migrations from continental northern Europe to eastern England from the later Roman to the middle Anglo-Saxon period (Gretzinger et al. 2022), it could also reflect genetic contributions from the later Danish Viking migration and settlement. The Viking Age Danish towns of Hedeby (in present-day Schleswig-Holstein, northern Germany) and Ribe (in South West Jutland, Denmark) were among the most important trading centres of the period. Both have shown substantial archaeological links with the Danelaw region in England (J. Kershaw and Røyrvik 2016; J. F. Kershaw 2020), supported by linguistic evidence (Fellows-Jensen 2008) and Y-chromosome markers studies (Lall et al. 2021), which align with the spatial distribution of our short IBD sharing data. Given the overlapping origins of Anglo-Saxons and Danish Vikings, as well as the continuous genetic exchange with neighbouring populations, it’s likely that the modern South Jutland population carries ancestry from both groups. Hence, both migration events are likely to have contributed to the genetic signals we see.

Although Derbyshire and the surrounding area lie within the regions that were historically under Saxon and Danelaw control, it is nonetheless intriguing that this specific area is a significant hotspot of short IBD sharing with Denmark. Several explanations may account for this. Firstly, the East Midlands and nearby Yorkshire might have been settled more extensively during these medieval migrations. Secondly, these regions might have had continuing or new demographic events such as trade and population movement lasting beyond the period of Anglo-Saxon and Danish Viking migration. Anglo-Danish trade continued despite the political reorientation of England after the Norman conquest (Sawyer 1986). It is known that Danish merchants were allowed to stay for a year and visit inland fairs, and grants of safe-conduct are known from Newcastle (North East, England), Lynn (East Anglia, England), Yarmouth (East Anglia, England), and London through the 12th and 13th centuries (Childs 1998). The toll records of shipping through the Øresund document that the Baltic trade had strong participation of skippers originating in Norfolk (East Anglia, England), the Firth of Forth (Edinburgh, Scotland), Hull (Yorkshire, England) and London (Veluwenkamp, Scheltjens, and Van der Woude 2021). Finally, other population movements in Britain afterwards may have diluted the signal in other regions more extensively, especially in the urban areas compared to the largely rural western and southern Derbyshire(“Rural Urban District” 2019). Thus, the IBD sharing between hotspots could be a residual signal after subsequent demographic events elsewhere. These explanations are not mutually exclusive - some mixture of all of these might have occurred.

Continued genetic exchange between Britain and Denmark after the early medieval period is also supported by the increased sharing between Norfolk and South Jutland through mid-length IBD (the figure panel of 4-7cM in Figure S28 and Figure S30; MAP age=1255AD-1555AD; 50% credible interval = 849AD - 1717AD), aligning with the medieval history of trade between King’s Lynn (Norfolk), Ribe (South Jutland, Denmark) and the nearby Hanseatic towns Lübeck and Hamburg in Northern Germany(Lloyd 2011; Holm 2002).

The transition towards long-IBD sharing between the cities of Britain and Denmark likely reflects a shift in the economic and trading centres of both countries, alongside the well-documented increased shipping between them from the mid-16th century onwards (Møller 1988). Potentially, the increased ease of transport and corresponding movement of workers, as well as trade-associated movement, lead to a greater incidence of genetic exchange between cities. However, it is notable that some other cities with strong maritime or industrial histories such as Bristol, Newcastle, Brimington and Southampton, whose populations are well ascertained in the UK Biobank, do not have the same level of sharing with København as London, Manchester, Glasgow, or Leeds (Figure S31), suggesting heterogeneity in the shift of genetic exchange. The genetic links between London and the eastern island towns of København, Aabenraa and Odense are consistent with strong migratory links between London and the urban region of Southern Denmark around 1800 (Brock and van Lottum 2019). Again however, it is important to stress that such patterns are also consistent with migration from a third location to both the British and the Danish cities.

We also observe that the Dutch, German and French samples have increased IBD sharing with London in longer IBD segments (corresponding to early modern times and more recent centuries, including the Industrial Revolution) (Figure S42-S43). Although we are unable to consider geographical heterogeneity within these countries and to pinpoint the particular regions driving these links, based on our findings in Denmark, trade involving the coastal and urban populations, particularly from regions with trading links to London, could be one of the contributing factors. For example, Hamburg, Lübeck, Bruges and port cities in the Netherlands (e.g. Dordrecht) were major trading hubs of the Hanseatic League with connections to London. Compared with elsewhere in continental Europe, the comparative peace and economic development of England in the late medieval period attracted immigrants for trade and work. The historical tax records of 64,000 foreign residents in England between the 14th and 16th centuries (“England’s Immigrants 1330 – 1550,” n.d.) show that these immigrants were mainly servants, merchants, and craft people. While such immigration may have occurred continuously at a small scale until the industrial revolution, pulses of migration could have contributed to the observed IBD sharing signals. For example, Norman-French immigrants replaced most of the previous English landowners following the Norman Conquest in 11th century(*Domesday Book* 2003), settling mainly in the south and east of England, and more recently the migration of of 40,000–50,000 French protestants (“Huguenots”), mostly to London, around 1670 CE(Gwynn 1985).

## Discussion

In this study, we performed an extensive identity by descent (IBD) inference between two modern populations, Denmark and Britain, that have experienced complex patterns of genetic exchange, both internally, between each other and with other countries. To achieve this, we analysed shared genomic segments in 369 billion individual pairwise comparisons, and developed space- and time-efficient methods to store and analyse the results. Using a large and accurate database of Danish pedigree information, we showed that the amount of IBD sharing we observe closely aligns with theoretical expectations. The large-scale patterns of intra- and inter-country genetic sharing observed in Britain and Denmark revealed important aspects of Denmark’s demographic history, and episodes of gene flow between the two countries from the Middle Ages through to the Industrial Revolution. In particular, we identified significant migration from Jutland to Zealand, coupled with a population decline in Jutland during medieval period. Our findings also reveal the fine-scaled geographical distribution of the genetic origins and legacy of medieval migration across the North Sea, potentially associated with Anglo-Saxon and Viking migration. We also detected distinct genetic evidence of post-Viking Age migration connecting maritime centres in Britain and Denmark, likely driven by ongoing trade and population movements. Analysis of IBD sharing between Denmark and Britain and other neighbouring countries revealed further shared and population-specific migration patterns.

There are limitations to our study. Our analyses rely on the sampling of people in the current era. The ascertainment of the cohort will often have geographic and volunteer bias, both of which have been shown in UK Biobank (van Alten et al. 2024; Fry et al. 2017). However, the recruitment centres used for UK Biobank are spread across the country, and many places that might have had a strong exchange, e.g, Newcastle, are well sampled. The Danish cohort is a combination of a hospital-centred in- and outpatient cohort in København, with a natural bias towards old-age populations in the region and the Danish Blood Donor Cohort, which is across Denmark, but biased towards healthy volunteers (Hansen et al. 2019). Although we would have preferred a more even ascertainment in Britain and Denmark, we do not believe that the analysis of IBD sharing patterns presented here is significantly skewed by these factors.

A significant challenge in relating genetic data and historical records is the uncertainty of genetic age estimates. This arises partly from inherent variance in the underlying statistical distribution of posterior segment age given segment length (particularly for short IBD sharing), and partly from methodological uncertainty, such as the accuracy to estimate IBD segment length, which is limited by the density of SNP markers. Additionally, there is the fact that our posterior estimates do not account for complex demographic histories in the populations involved. By way of illustration, we have explored the impact of different demographic scenarios in the Supplementary Note, showing that different assumptions can lead to both earlier and later estimated dates. A full model incorporating both time and migration history is complex and the subject of ongoing research. Additionally, individual variation in intergenerational time introduces further uncertainty. It is also unclear whether the average intergenerational time has changed over the centuries, as historical records show variation in average age at marriage (“What Age Did People Marry in the British Past?,” n.d.). These uncertainties limit our current ability to assign precise historical dates or generational resolution given segment lengths, with uncertainty increasing for earlier time estimates.

As discussed earlier, our analyses are also limited to separate the genetic legacy of Anglo-Saxon and Danish Viking migrations. In our studies, we have not used ancient DNA sources, partly due to their low and sparse representation, and partly because it is unclear how extensive IBD mapping at this scale could be biased by the phasing of ancient DNA with modern Northern and Western European panels. An interesting modelling question in the future is how to integrate substantial IBD data from modern populations with the sparser ancient DNA data that have specific known dates, and whether we could identify distinct genetic markers to distinguish the contribution of these migration events.

Inferring the migration direction or origin of any given IBD sharing pattern based on genetic data is another challenge we have not addressed. The pairwise IBD sharing data we generated in this study cannot resolve this directly. However, future research using genealogical tree reconstruction methods (Gunnarsson et al. 2024; Zhang et al. 2023; Kelleher et al. 2019; Speidel et al. 2025) might offer opportunities to estimate the direction of gene flow from the tree structure.

In this paper, we brought together two of the currently largest genetic cohorts, the UK Biobank, and the Copenhagen Hospital Biobank & Danish Blood Donor Study in Denmark. It is conceptually feasible, though both technically and legally complex, to imagine extending this approach across even more cohorts, such as FinnGen (Kurki et al. 2023), All of Us (All of Us Research Program Genomics Investigators 2024), the Estonian BioBank (Leitsalu et al. 2015), the China Kadoorie Biobank (Chen et al. 2011) and the BioBank Japan (Nagai et al. 2017). To achieve this, we would need further innovation on the computational methods to discover and store IBD regions as well as substantial computing resources. Alongside providing a baseline understanding of these cohorts for basic and medical research, this would provide an unprecedented view of all substantial components of genetic exchange between the studied populations, allowing a comprehensive map of genetic exchange from medieval times to the modern day. This would serve as a highly complementary resource to the documented histories, and most likely, as we show here, both support and challenge aspects of these histories. Indeed, it is feasible to consider generating a comprehensive map of all European migrations from the medieval era onward by performing similar analyses across a pan-European modern cohort, and similarly for other continents and cross-continental migrations. From our study, we estimate that an even sampling of around 0.1% of the population (i.e. median sample size/median population size) would provide a resolution at the NUTS3 level of regionality in Europe. In addition, methodological improvements in integrating ancient genomes with modern ones would not only provide more support for specific migration patterns but also probably deepen the time resolution. We call for more interdisciplinary research between geneticists and historians, and the resources to be able to unlock this natural historical record inherent in our DNA.

## Methods

### Data Collection and Processing

#### Genetics data in the UK Biobank

We used 488,376 participants in the UK Biobank (UKBB) with SNP array data available, which are genotyped using the UK Biobank Axiom Array as a sampling of the UK population. There are 783,966 genotype markers in total after lifting over to the GRCh38 genome build. We define individuals with British ancestry as those ones self-reported as “White British” (UKBB Data Field: 21000) and born in the UK (UKBB Data Field: 1647). In total, we have 410,266 individual samples with 354,847, 35,912 and 19,507 from England, Scotland and Wales, respectively. While we might have included individuals with recent family immigration history into the UK based on this definition, we assume the proportion is relatively small, given we have a very large sample size.

To map individuals’ birthplaces into geographic areas, we transformed the birthplace coordinates (UKBB Data field: 129 and 130) into geographical areas including UK nations, UK NUTS level 2 (Regions; 2016 version), UK NUTS level 3 (2016 version) and councils.

#### Genetics data in Danish biobanks

For the Danish population, we included 370,259 individuals from Denmark with genotype data available from previous studies conducted under the Copenhagen Hospital Biobank (CHB) (Sørensen et al. 2021) and the Danish Blood Donor Study (DBDS) (Hansen et al. 2019; Erikstrup et al. 2023) including *Genetics of healthy ageing and specific diseases among blood donors – a GWA study under the Danish Blood Donor Study* (NVK-1700407, P-2019-99), *Genetics of cardiovascular disease - a genome-wide association study on repository samples from Copenhagen Hospital Biobank* (NVK-1708829, P-2019-93), and *Genetics of pain and degenerative musculoskeletal diseases - a genome-wide association study on repository samples from Copenhagen Hospital Biobank* (NVK-1803812, P-2019-51). All the samples were genotyped using the Illumina Global Screening Array with a total of 627,546 genotype markers. Reusing these genetic datasets for demographic and genealogy studies does not require specific scientific ethical approval.

We retrieved individuals’ birthplaces through the Danish civil registration record, i.e. Det Centrale Personregister (CPR register). There are 316,512 and 985 individuals born in Denmark (“continental Denmark”) and the Faroe Islands respectively. As we were aware of the evolving discussion about the use of genetic data from Greenland during this analysis we chose not to include samples from Greenland in this study.

Since the Danish administrative data does not contain individual ethnicity information, we adopted genomic approaches and inferred individuals’ global ancestry proportions (Method - Global ancestry inference using ADMIXTURE below). We projected the best-fitted unsupervised clustering via ADMIXTURE trained on a reference panel representing diverse world populations across continents onto Danish samples to infer their ancestral proportions at continental levels. There are 93.1% (294,782 out of 316,512) and 94.6% (932 out of 985) individuals born in mainland Denmark and the Faroe Islands, respectively, inferred to have more than 90% estimated European ancestral component. Unless otherwise stated, throughout the manuscript, we refer to the inferred European samples as European ancestry Danes (“Danes”). These samples might include individuals with recent immigration history in the last few generations from other European countries, given that we have a large sample size and the resulting IBD sharing has mirrored the geography of Denmark, we think most of the samples have major family ancestry from Denmark.

We adopted the nomenclature of geographic divisions, including regions, provinces and municipalities, provided by Statistics Denmark (URL: https://www.dst.dk/en/Statistik/dokumentation/nomenklaturer/nuts).

To validate the observed population structure in Denmark, we also curated a subset of individuals who have both of their parents in the DNK cohorts and were born in the same province as both their parents. As the dataset is much smaller, we also took a rigorous approach to remove one individual in close relative pairs (up to third-degree) to avoid family-specific patterns. This results in a refined subset of 28,135 individuals.

#### Danish Multi-Generation Family Registry

For the Danish Multi-Generation Family registry (MGR-lite registry), there are a total of 7,771,981 individuals recorded. MGR-Lite is a registry that documents the complete parental relationship for children (i.e. legal guardian) born in 1953 or later and still alive in 1968 to their parents based on both the CPR register and family relationships recorded in church books (https://www.registerforskning.dk/projekter/mgr-lite/) and partially includes parental relationships for children born before 1953. Additionally, we excluded relationships where parental ages were unreasonable or where the relationship was recorded as an adoption built on our previous work (Westergaard et al. 2024). In data preprocessing, we removed 866,638 individuals with no first-degree relatives’ records (who have neither parents nor children). We also removed five individuals recorded with multiple pairs of parents. In total, 6,845,337 individuals are analysed.

Given that the raw record in the MGR-lite registry only includes the parent-child relationship, we infer other familial relationships according to the following steps: (1) extract ancestor-offspring relationships. We use an adjacency matrix to record whether individual *i* is an ancestor to individual *j* and the value of the element indicates the number of separating meioses; (2) extract cousin relationships; To reduce the amount of computation, we only extracted the cousin relationships for individuals with both familial record and genetic data available. We iterate over each individual as a potential ancestor extracted from step (1) and construct the cousinship among all his/her descendants. For each pedigree relative, we infer the number of separating meioses. In total, we extracted 56,329 pairs of ancestor-offspring relatives and 54,089 pairs of cousins (Table S2 and Table S3), which we are going to use to validate the genetic data.

We are limited to extract pedigree relationships for relatives of distant degrees larger than five due to the incomplete record of parental relationships for individuals born before 1953 thus the relatives sharing ancestors born before 1953 are only partially identified in our familial registry data.

Note that we only assume individuals are related through a single genealogical path, given that we are assessing close relatives. While it’s likely that individuals can be related through multiple genealogical paths, especially for individuals relatively distantly related, we are unable to identify them comprehensively given our limited information about distant pedigree relatives in the data; thus, we ignore this scenario in our analysis.

### Genotype data processing

To process the UKBB SNP array data, we conducted sample-level quality control (QC) and SNP-level quality control. For sample-level quality control, we removed individual outliers due to poor genotyping quality based on heterozygosity and missing rates based on standard QC output provided by UKBB (resource 531 URL: https://biobank.ndph.ox.ac.uk/ukb/refer.cgi?id=531). For the SNP array data in Danish biobanks, we conducted the SNP-level QC through the following steps: (1)Filter SNPs with >0.1 missing, (2)remove duplicate SNPs.

### Haplotype Phasing

We used Eagle v2.4.1 (https://alkesgroup.broadinstitute.org/Eagle/) (Loh et al. 2016) to estimate the haplotype phases. The genetic map was downloaded from HapMap (ftp://ftp.ncbi.nlm.nih.gov/hapmap//recombination/2011-01_phaseII_B37/genetic_map_HapMapII_GRCh37.tar.gz).

### Estimating IBD sharing

We called IBD using FastSMC(Nait Saada et al. 2020) to detect IBD shared within the past 50 generations with a minimum IBD segment length of 1cM using parameters “--time 50 – min_m 1.0”. We conducted independent IBD calling based on three different SNP marker sets: (1) within the UK, using all SNP markers from UKBB; (2) within Denmark, using all SNP markers from Danish samples (DNK); (3) between the UK and Denmark, using shared SNP markers from the combined UKBB and Danish datasets (UKBB+DNK). All computations were performed within the secure computing infrastructure of the Danish National Genome Center and EMBL’s European Bioinformatics Institute.

While imputation could potentially improve the IBD mapping between the UK and Danish samples, we considered and then rejected adopting this approach. On one hand, since British and Danish populations are two closely related populations (Hudson Fst = 0.00075), we were concerned about the biases introduced, for example, the need to use predominantly Northern and Western European, with substantial British and Scandinavian haplotypes included in the imputation panels. On the other hand, our sensitivity analyses of intra-Danish comparisons confirm that using intersecting markers, despite underestimating IBD segment counts, show distributions highly aligned with the full marker set (Spearman correlation coefficient *rho* is between 0.99 and 1) and have no qualitative differences (Figures S3-S4), supporting their use in downstream analyses.

We computed the joint IBD calling between the UKBB and DNK (UKBB+DNK) by using the following procedures:

1. Phase each biobank independently
2. Lift over the UKB phased genotype data from Human reference genome build GRCh37 to GRCh38. We discarded the positions that could not be lifted over.
3. Extract the subsets of SNP markers in both biobanks
4. Use the above subsets to call IBD

We conducted additional filtering after IBD calling to reduce the false positives caused by low SNP density in the IBD calling in the joint SNP data. We removed IBDs across areas with SNP density lower than 50 SNPs/cM, which would likely lower the true positive rates but increase the positive predictive value.

In the downstream analysis, we focus on IBD segments larger than 2cM given that the IBD 1-2cM shows lower sharing rate compared to 2-3cM using the DNK IBD call set (Figure S5), reflecting the reduced sensitivity of IBD estimation at 1-2cM. Given we have lower SNP density in the joint IBD calling, IBD 1-2cM might be underestimated to a stronger degree.

To summarise the level of IBD sharing among pairs of individuals at the population level, we calculate the total IBD sharing (>2cM) and number of IBD segments (>2cM) by summing over all the identity-by-descent segments across autosomes between the two individuals. To handle the computing efficiently, we developed bespoke analysis code in C based on a previously published software IBDkin (Zhou, Browning, and Browning 2020). Unless specified, we removed pairs of close relatives up to third-degree relatedness (equivalent to first cousins, great-grandparents andgreat-grandchildren) using the lower bound of the expected total IBD sharing threshold of 634cM for analysing the total IBD sharing(Manichaikul et al. 2010). This shall reduce the impact of ascertainment bias due to close genetic relatedness.

Similarly, the total length of ROHs and a number of ROH segments were calculated per individual by summing the total IBD sharing and the number of IBD segments between two chromosome copies across autosomes. We first evaluated the ROHs based on all the available samples. Given that we want to reduce the effects of family-specific patterns especially in communities of small size (autozygosity is correlated between siblings though the correlation is highly variable(Kirin et al. 2010; Thompson 2013)), we also took a stringent manner to only include 160,957 unrelated samples in the DNK cohorts after randomly removing one individual in each pair of close relatives (up to third-degree relatedness). We find the results are highly comparable in the two evaluations (Figure S10), suggesting the high ROHs are likely driven by a small effective population size instead of specific family patterns.

To study IBD sharing for a specific length, IBD segments were grouped into bins of different lengths defined as half-open intervals, where each bin includes the lower bound but excludes the upper bound, denoted as [a,b). For example, an IBD bin between 2-4cM indicates all IBD segments *x* such that *2* ≤ *x* < *4* cM.

### Validation of pedigree relationship

In the validation, we compared the mean total IBD sharing observed from IBD estimation with the expected given the number of separating meioses inferred from the pedigree relationship in the MGR-lite. The results are shown in Figure 2. The expectation of IBD sharing is computed based on Huff et.al 2011(Huff et al. 2011). We find that the distribution of observed total IBD sharing given the number of separating meiosis largely follows the theoretical expectation. For parent-child and grandparent-grandchild relatives, we also observe long tails on the lower end, suggesting that there could be unbiological related pairs or possibly unrecorded adoptions by relatives. Consequently, we also observe that the respective distributions among pairs of shared ancestors are skewed towards the lower end. For pairs of shared ancestors, a second mode with less density is also observed, indicating that there are pairs sharing only one ancestor, e.g. half-siblings or half-cousins.

We also compared the number of IBD segments observed with the expected. The results are shown in Figure S2. For ancestor-offspring pedigree relatives and cousins up to first-degree cousins, we find that the observed number of IBD segments is higher than expected. Given that the total genomic sharing observed for these relatives aligns with expectations, we believe this is likely explained by statistical phasing error. The switch error of the statistical phasing method we applied to the Danish cohorts is likely similar to that in UKBB, occurring at a rate of approximately one error per 20 cM (Loh, Palamara, and Price 2016; Naseri et al. 2019). Thus, we expect phasing errors to more significantly affect close relatives (those less than third-degree) since they are expected to share long IBD segments (≥20 cM). Given this threshold for phasing error, we consider it less of a concern for IBD sharing estimation in distant relatives, which we used in the migration analysis.

### Estimate the effective population size

We applied the IBDNe(Browning and Browning 2015) to estimate the trajectories of effective population size over time. The following options were used: “filtersamples=TRUE gmax=50 mincm=2” to remove close relatives and only use the minimal IBD segment length at 2cM up to 50 generations ago. 95% confidence intervals of Ne were estimated by using the default options.

### Dimension reduction of pairwise IBD sharing via multidimensional scaling

We use the pairwise matrix of mean total IBD sharing (cM) as a similarity matrix *S*. We then convert *S* into a distance matrix *D* by subtracting each element of *S* from its maximum element value, i.e.*D_i,j_* = *max*(*S*) − *S_i,j_*. Multidimensional scaling was then computed with the input distance matrix *D* using the function “cmdscale” in the “stats” R library.

### Global ancestry inference using ADMIXTURE

We applied unsupervised clustering implemented in ADMIXTURE(Alexander, Novembre, and Lange 2009) to infer the continental ancestry proportions of individuals in the Danish cohorts. To train the clustering model, we used the pre-curated harmonised dataset (1kGP+HGDP) with 3,262 individuals(Koenig et al. 2024), which combines both the 1000 Genomes Project (1kGP) and Human Genome Diversity Project (HGDP) as training data. It includes African (AFR), Admixed American (AMR), European (EUR), Middle Eastern (MID), CSA (Central/South Asia), and East Asian (EAS). In the training, the genotype data for each sample individual is modelled as a mixture of *K* ancestral components. We first removed the SNP markers with strong LD in the DNK cohorts by applying PLINK with option “--indep-pairwise 50 10 0.5” (pruning variants within 50kb with a squared correlation greater than 0.5). We used the subset of autosomal genotype markers (n=385,272) from the 1kGP+HGDP dataset that overlap with the LD-pruned DNK cohorts. To estimate the best number of ancestral components *K* in the data, we applied a cross-validation to compare the predictive accuracy on hold-out test data using a range of values from *K*=2 to *K*=8. *K*=5 is chosen given that the slope of the cross-validation error curve is not changing drastically with the increasing *K* (Figure S12). Each ancestral component is annotated based on its proportions in each continental sample. The ancestral proportions of the Danish samples were predicted by projecting them onto the learned allele frequencies from the training samples.

### Identify populations of other major European ancestry in the UKBB+DNK dataset

To investigate the influence of other European countries on Danish and British ancestry, we assembled a reference panel for European populations using the combined samples (UKBB+DNK) from the UK Biobank (UKBB) and Danish cohorts (DNK). Here, we outline the steps involved in our methods:

#### Sample Inclusion

To include samples with non-British and non-Danish European ancestry in the UKBB, we selected samples born in Europe (excluding the UK and Denmark) (UKBB Data Field: 20115) who self-reported their ethnicity as “White” or “Any other white background” (UKBB Data Field: 21000). For individuals with Irish ancestry, we specifically selected those born in Ireland and self-reported with “Irish” ethnicity. In the Danish cohorts, we included individuals with predominantly predicted European (>90%) ancestry who were born outside Denmark and the UK. We exclude any country with fewer than 100 samples to avoid the bias of small sample sizes, which might have excluded samples from some East European and Southern European countries due to their low ascertainment in our datasets.

To make sure the clustering results would not be driven by the fine-structure of populations of British and Danes, given their large sample size in total, we downsampled these groups. For the British, we randomly selected 9,000 British individuals from the UKBB, with 3,000 from each of England, Scotland, and Wales. For the Danish cohorts, we also randomly selected 3,000 individuals from a subset where both the child and parents were born in the same municipality.

In total, we included 34,555 individuals with 22,555 ones born in 26 European countries (excluding the UK and Denmark) and 12,000 ones born in the UK and Denmark to construct a reference panel of European populations.

#### Community detection using individuals’ pairwise IBD sharing network

Given that we only have information on individuals’ countries of birth, it’s important to note that these may not fully indicate their ancestral backgrounds. They could be British or Danes born overseas. To identify other European populations, we assume individuals in the same population would highly share their ancestry compared with those in other populations. To identify groups of individuals with shared ancestry, we used an unsupervised community detection approach, assuming that individuals with similar ancestry background would be closely clustered in the IBD-sharing network. Thus, the resulting detected clusters could allow us to identify different populations.

We constructed an IBD-sharing network based on pairwise total IBD sharing among the selected 35K samples. To reduce the impact of recent migration, we applied a more stringent criterion by removing pairs with total IBD sharing greater than 56 cM (the expected total sharing of third-degree cousins), thus removing relatives up to third-degree cousins.

To identify population clusters, we applied the Leiden community detection algorithm(Traag, Waltman, and van Eck 2019), which optimises the modularity to partition the network into distinct clusters. We adopted a multi-level hierarchical approach. We detect sub-clusters from the clusters of one level up by varying the resolution parameter from 1 to 2, increasing by 0.1 at each step until sub-clusters are identified.

Using four levels of clustering, we identified 45 clusters in total, with 30 mainly composed of individuals with non-Danish or non-British ancestry. The clustering results captured broadly the ancestry diversity within our samples at the country level (Figure S20). We annotated each cluster based on the proportions of members’ country of birth. Thus, individuals in the same cluster were considered to share a major ancestral background from their country of birth. We also made sure there are comparable proportions of samples from both the UKBB and DNK cohorts to form the same population cluster. For example, German clusters were identified only if German-born samples from both the UKBB and DNK cohorts clustered together. We only considered the clusters with a sample size larger than 200 to ensure well-supported results. We did not identify further sub-clusters within each country as it’s outside the scope of our study, and we don’t have detailed birthplace data to validate.

Reassuringly, the majority of Danish samples (97.1%) and British samples (91.2%) are grouped within their clusters. The community detection approach also helps us to remove individuals born in one country but placed in a different cluster, suggesting they are likely to have mixed ancestry or recent immigrant history.

As our clustering results depend on both the sample size and underlying ancestry diversity in our samples, the clusters should be interpreted as groups of individuals with shared ancestry rather than distinct populations.

The resulting clusters are used to identify proxy populations with major ancestry from the neighbouring countries of Britain/Denmark used in Figure 4 and Figure 6. The sample size for each proxy population is shown in Table S5.

### Estimate the Age of Most Recent Common Ancestors (TMRCA) using IBD segment length

Given two individuals with a shared IBD of length *l*, we are interested in inferring the time to their most recent common ancestor *g* (TMRCA) or segment age. In the manuscript, we applied the coalescent theory and Bayesian approach to estimate the posterior distribution of TMRCA *g* given an observed segment length.

#### Maximum a posteriori (MAP)

We first derived the posterior probability of *g* given a random segment of length *l* following previous literature(Palamara 2014; Fournier et al. 2023).The posterior probability of *g* given *l* taking into account the prior distribution *f*(*g*), which defines the coalescent probability between individuals and further depends on the underlying demographic history, the likelihood function *f*(*l*|*g*) which describes the distribution of IBD lengths if TMRCA is known, and the marginal probability of observing a segment of length *l:*

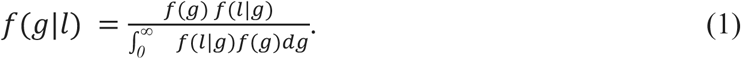

Under a simple demographic model with constant population size *N* (number of haploids), the coalescent probability *f*(*g*) at a random site between two haploids follows an exponential distribution with parameter 1/*N* (or geometrical distribution for time measured in discrete units),

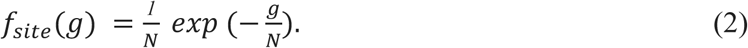

Given that the occurrence of recombination breakpoints along the genome follows a Poisson process (ignoring crossover interference and chromosome ends) under the Haldane’s model(Haldane 1919), the genetic distance from a random site to the recombination breakpoint over *g* meioses separating an offspring and its ancestors (i.e. segment length inherited from ancestors g generations ago), either to the left or right follows an exponential distribution with parameter *g* (the number of breakpoints per Morgan). For two haploids sharing the same ancestor from *g* generations ago (separated by *2g* meioses), the distance from a chosen site to the closest recombination breakpoints from the left or right corresponds to an exponential distribution with rate *2g (Palamara 2014; Temple and Thompson 2024).* Thus, the length of shared IBD segments overlapping the chosen site, which is the sum of two independent exponential variables (one from each side), follows an Erlang-2 distribution with rate parameter *2g*:

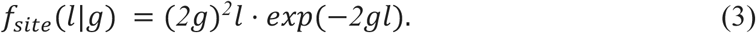

Thus, the posterior probability *f*(*g*|*l*) could be derived by substituting Equations 2 and 3 into Equation 1:

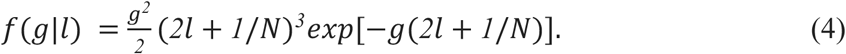

Note that the above derivation of the posterior distribution is conditional on a random site, but it’s equivalent to the distribution derived using a random segment. For a random segment, following previous literature considering the IBD process under the SMC model, the prior distribution of segment age *g* is distributed as (note that *g* here in unscaled generation time) (Carmi et al. 2014)(Li and Durbin 2011):

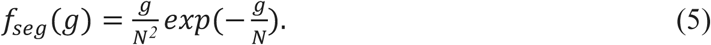

Given a TMRCA=*2g* separating two extant haploids, their shared segment length *l* follows an exponential distribution with parameter *2g*. Thus, we can derive the same posterior density as Expression 4.

We visualised the above posterior density distribution *f*(*g*|*l*) for different IBD lengths in Figure S45. For a given IBD length, the possible age of the segment spans a wide range. Shorter IBD segments are associated with a wider range of possible ages, and longer segments with less. We use the age with the highest posterior probability for a given IBD length as the most probable generation time, i.e., the Maximum a Posteriori (MAP) estimate 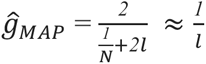 (the human populations in our studies have a large *N* with the scale of *10^5^* − *10^8^* thus 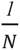 could be ignored).

To quantify the uncertainty of the posterior age, we calculated the 50% highest density interval (HDI), which is the most credible age range that covers 50% of the posterior distribution, i.e.,*P*(*g_1_* < *g* < *g_2_*| *l*)=50%. Age values within the 50% HDI are more credible (i.e., have higher probability density) than values outside the HDI, and the values inside the HDI have a total probability of 50%. We derived the HDIs numerically by grid approximation following the implementation of Kruschke (John 2015). The calculated MAP age and 50% HDI for each IBD length are displayed in Table S6.

In the manuscript, for IBD sharing with a length bin, we define the bin by grouping lengths with similar geographic distribution patterns of IBD while also maximising the differences between different distinct bins. Accordingly, for each length bin, we define the MAP age range by taking the minimum and maximum MAP ages across the lengths in the bin. Similarly, we define the combined 50% HDI for the bin by taking the minimum and maximum of the 50% HDI across those lengths.

Since the prior depends on underlying demographic scenarios, we evaluated the impact of complex demographic histories on the posterior distribution of segment age with ancestry simulations as detailed in the Supplementary Note. Different scenarios, such as exponential growth, population bottleneck and mass migrations, can shift both the mode and the credible interval of the posterior distribution.

#### Converting Generations to Calendar Time

We assume a generation time of 28 years (Moorjani et al. 2016). For UKBB samples, the median year of birth is 1950 CE. For Danish samples, the median year of birth is 1960 CE. Across the two cohorts, we use the median birth year of 1955. We use this point in time as the reference for the “present” relative to our samples.

In summary, there are various sources of uncertainty in our estimate of TMRCA: (1) the population demographic history; (2) statistical property of the posterior distribution, i.e. its credible interval is quite wide; (3)the estimation of IBD segment endpoints, which depends on the density of SNP markers and thus affects the accuracy of IBD length estimates; and (4) uncertainty in the generation time, with Moorjani et.al (Moorjani et al. 2016) estimating the mean generation interval to be between 26-30 years based on recombination clock, and Wang et.al estimating 24-30 years based on mutation clock (Wang et al. 2023; Moorjani et al. 2016).

### Statistical test of spatial autocorrelation

We used the Moran test to assess the spatial association of IBD sharing. Global Moran’ test is first conducted to assess whether the IBD sharing is randomly distributed. The local Moran’s test is used to assess whether the degree of IBD sharing of a given area is associated with its neighbouring area. We define neighbours as the five areas with the shortest geographic distance to the centroid of a given area. For IBD sharing between Britain and Denmark, the local Moran’s test was conducted for each area, and the results are categorised as the following groups (shown in Figure S24-S25):

Significant spatial associations between the tested area and its neighbours are identified with a *P*-value threshold of 0.05, which include High-High(hotspots, high IBD sharing with high IBD-sharing neighbours), Low-Low (coldspots, low IBD sharing with low IBD-sharing neighbours), High-Low (outliers, high IBD sharing with low IBD-sharing neighbours) and Low-High (outliers, low IBD sharing with high IBD-sharing neighbours); No significant spatial association (annotated as “>0.05”) or missing data.

## Code Availability

The programs to compute the IBD sharing statistics are available at https://github.com/XiaoleiZ/UK_DNK_IBD.

## Ethical Approval

The UK Biobank has received approval from the National Information Governance Board for Health and Social Care and the National Health Service North West Centre for Research Ethics Committee (Ref: 11/NW/0382). This research was conducted using the UK Biobank Resource under project 49978.

All investigations were conducted in accordance with the tenets of the Declaration of Helsinki. The genotype datasets from the Copenhagen Hospital Biobank (CHB) and Danish Blood Donor Study (DBDS) were collected from previous studies (NVK-1700407, P-2019-99; NVK-1708829, P-2019-93; NVK-1803812, P-2019-51). Ethical approval was not required for reusing these datasets in demographic and genealogical research. The use of data from the MultiGeneration Registry-Lite was approved by the Danish National Archives (Case number 22/33918; Reference number KAMB-MOEL). Informed consent or evaluation by a research ethics committee is not required for registry-based research in Denmark.

## Contributions

S.B. and E.B. conceived the study. X.Z., S.B., and E.B. designed and developed the study. X.Z. performed the data analysis. X.Z., P.H., A.S., S.B. E.B. interpreted the data with the other co-authors’ inputs. P.H. contributed to analyzing historical evidence. X.Z., P.H. and E.B. drafted the manuscript with input from all co-authors.

## Acknowledgements

We thank all the volunteers for their participation in our study cohorts, including the UK Biobank, the Copenhagen Hospital Biobank and the Danish Blood Donor Study. This study has also received valuable suggestions and support from the members of the Birney and Brunak groups, particularly Kumar Gaurav, Maria Herrero-Zazo, Troels Siggaard and Lisa Cantwell. The authors also thank the experimental analyses conducted by Casper Siu.

This study was supported by the Novo Nordisk Foundation (grants NNF170C0027594, NNF14CC0001). XZ has also been funded by the Borysiewicz Interdisciplinary Fellowship from the University of Cambridge during the study.

The funding bodies had no role in the design and conduct of the study.

