## Supplementary Information for "Population-scale inheritance analysis of 858,635 individuals reveals North Sea migration from the Middle Ages to the Industrial Revolution"

### Supplementary Figures

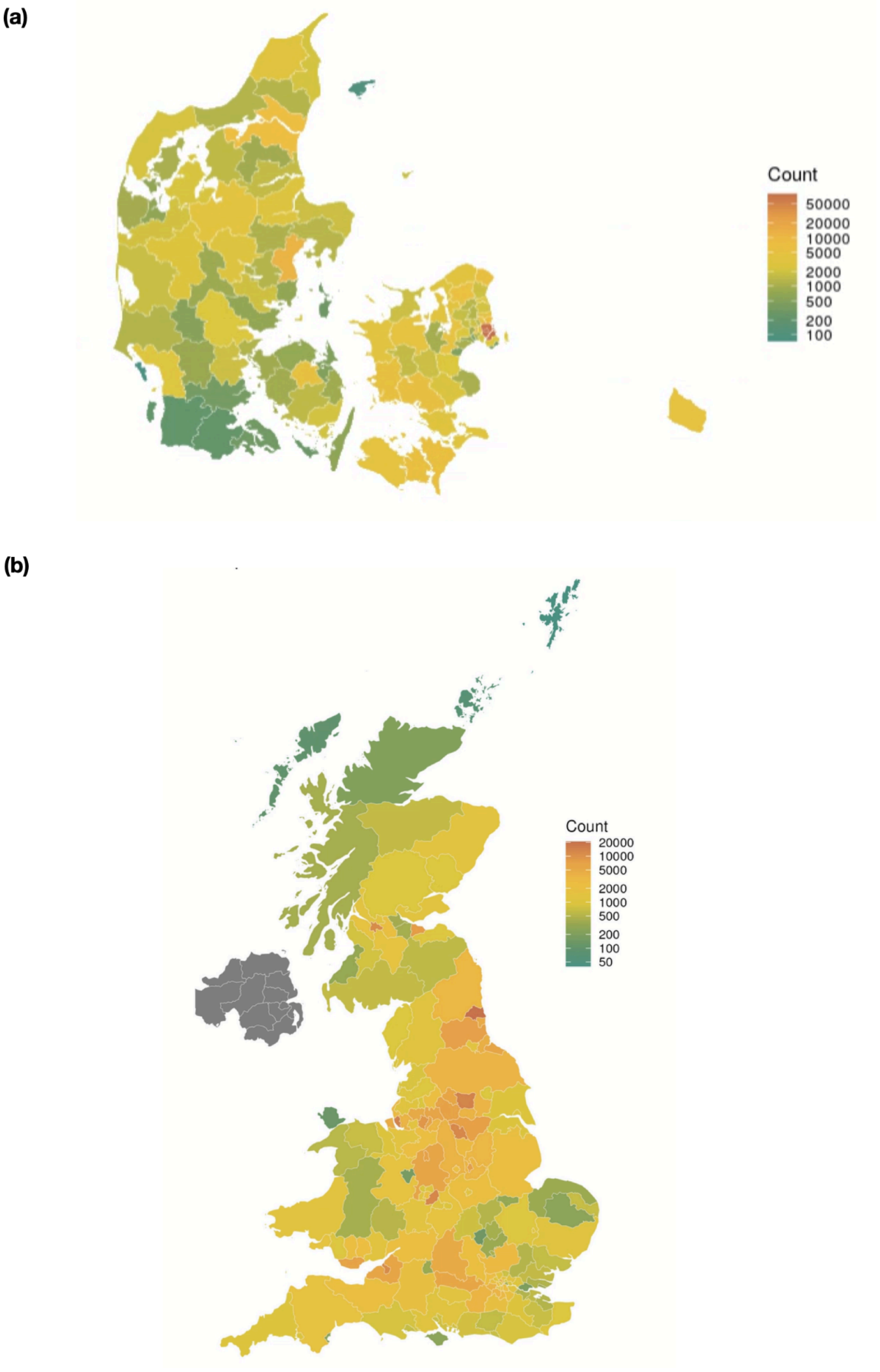


**Figure S1. Geographic distribution of samples' birthplace and their sample sizes at the finest scale considered in the study.** For Denmark, we use the municipality level (in total 98 municipalities), with sample sizes ranging from 73 to 79,977. For Britain, we use the NUTS level 3, with sample sizes ranging from 40 to 20,781 (in total 168 subdivisions).

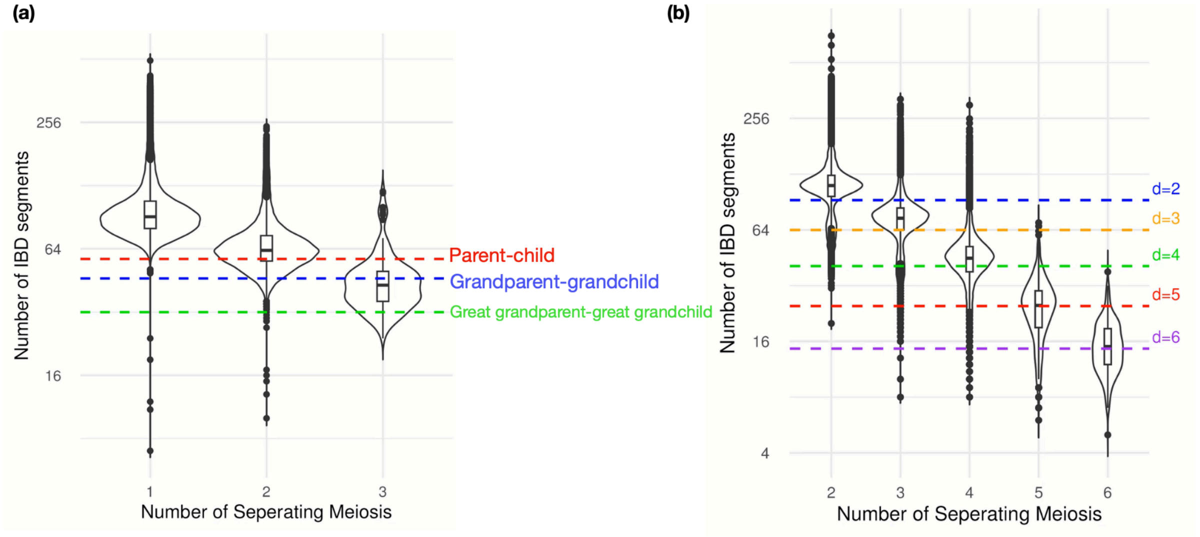


**Figure S2. Comparison of the number of IBD segments observed and expected, given the inferred pedigree relationship** (measured as the number of separating meioses). (a) among individuals with a known number of meioses separating them from a direct pedigree ancestor (b) among relatives sharing a common pedigree ancestor, excluding direct ancestor-offspring relationship. We discussed the discrepancy between the observed and expected number of IBD segments in the Methods. Note that IBDs are compared between haplotypes here instead of between individuals.

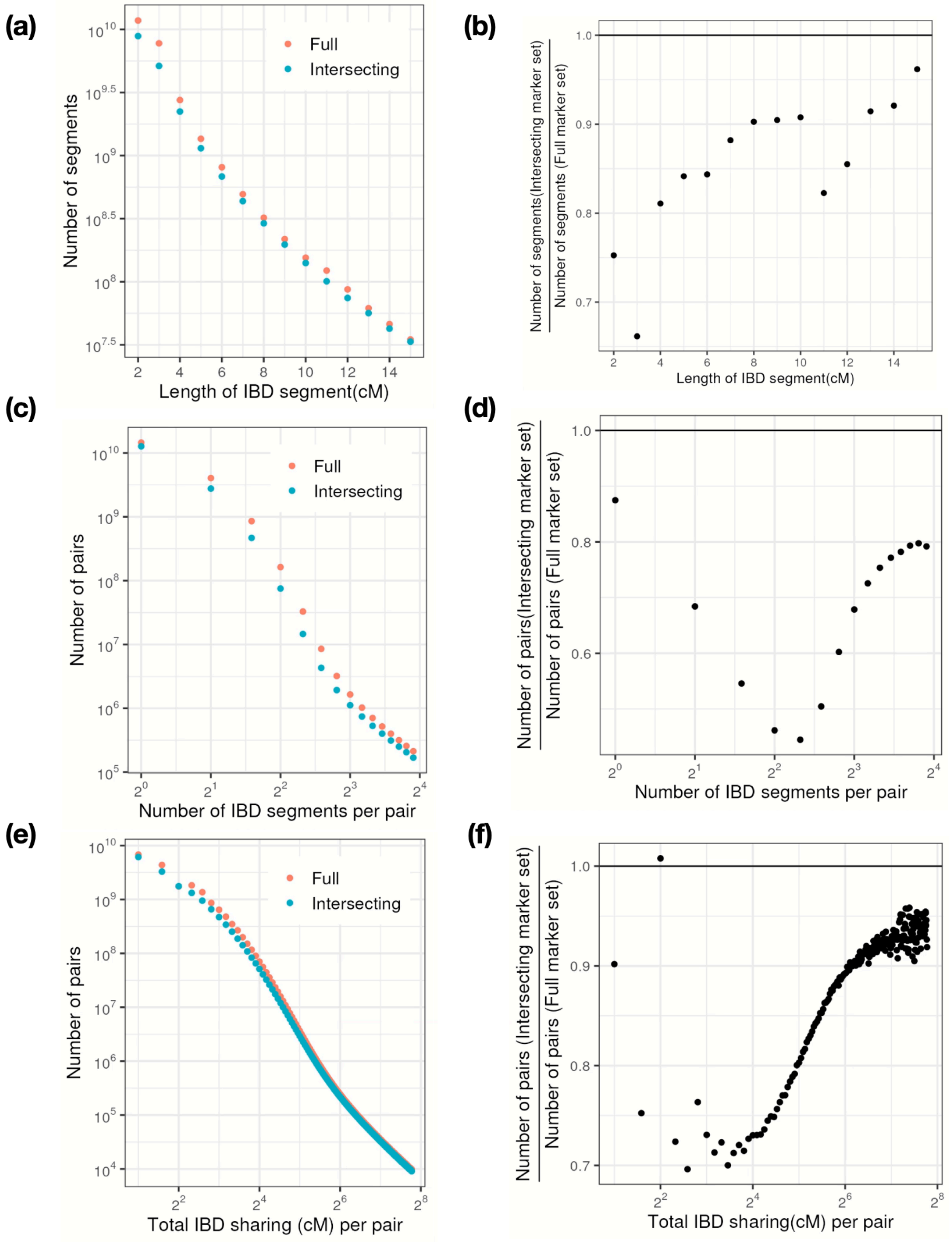


**Figure S3. Compare IBD estimation using intersecting versus full marker sets in the Danish cohorts. (a)**Number of inferred IBD segments (Y-axis) across different IBD length bins (X-axis) for intersecting and the full marker set (Spearman correlation *rho* = 1 across bin counts). **(b)** The ratio of the number of IBD segments detected by intersecting markers to those detected by the full marker set across length bins. Intersecting markers recover 66–96% of IBD segments compared to the full set, with shorter IBD disproportionately underestimated. **(c)** Distribution of the number of IBD segments per pair (X-axis) shared across pairs (Y-axis: number of pairs) for intersecting and the full marker set (Spearman correlation *rho* = 0.99), and the corresponding ratio shown in **(d)**. **(e)** Distribution of total IBD sharing (X-axis) shared across pairs (Y-axis: number of pairs) for intersecting and the full marker set (Spearman correlation *rho* = 1), and the corresponding ratio shown in **(f)**. Overall, intersecting markers identify 82% of pairs with IBD segments compared to the full set, with distributions of IBD sharing strongly correlated.

**(a)**

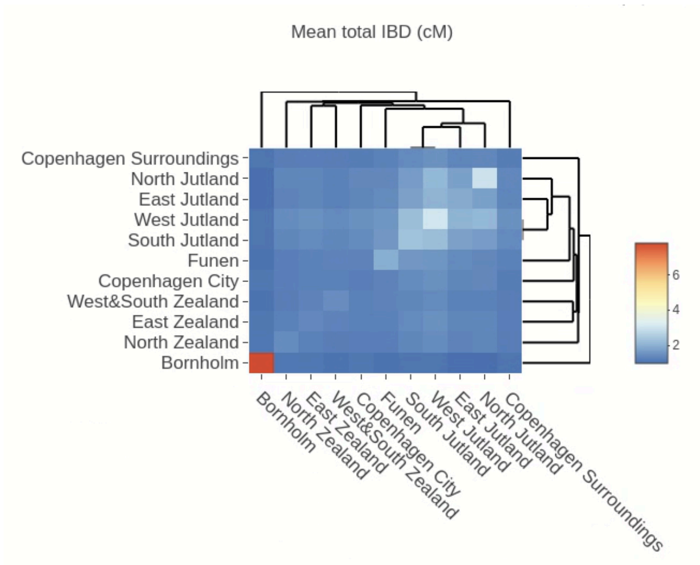


**(b)**

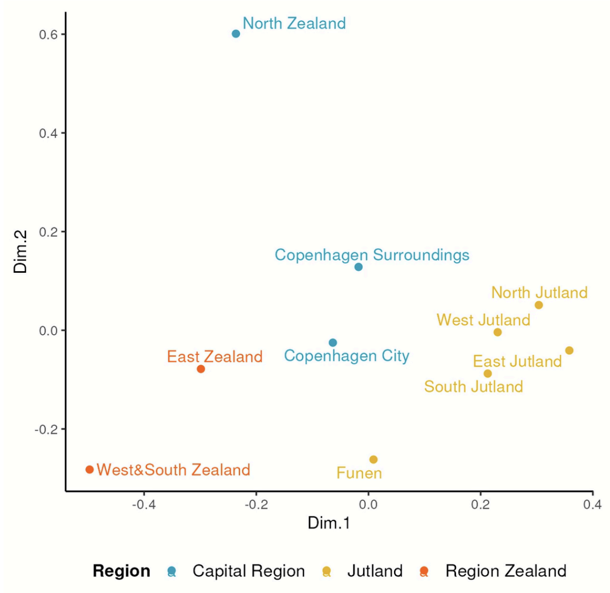


**Figure S4. Qualitative assessment of IBD estimation using intersecting markers in the Danish cohorts.** We analyzed the population structure within the Danish cohorts using IBD calling based on intersecting markers: **(a)** the mean total IBD sharing (cM per pair) across regions of Denmark, and **(b)** the Multi-dimensional scaling of the dissimilarity between regions of Denmark (except Bornholm), derived from pairwise mean total IBD sharing. Both plots closely resemble the results generated using the full SNP marker set, as shown in Figure 3(a) and Figure 3(b), respectively.

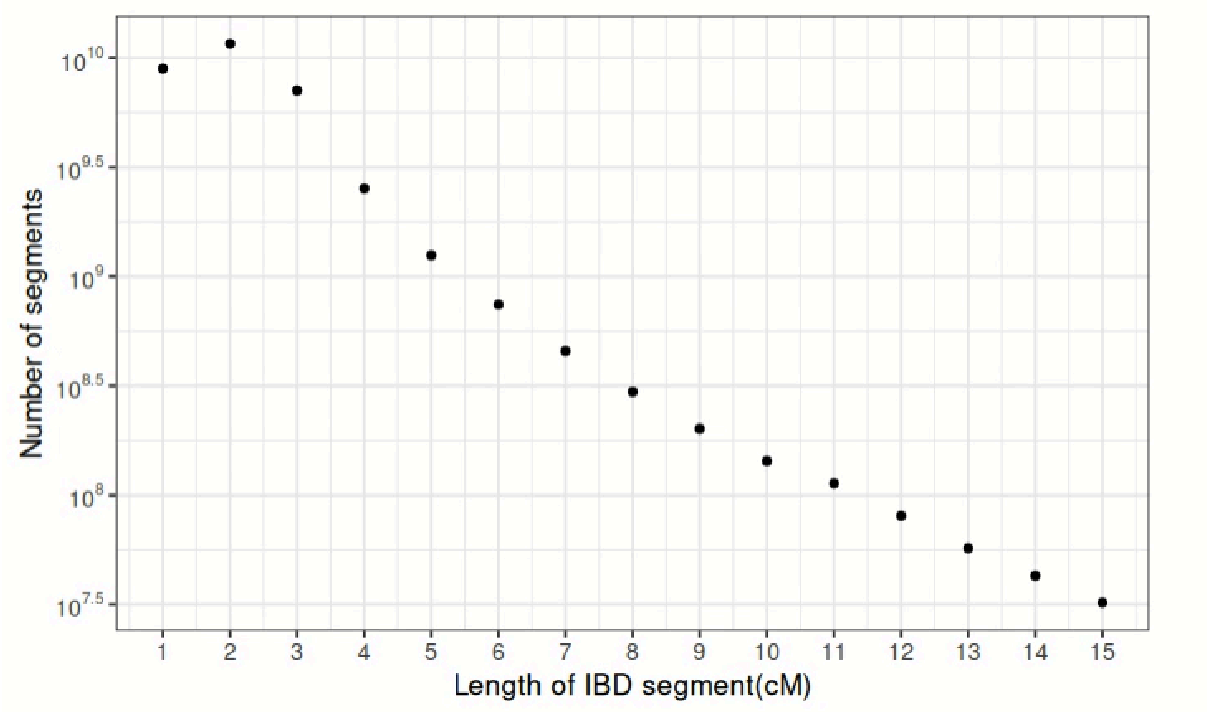


**Figure S5. Number of inferred IBD segments (Y-axis) across different IBD length bins (X-axis) for the DNK full marker set.**

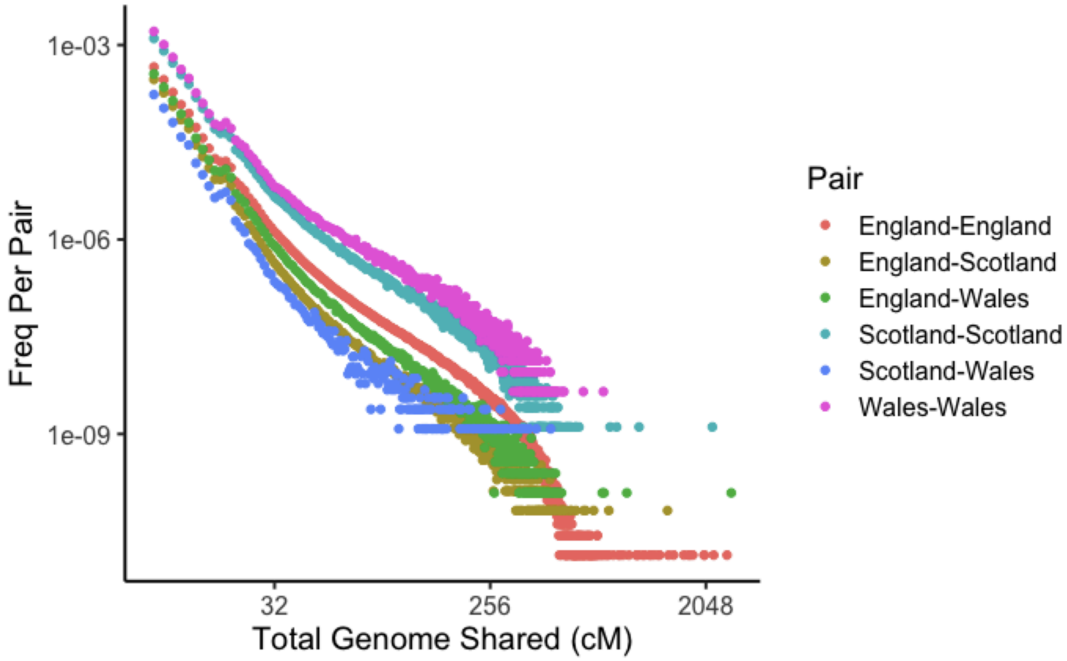


**Figure S6. Distribution of total IBD sharing across the three British subpopulations (English, Welsh and Scottish) in the UKBB.** The full UKBB SNP markers are used to estimate the IBD sharing.

**(a)**

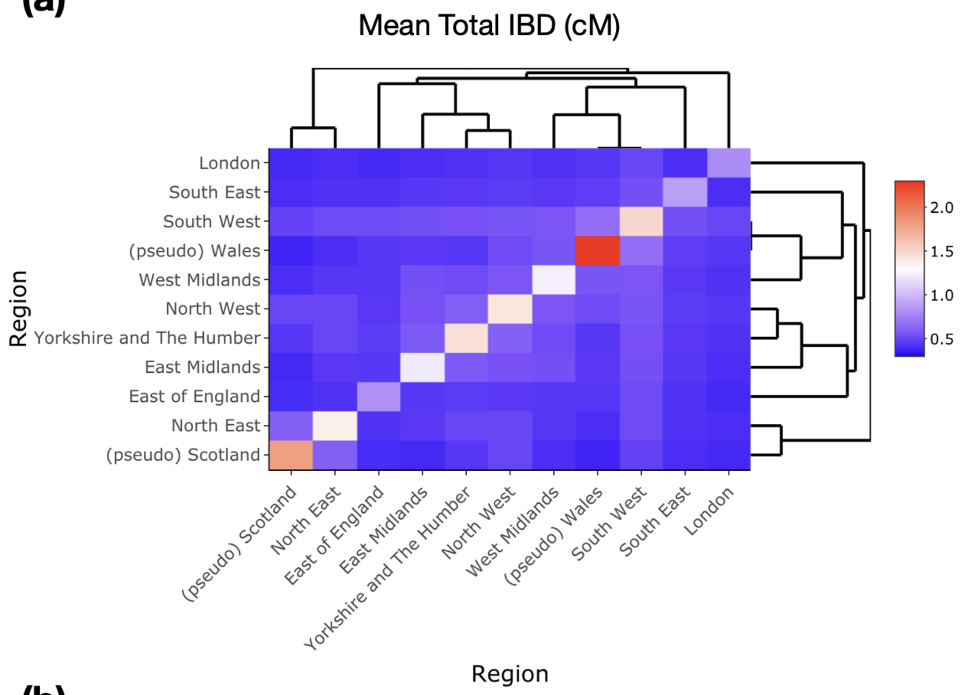


**(b)**

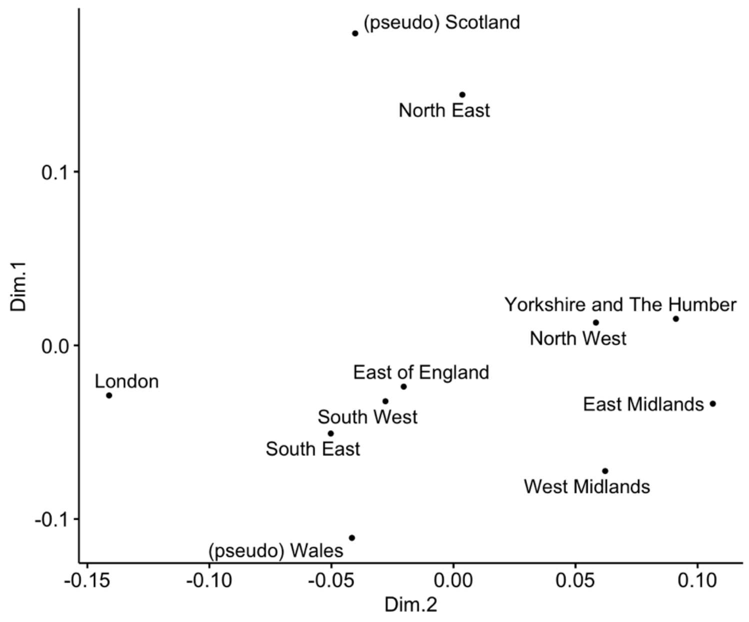


**Figure S7. The total IBD sharing at the level of Britain's regions reflects the geography (except London). (a)** The pairwise total IBD sharing across areas of Britain and **(b)** Multidimensional scaling on the dissimilarity between regions of Britain based on their pairwise total IBD sharing.

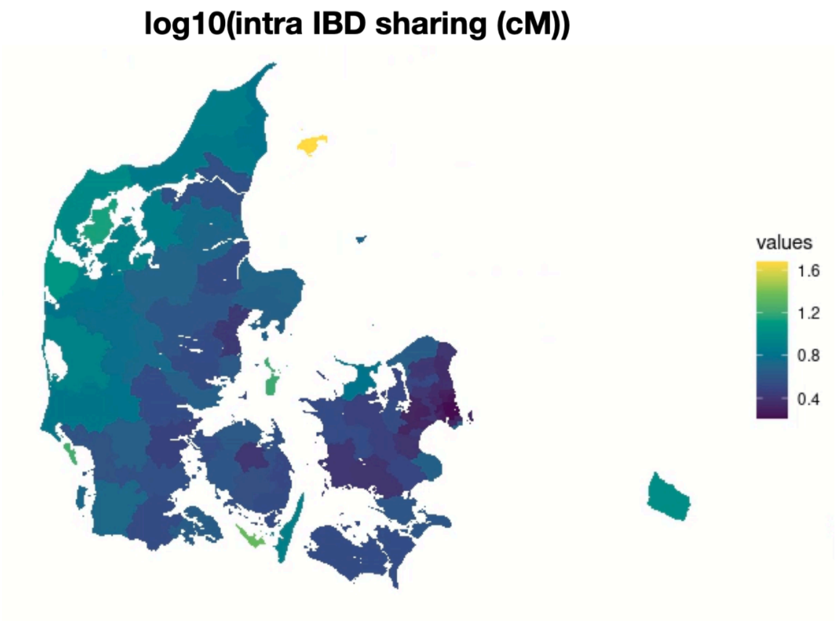


**Figure S8. The geographic distribution of intra-population IBD sharing across Danish municipalities.**

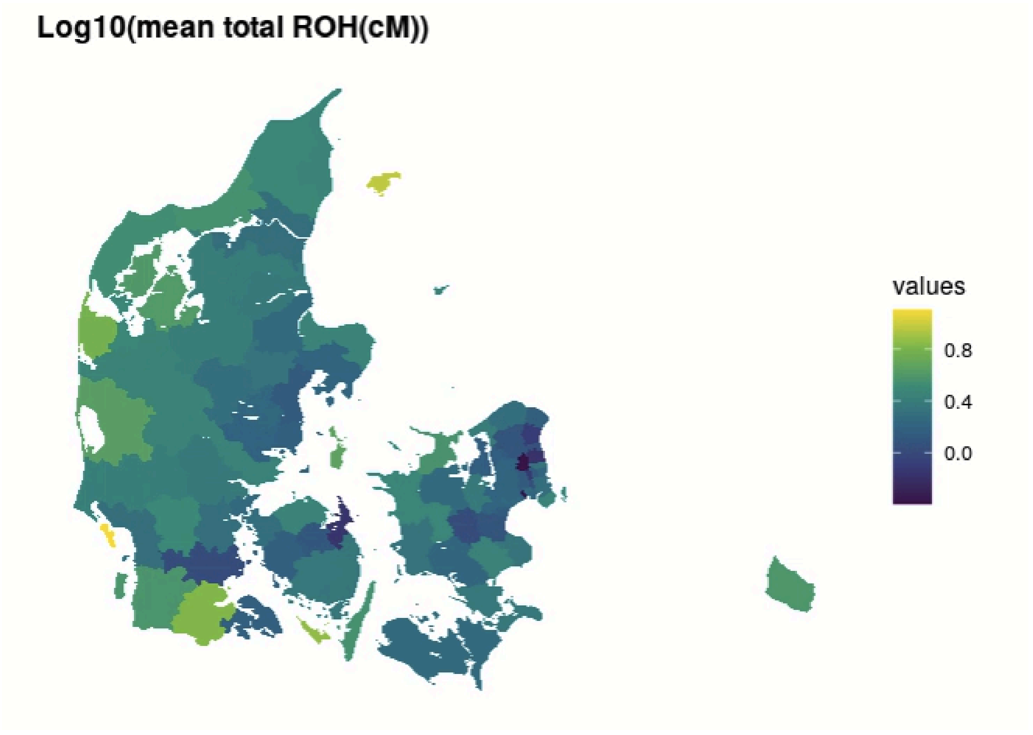


**Figure S9. The geographic distribution of the mean total length of ROHs within Danish municipalities based on Danes with a major European ancestry proportion.**

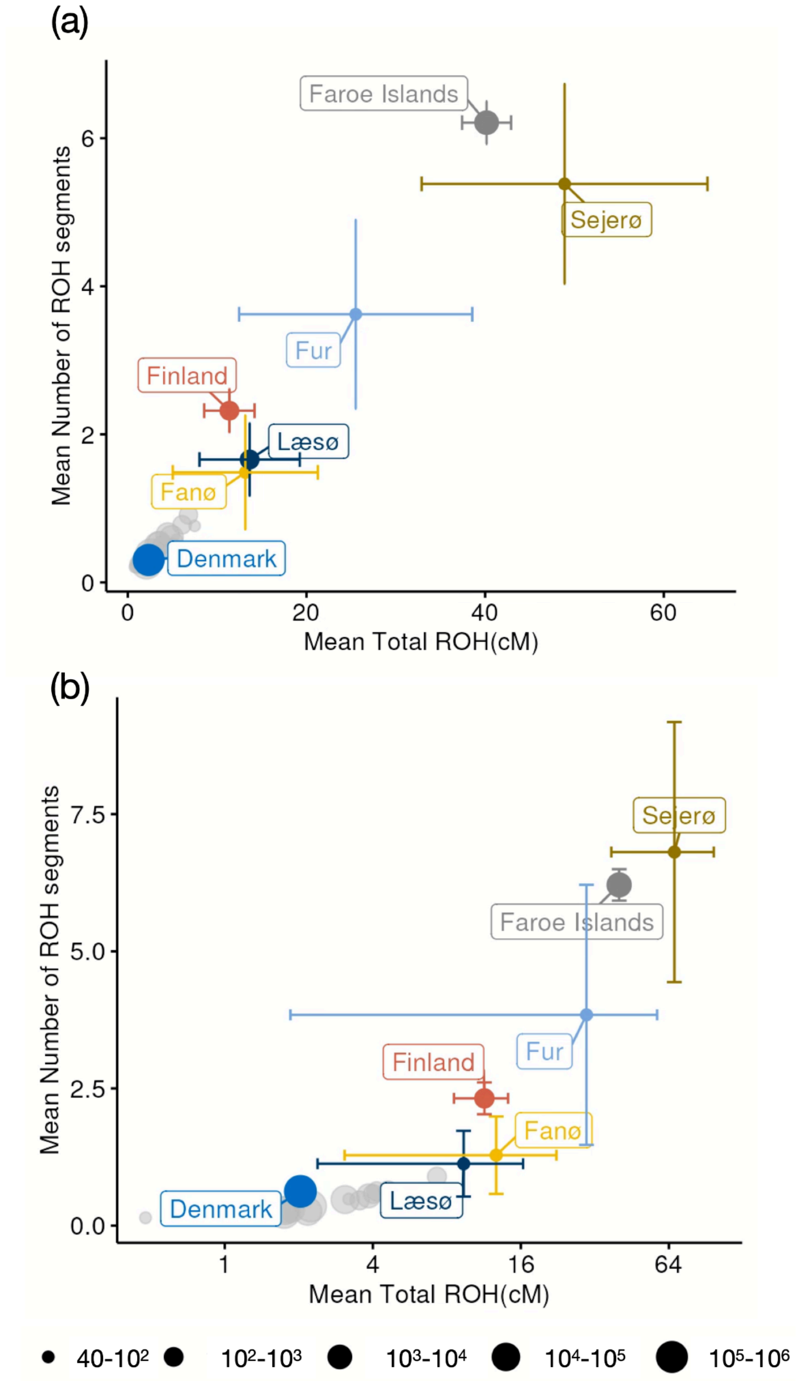


**Figure S10. Mean Total ROHs (X-axis) and mean number of ROH segments (Y-axis) of samples from islands of Denmark (only include those with sample size larger than 10, 24 islands in total), Denmark, the Faroe Islands, and Finland.** Error bars indicate the 95% confidence interval. Islands with mean total ROHs similar to or higher than Finland are highlighted, while others are shown in grey for reference. The dot sizes correspond to different scales of sample size. (a).The community ROHs of all available samples. (b)Out of concern for the results being biased by family-specific patterns, we also evaluated the ROHs in a subset of samples after filtering close relatives up to third degree (i.e., randomly removing one individual in a close relative pair). The mean of total ROHs and the mean number of ROH segments of the highlighted islands stay comparable to the levels of Finland after filtering, indicating likely community-wide patterns.

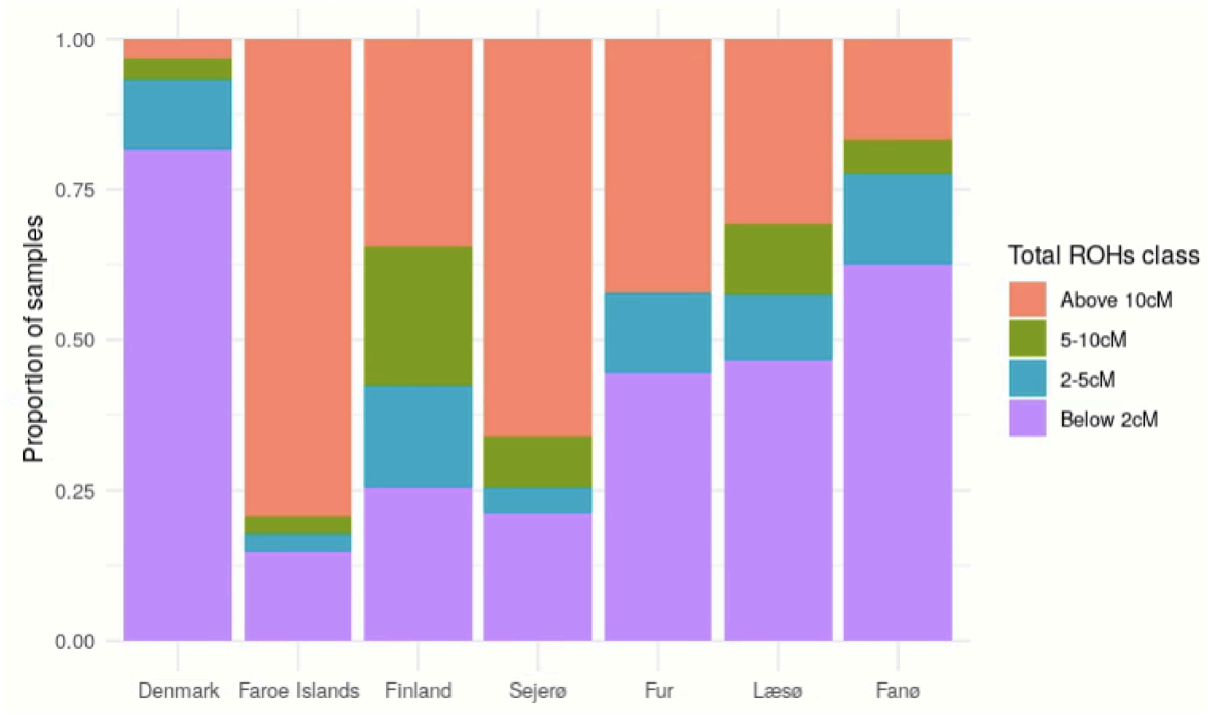


**Figure S11. Distribution of total length of ROHs across samples in selected populations: Denmark, Faroe Islands, Finland, Sejerø, Fur, Læsø, and Fanø.** The total length of ROHs is calculated per individual sample. For simplicity, we grouped the values into four classes as indicated in the legend. The calculation in this figure is based on all unrelated samples (the same set used in Figure S11(b)).

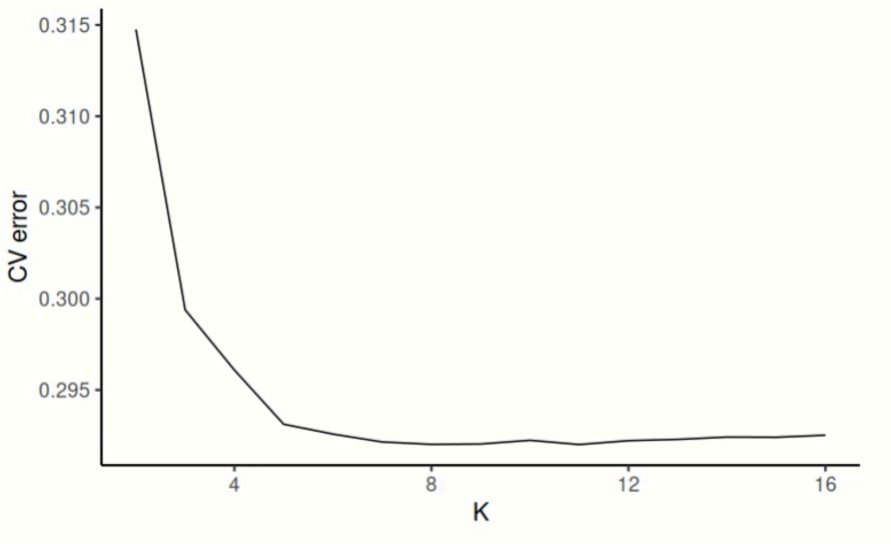


**Figure S12. 5-fold cross-validation error of ADMIXTURE unsupervised clustering across reference populations from 1kGP+HGDP.**

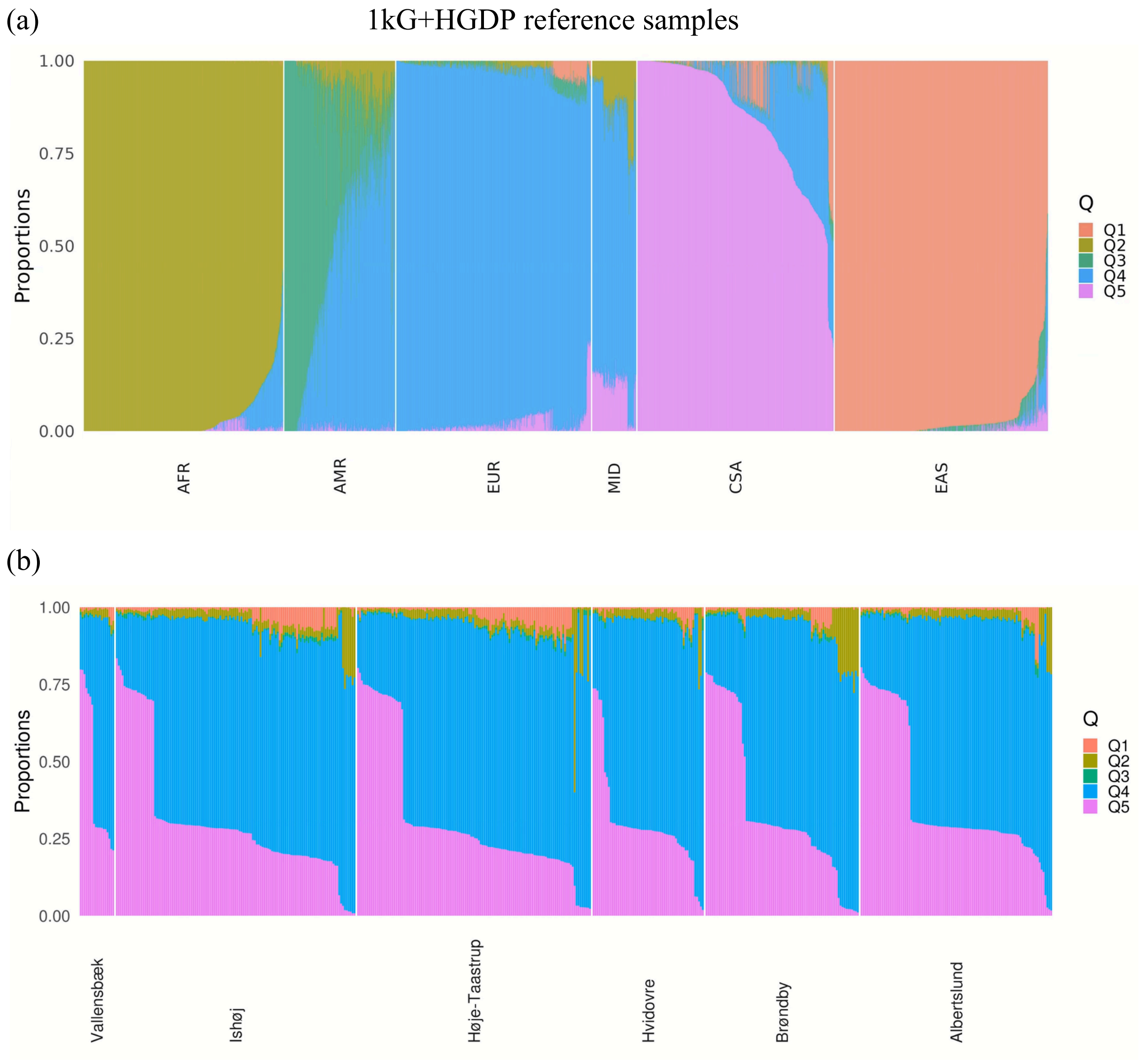


**Figure S13. ADMIXTURE analysis to infer continental ancestry components in the samples with high ROH (>10cM).** (a) Unsupervised clustering of 1kGP+HGDP reference samples from six super populations using ADMIXTURE with best-fitted number of ancestral components K=5. The super populations include African (AFR), Admixed American (AMR), European (EUR), Middle Eastern (MID), CSA (Central/South Asia), and East Asian (EAS). We find that the fitted ancestral components have different frequencies in these super populations, thus, they can be used to infer individual continental ancestries. (b)The estimated continental ancestry proportions of individuals with high levels of total ROH (>10cM) in the top-ranked communes with high ROH levels (n=503).

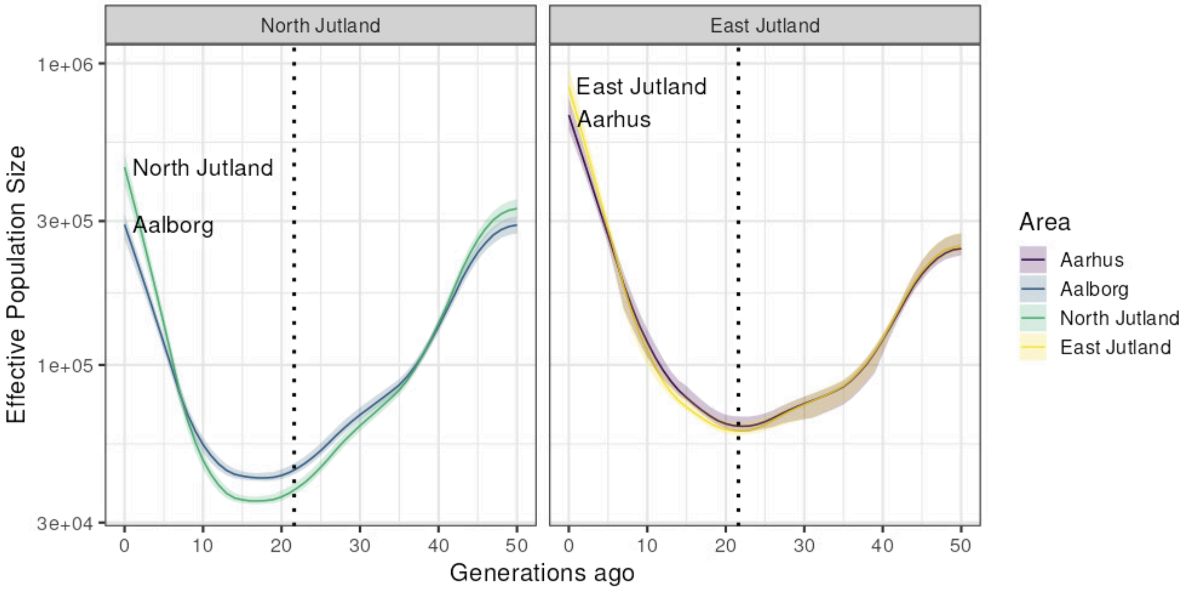


**Figure S14. Effective population size estimate in North Jutland and East Jutland across time (up to 50 generations ago)**. Aalborg and Aarhus are included as examples of urban areas in Jutland. The dotted line indicates the estimated generation age of the black death (21.6 generations ago) based on a generation time of 28 years.

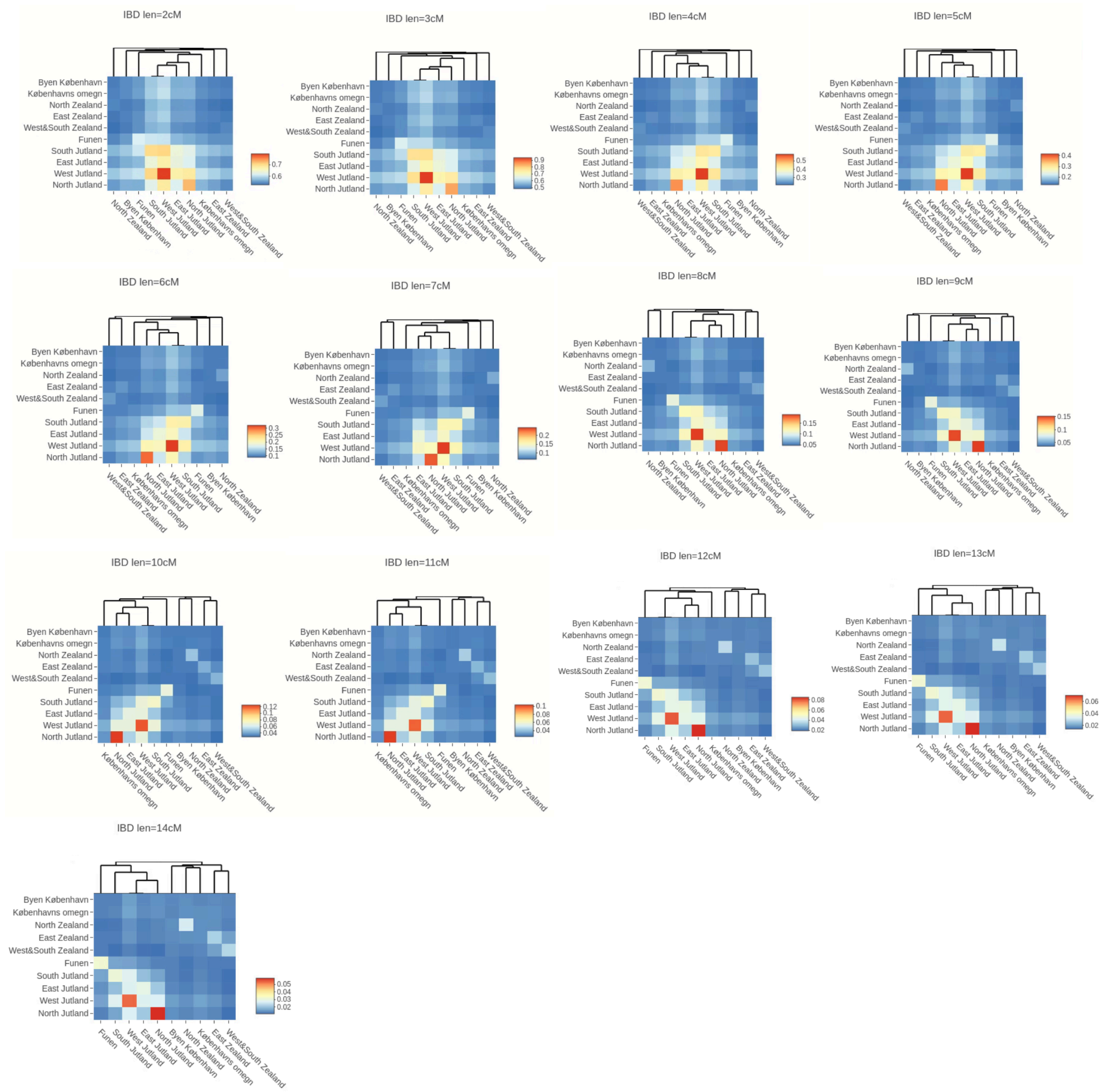


**Figure S15. IBD sharing across regions within Denmark across length bins of IBD segments (excluding Bornholm here). Each subplot corresponds to IBD sharing within the X and X+1 cM ($l \in [X, X+1)$ cM) range.**

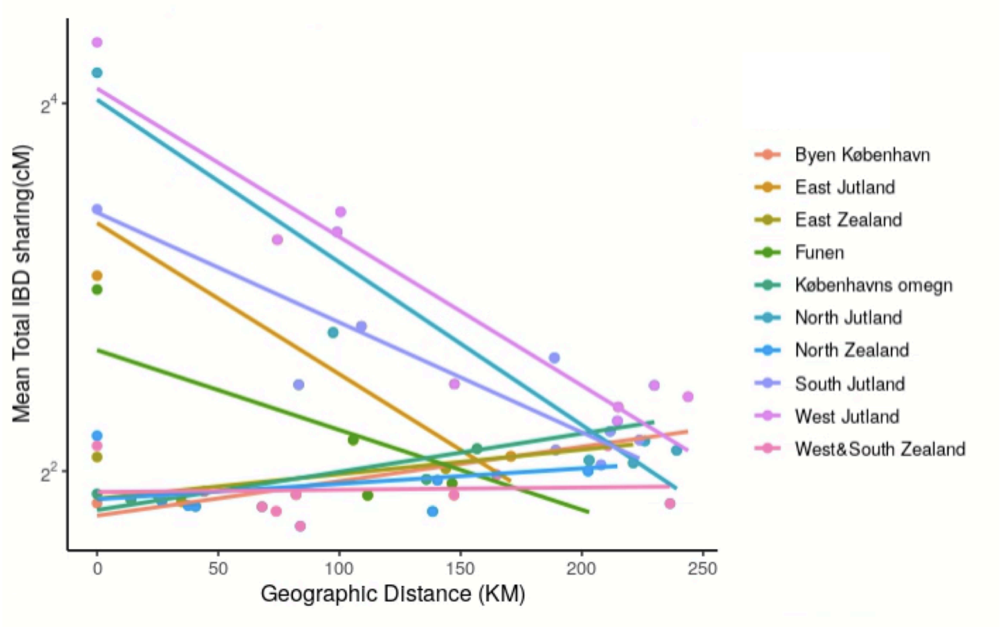


**Figure S16.** The relationship between geographic distance (X-axis) from a focal province to all the other provinces (each dot represents one) and their mean total IBD sharing (Y-axis) (except Bornholm). The linear line indicates the fitted linear regression model.

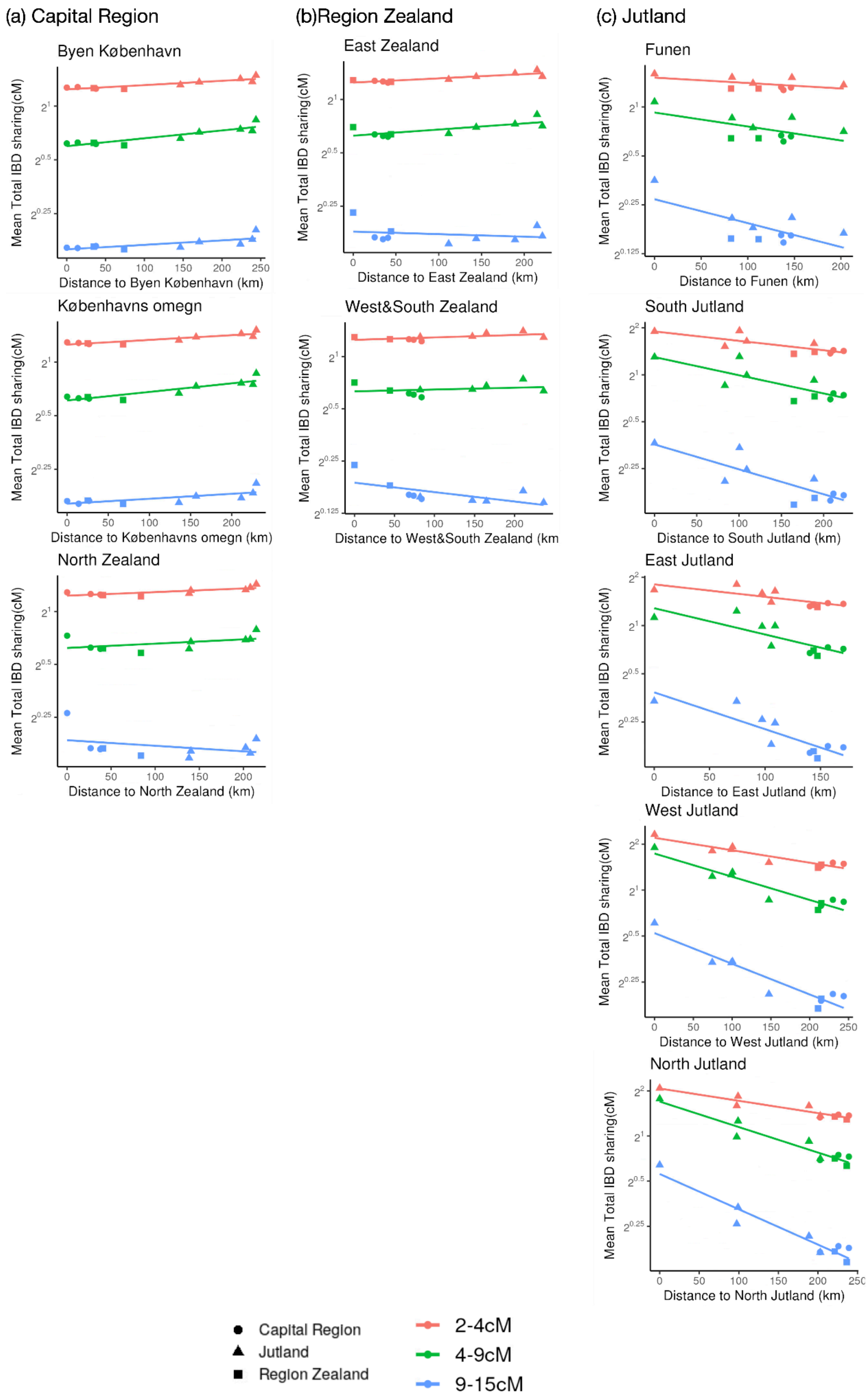

(a) Capital Region
(b)Region Zealand
(c) Jutland
Byen København
Distance to Byen København (km)
Københavns omegn
Distance to Københavns omegn (km)
North Zealand
Distance to North Zealand (km)
East Zealand
Distance to East Zealand (km)
West&South Zealand
Distance to West&South Zealand (km
Funen
Distance to Funen (km)
South Jutland
Distance to South Jutland (km)
East Jutland
Distance to East Jutland (km)
West Jutland
Distance to West Jutland (km)
North Jutland
Distance to North Jutland (km)
Mean Total IBD sharing(cM)
Capital Region
Jutland
Region Zealand
2-4cM
4-9cM
9-15cM

**Figure S17.** The relationship between geographic distance from a focal province to all the other provinces in Denmark (X-axis; excluding Bornholm) and their mean total IBD sharing (Y-axis on a log2 scale), stratified by three ranges of IBD lengths. The shape of the points indicates the categories of the provinces being compared (either from the Capital Region, Region Zealand, or Jutland). The colored lines represent fitted linear regression models for each IBD length range. The fitted slopes and their confidence intervals are summarised in **Figure S18.**

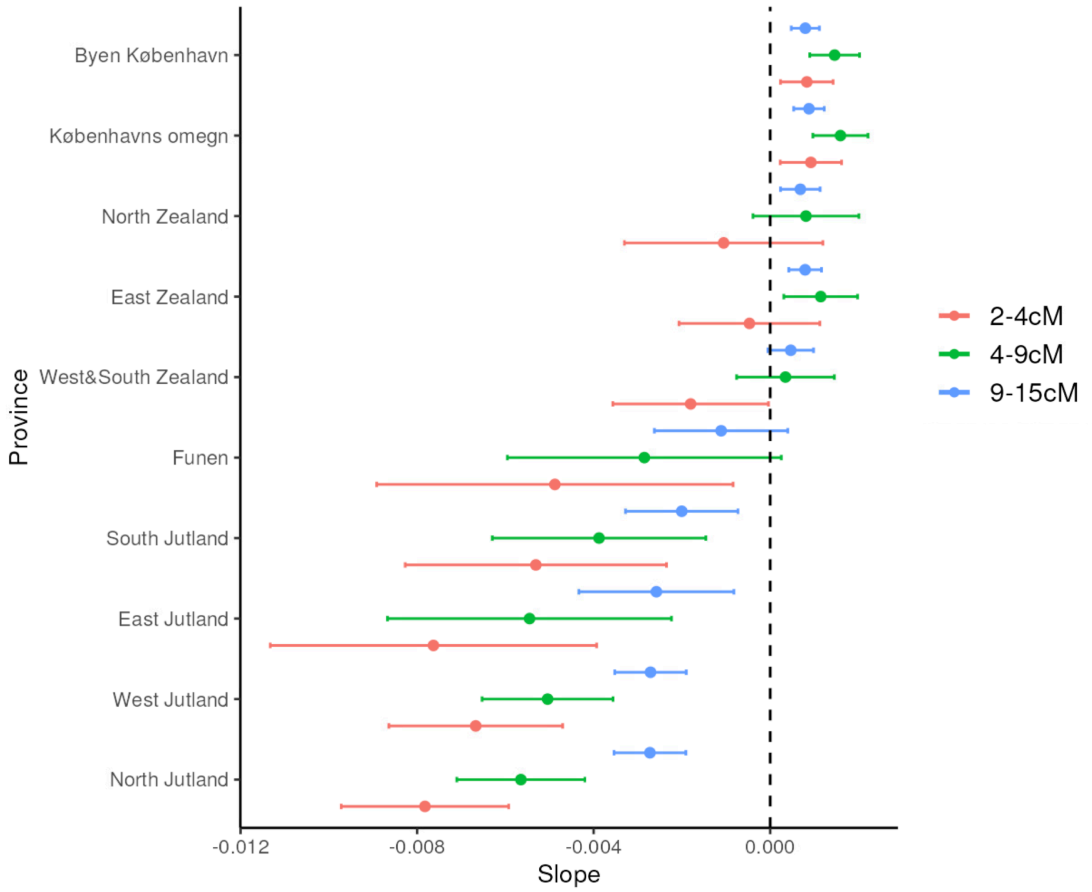


**Figure S18. The estimated slopes and 95% confidence intervals for the relationship between geographic distance from a focal province to all other provinces in Denmark (X-axis; excluding Bornholm) and their mean total IBD sharing (Y-axis, log 2 scale), stratified by three IBD length ranges.** A significantly negative slope indicates a decline in IBD sharing with increasing geographic distance, consistent with the isolation-by-distance model. In contrast, slopes that are positive or near zero suggest a weak or absence of the relationship between geographic distance and IBD sharing, indicating potential deviations from the isolation-by-distance pattern.

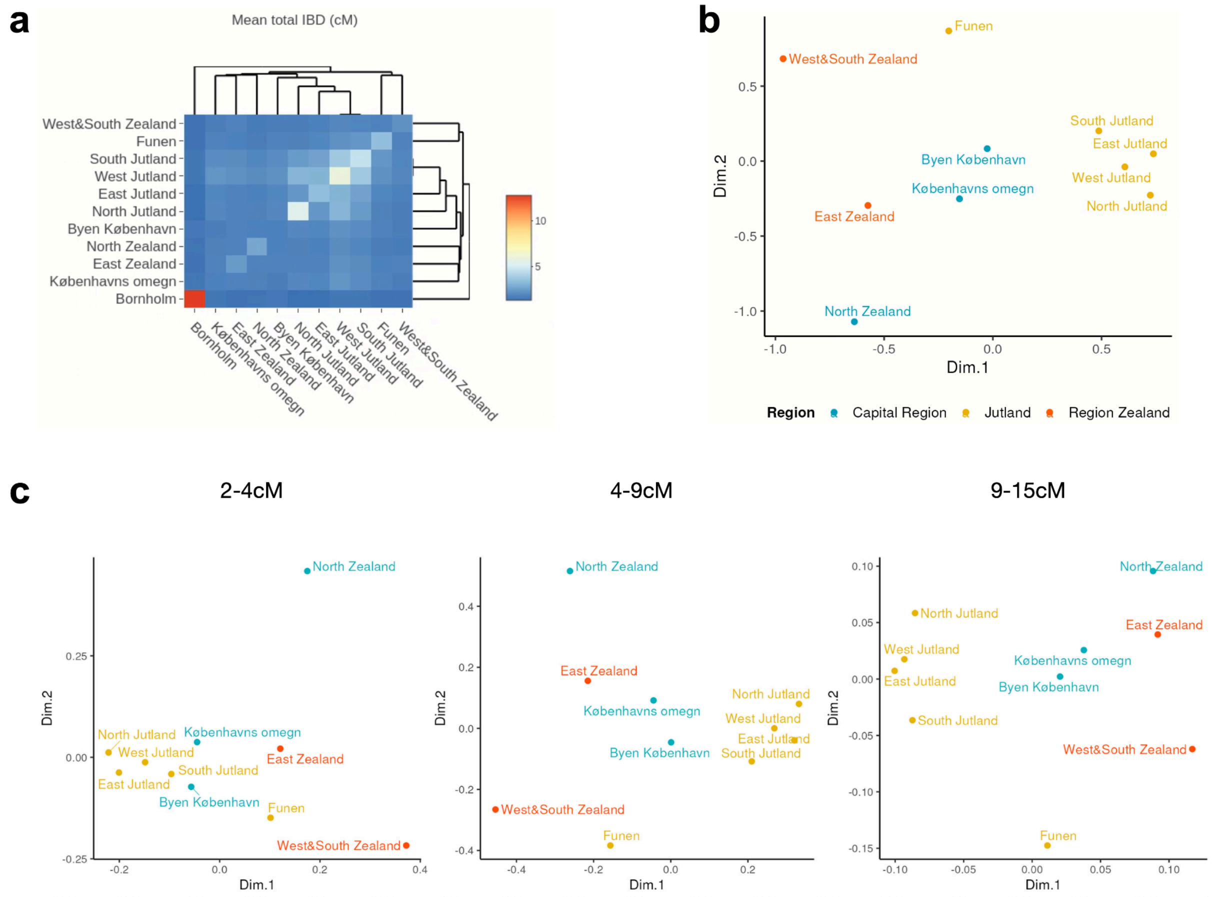


**Figure S19. Validation of IBD sharing patterns in Denmark using a refined subset of 28K individuals born in the same province as both of their parents.** (a) Pairwise total IBD sharing between provinces of Denmark. (b)Multidimensional scaling on the dissimilarity between provinces of Denmark (except Bornholm) transformed from pairwise total IBD sharing. (c) Multidimensional scaling on the dissimilarity between provinces of Denmark (except Bornholm) stratified by three ranges of IBD lengths.

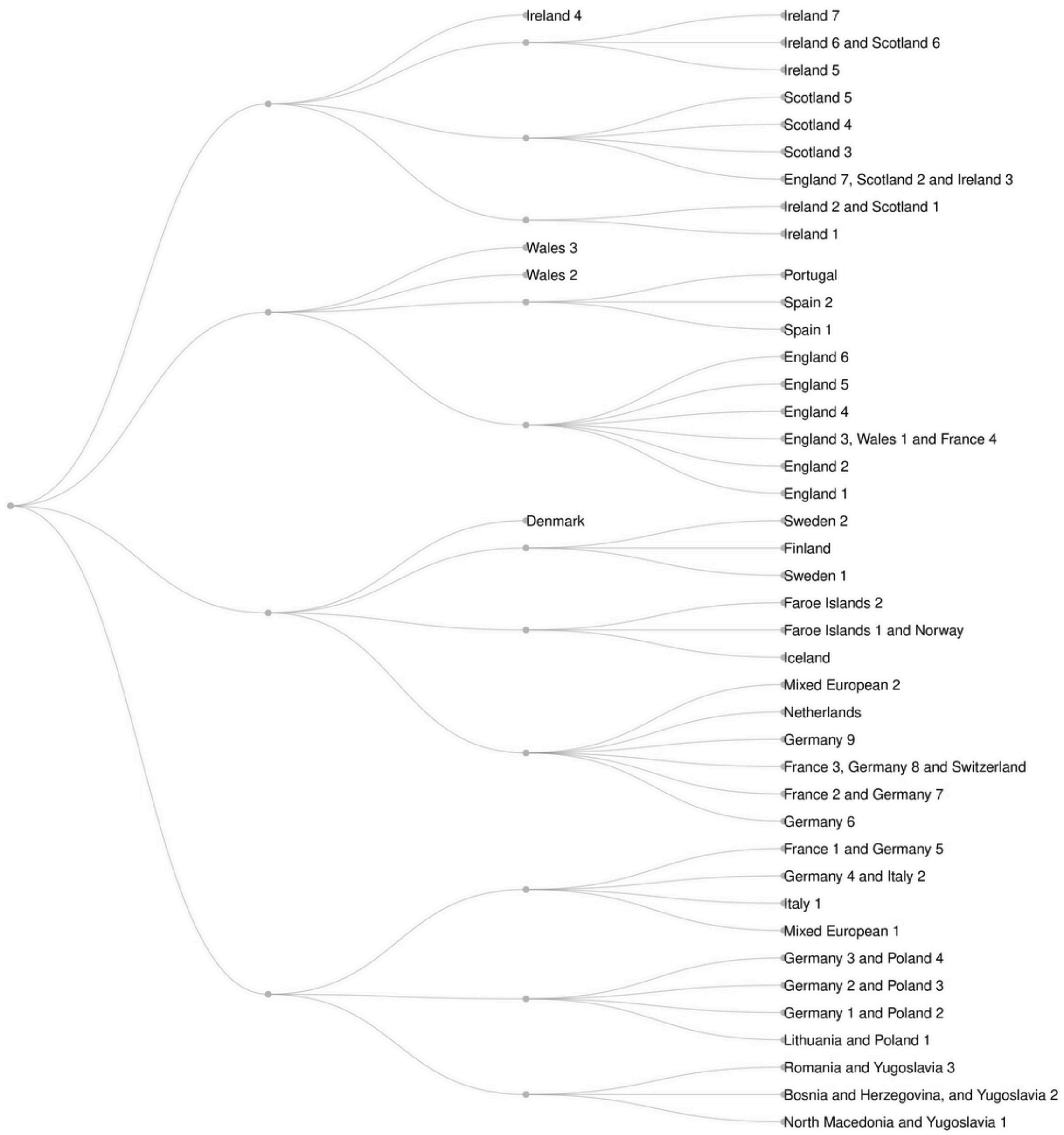


**Figure S20. Identification of European population clusters from UKBB+DNK datasets based on Leiden community clustering.**

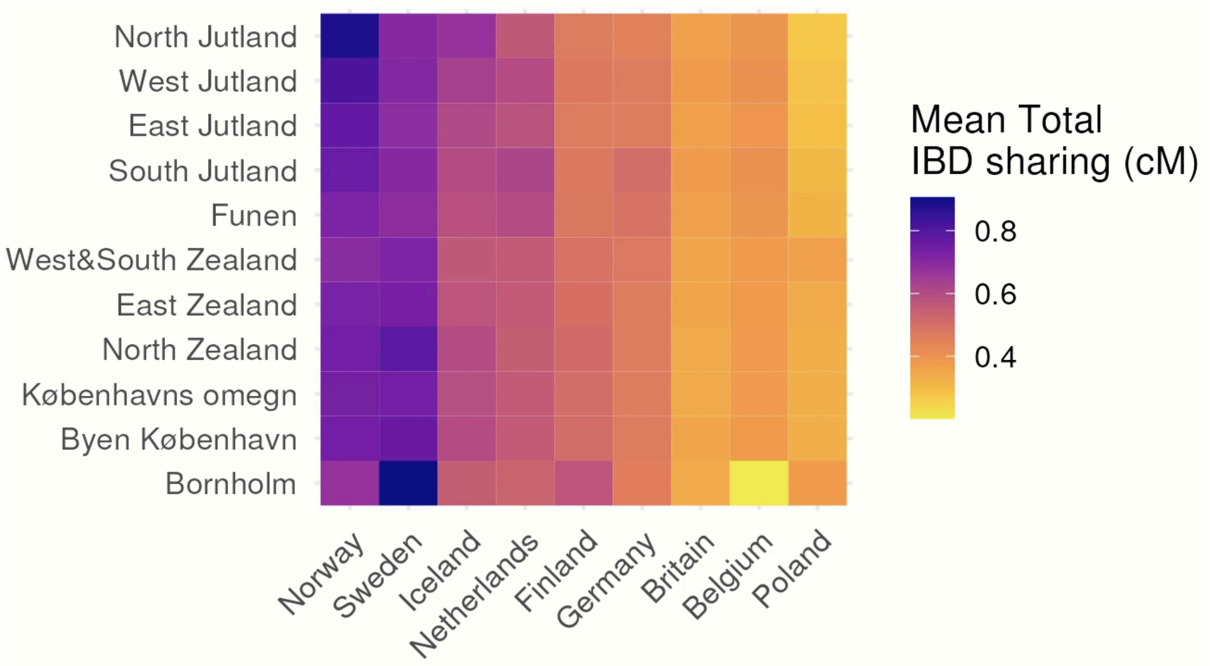


**Figure S21. Mean Total IBD sharing between provinces of Denmark and its neighbouring countries**. The colour scale is normalised across the neighbouring countries being compared.

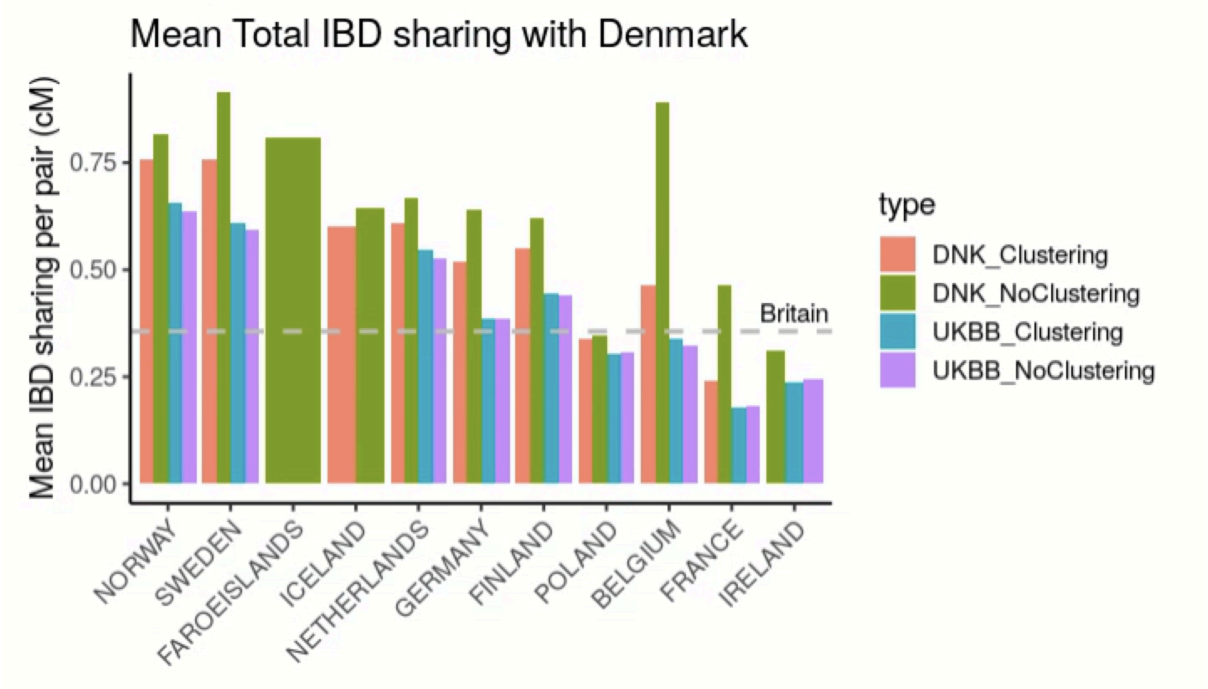


**Figure S22. Mean Total IBD sharing between Denmark and its neighbouring countries**. The dotted line indicates the value between Denmark and Britain. Using the samples identified in the Leiden community clustering approach, the results are largely qualitatively consistent regardless of the cohorts from which the samples are ascertained. Unless specified, we use the samples identified in clustering from the UKBB and DNK cohorts in the main text and other supplementary figures.

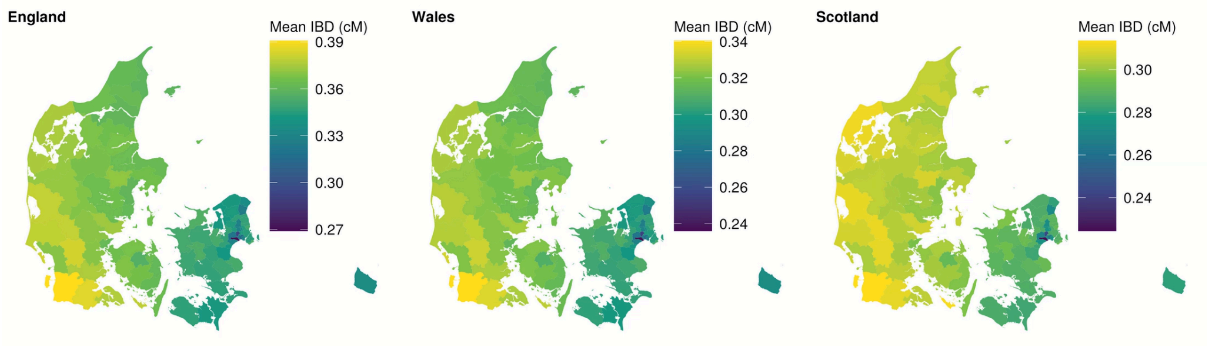


**Figure S23. Overall, IBD sharing between Danish populations at the municipality level and British populations stratified by English, Welsh and Scottish.**

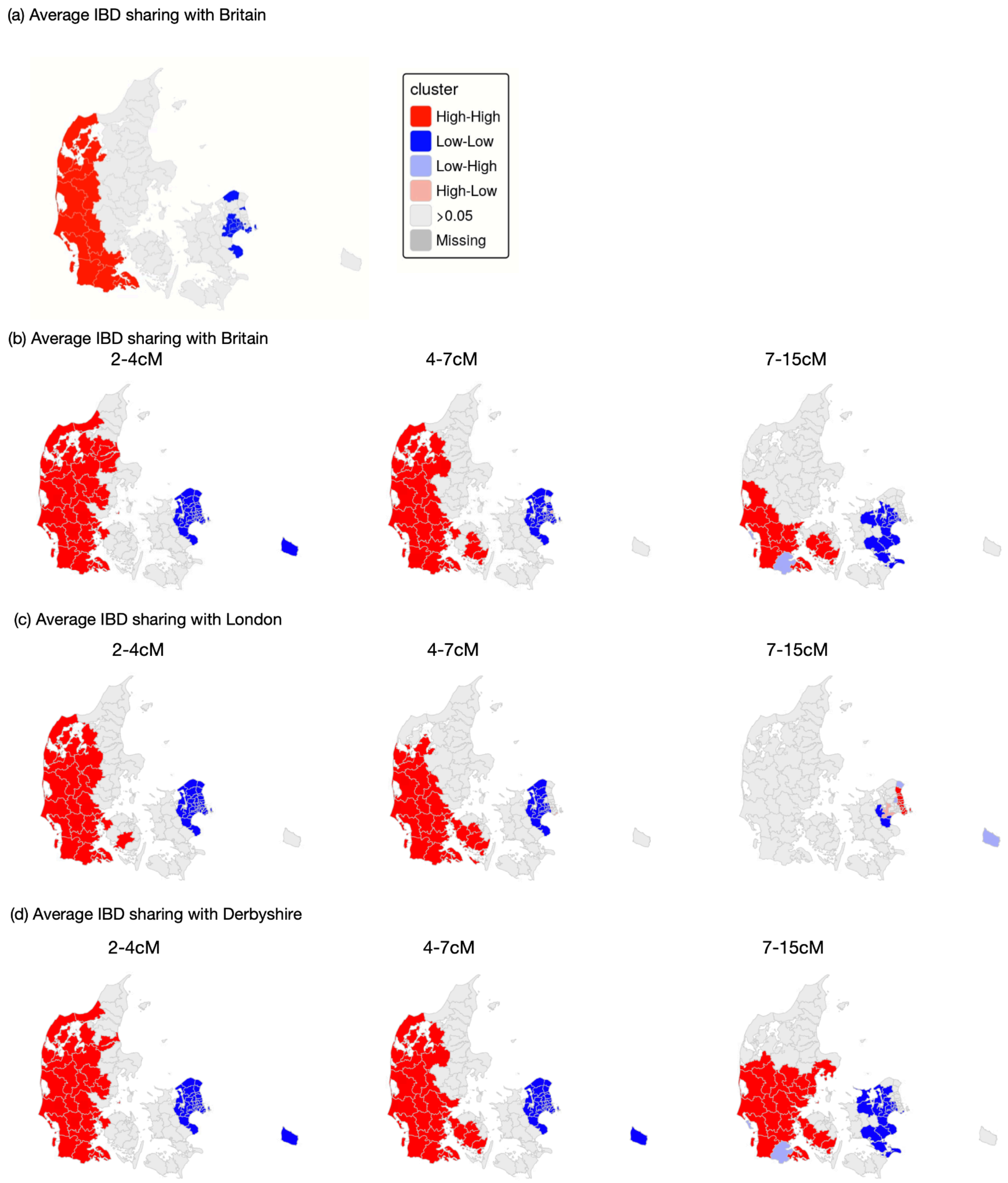


**Figure S24. Local Moran's test on spatial randomness of IBD sharing between Britain and regions of Denmark.**

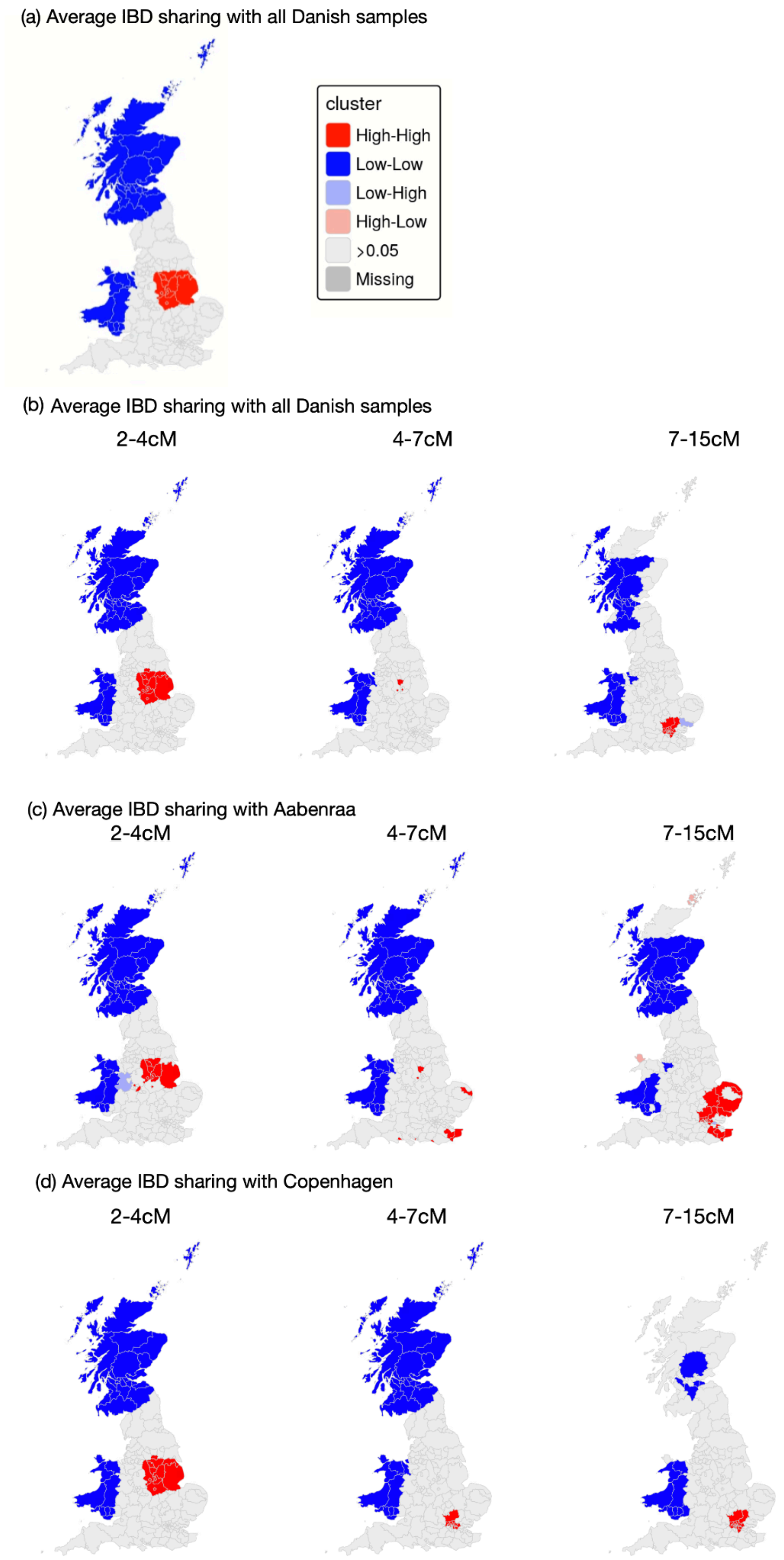


**Figure S25. Local Moran's test on spatial randomness of IBD sharing between Denmark and the regions of Britain.**

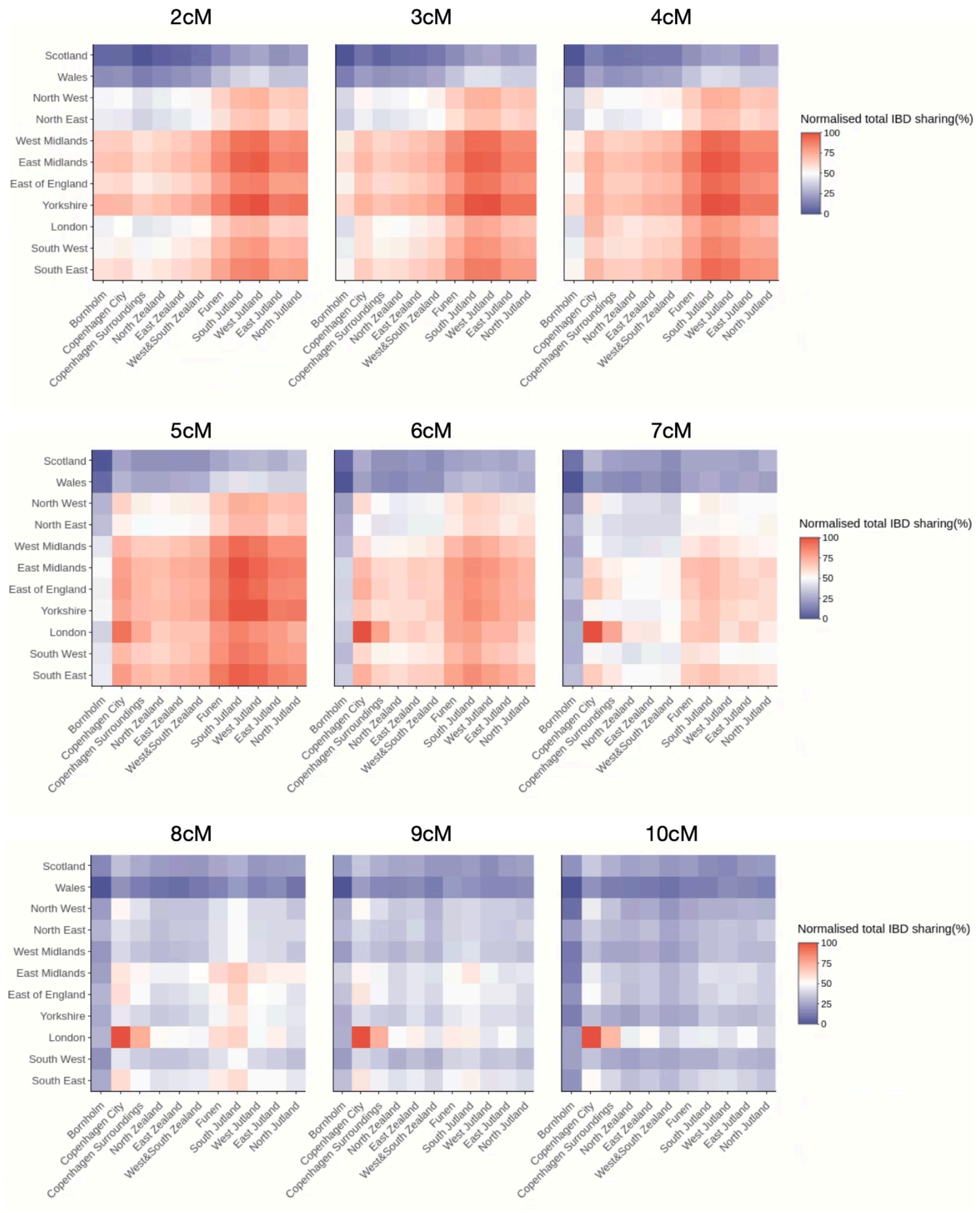


**Figure S26. IBD sharing across regions of the UK and Denmark across lengths of IBD segments between 2-10cM.**

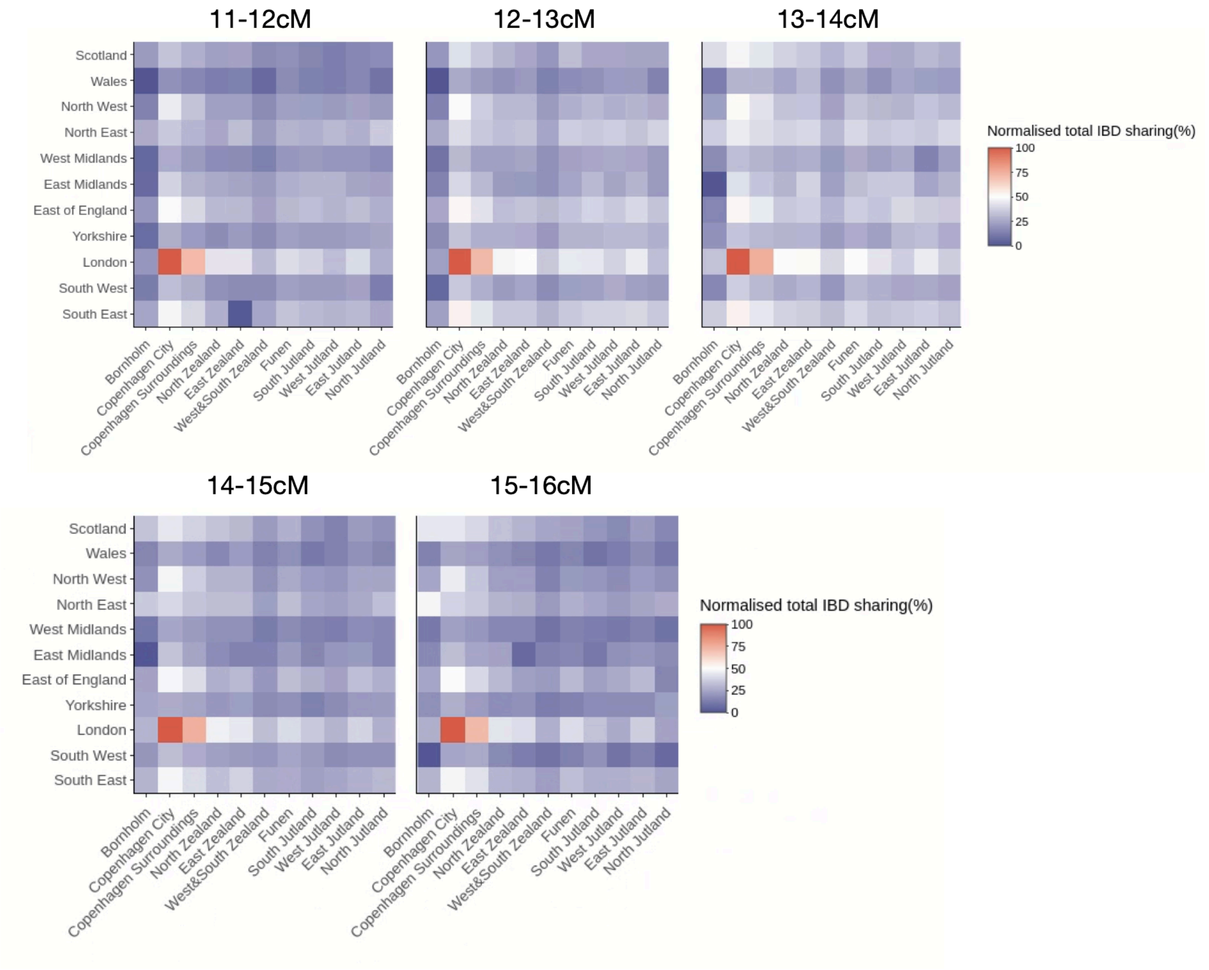


**Figure S27. IBD sharing across regions of the UK and Denmark, across lengths of IBD segments between 11-16cM.**

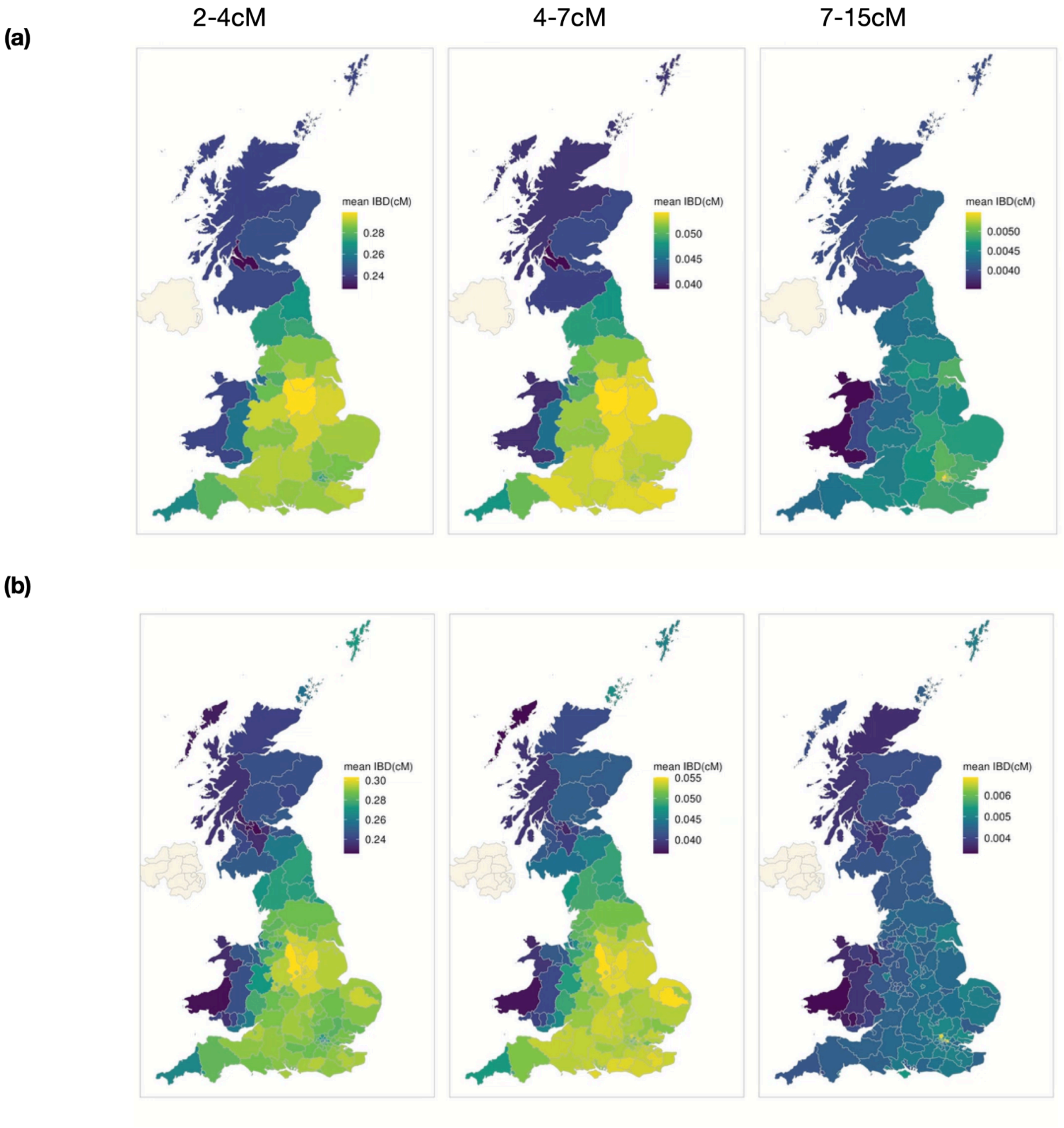


**Figure S28. Mean IBD sharing between mainland Denmark and regions of the UK aggregated across varying lengths of IBD segments (unit: cM) is shown at the UK NUTS Level 2 (top) and NUTS Level 3 (bottom).**

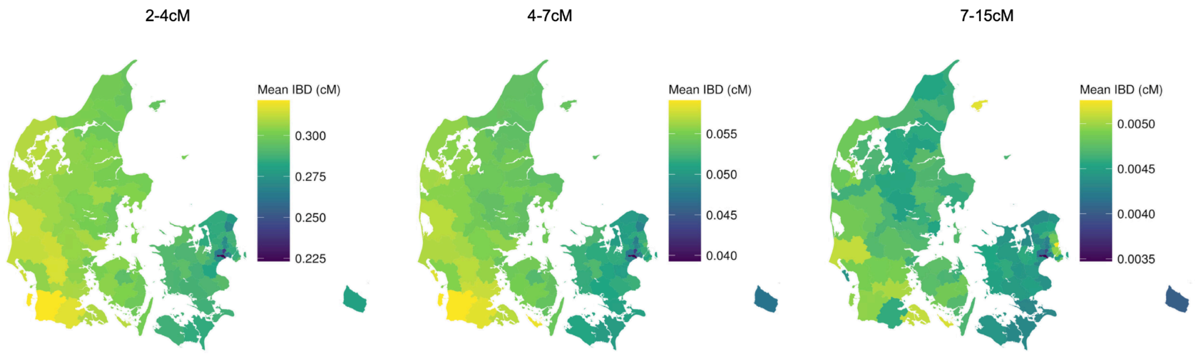


**Figure S29. Mean IBD sharing between Britain and regions of Denmark aggregated across varying lengths of IBD segments (unit: cM).**

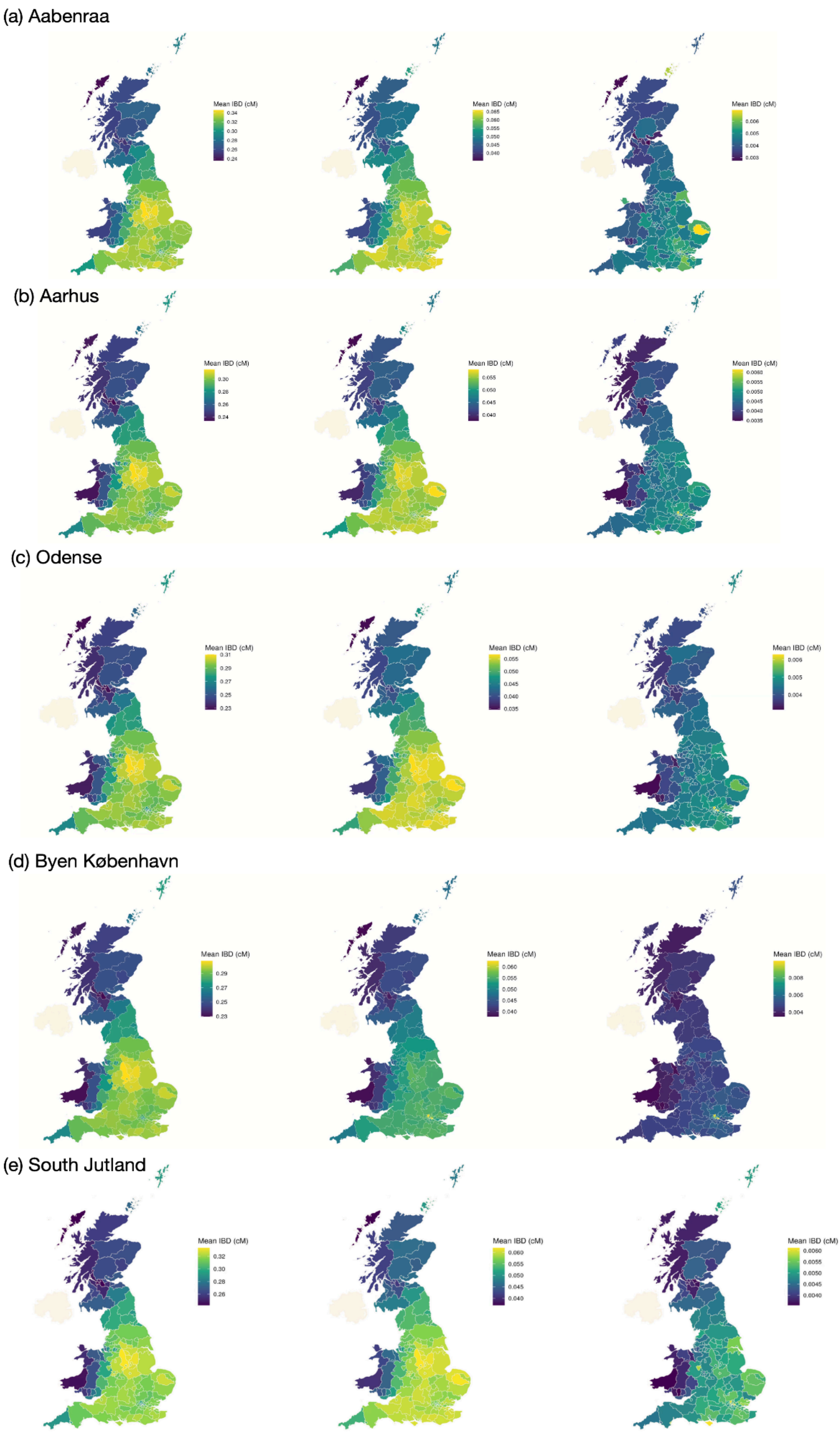
(a) Aabenraa
Mean IBD (cM)
0.34
0.32
0.30
0.28
0.26
0.24
Mean IBD (cM)
0.065
0.060
0.055
0.050
0.045
0.040
Mean IBD (cM)
0.006
0.005
0.004
0.003
(b) Aarhus
Mean IBD (cM)
0.30
0.28
0.26
0.24
Mean IBD (cM)
0.055
0.050
0.045
0.040
Mean IBD (cM)
0.0060
0.0055
0.0050
0.0045
0.0040
0.0035
(c) Odense
Mean IBD (cM)
0.31
0.29
0.27
0.25
0.23
Mean IBD (cM)
0.055
0.050
0.045
0.040
0.035
Mean IBD (cM)
0.006
0.005
0.004
(d) Byen København
Mean IBD (cM)
0.29
0.27
0.25
0.23
Mean IBD (cM)
0.060
0.055
0.050
0.045
0.040
Mean IBD (cM)
0.008
0.006
0.004
(e) South Jutland
Mean IBD (cM)
0.32
0.30
0.28
0.26
Mean IBD (cM)
0.060
0.055
0.050
0.045
0.040
Mean IBD (cM)
0.0060
0.0055
0.0050
0.0045
0.0040

**Figure S30. IBD sharing between British populations and industrial cities in Denmark: Aabenraa, Aarhus, Odense, København and South Jutland across three ranges of IBD segment length: 2-4cM, 4-7cM, and 7-15cM.**

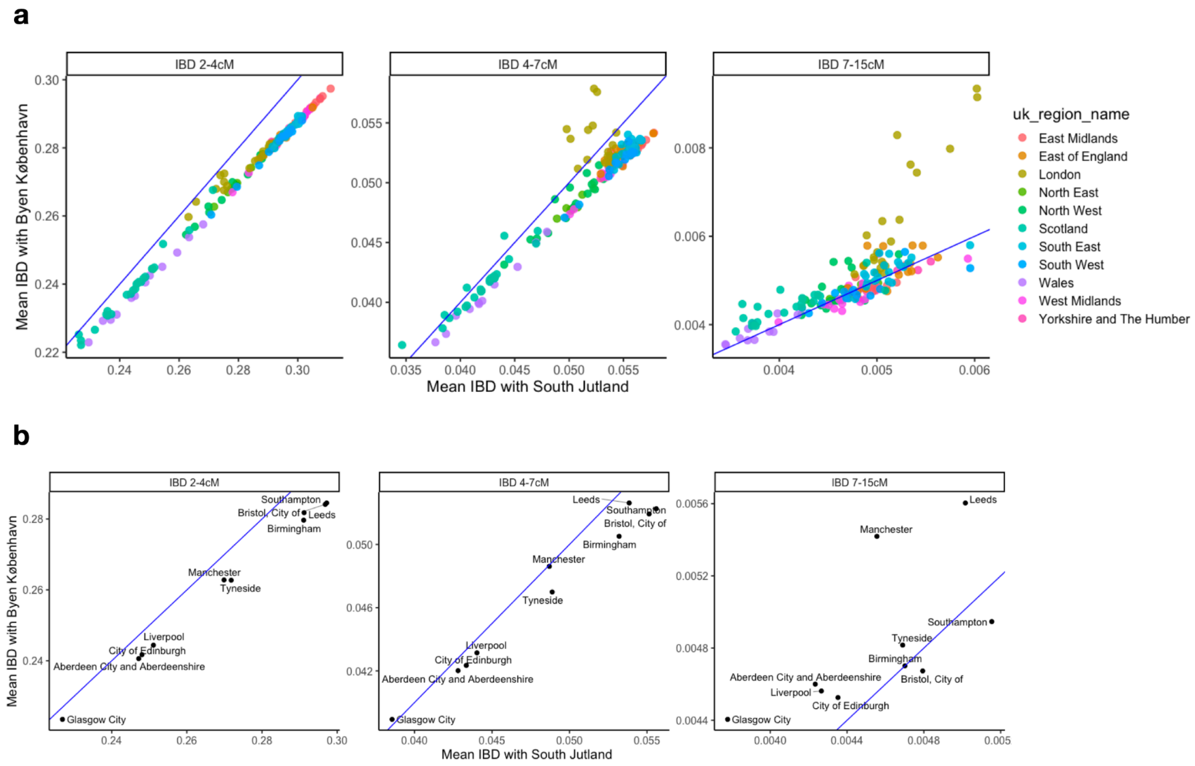


**Figure S31. Comparison of mean IBD sharing of UK municipalities (defined by NUTS Level 3) with South Jutland vs København across different ranges of IBD lengths**. The units for both the X-axis and Y-axis are in cM. The blue lines for all plots correspond to y=x, which is used as a reference here. A dot of municipality above the line shows it has higher IBD sharing with København than South Jutland, and vice versa. Panel (a) includes all municipalities. We see that multiple municipalities in London have increasing IBD sharing with København for the mid and long IBD segments. Panel (b) highlights other selected cities in Britain.

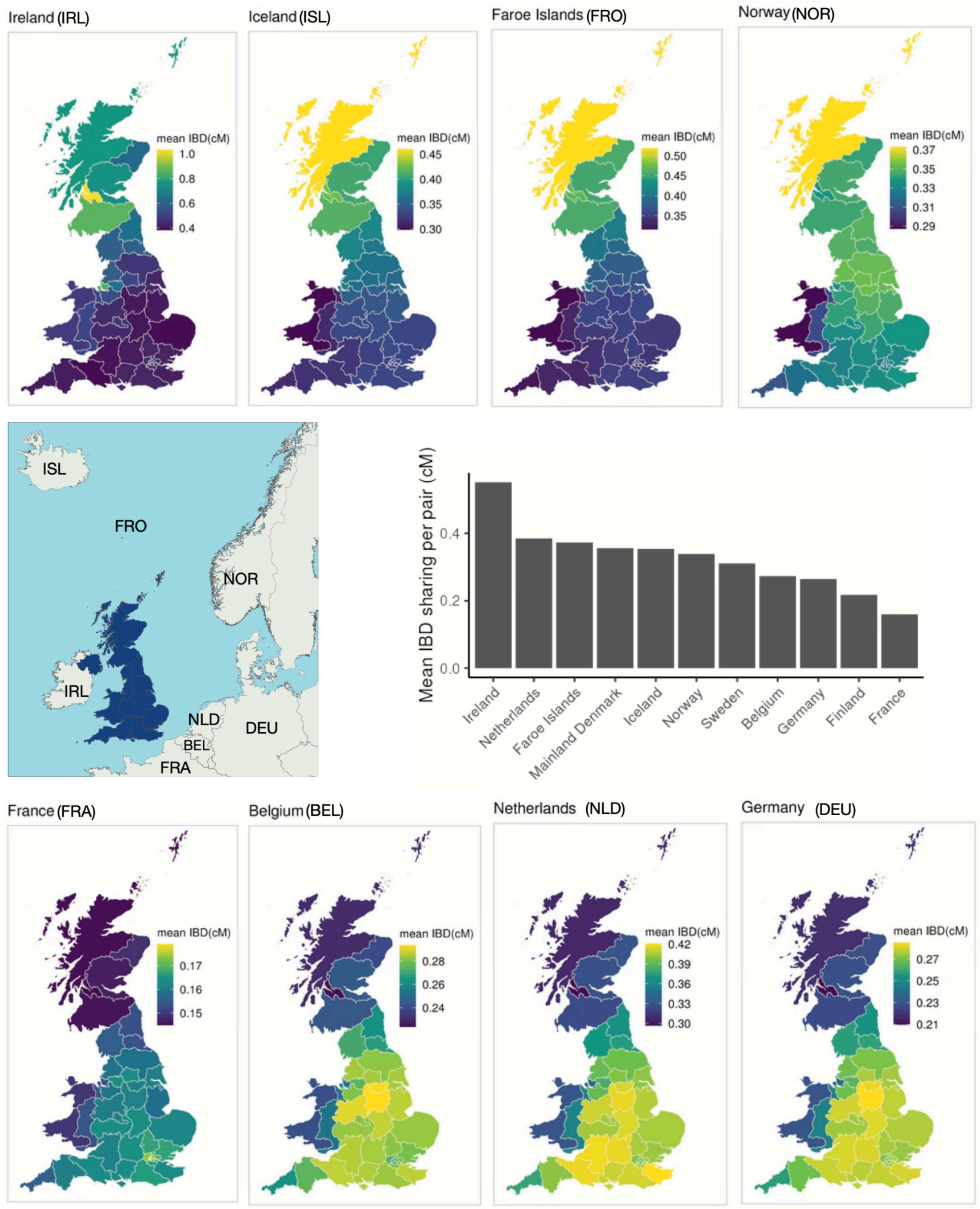


**Figure S32. Mean IBD sharing between British and other neighbouring European populations is shown at the UK NUTS Level 2.**

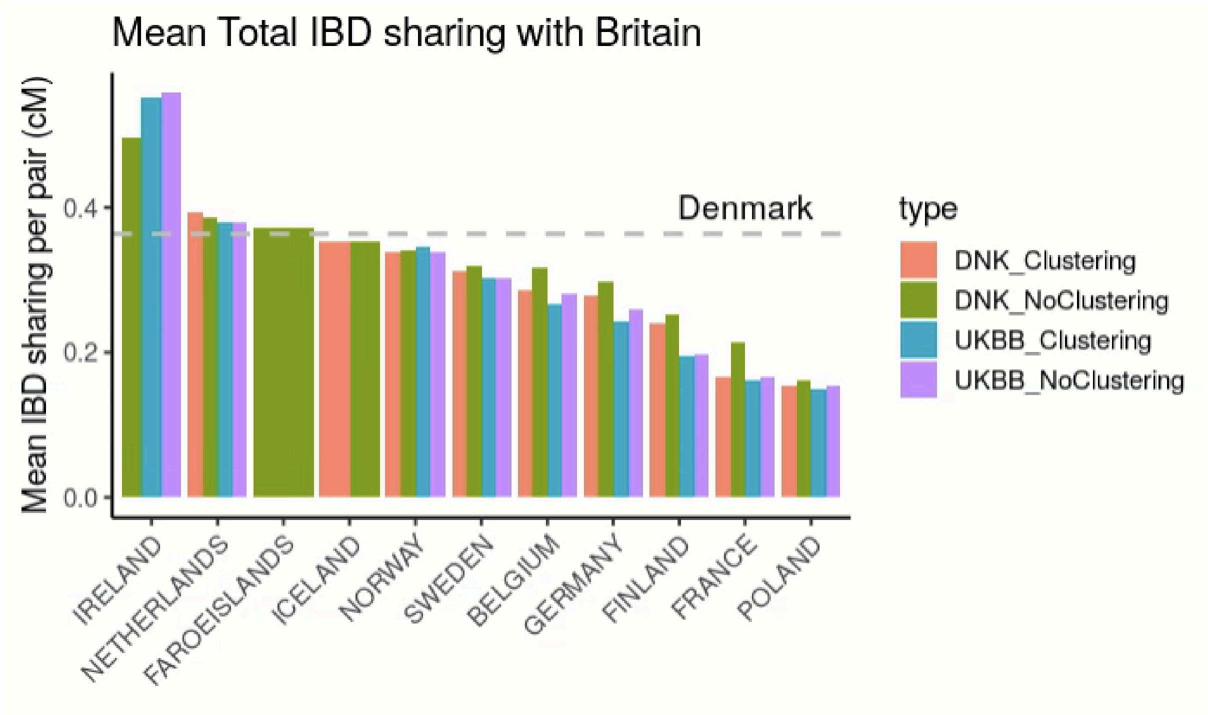


**Figure S33. Mean Total IBD sharing between Britain and its neighbouring countries**. The dotted line indicates the value between Britain and Denmark. Using samples identified from the Leiden community clustering approach, the results are qualitatively consistent regardless of the cohorts from which the samples are ascertained. Unless specified, we use the samples identified in clustering from the UKBB and DNK cohorts in the main text and other supplementary figures.

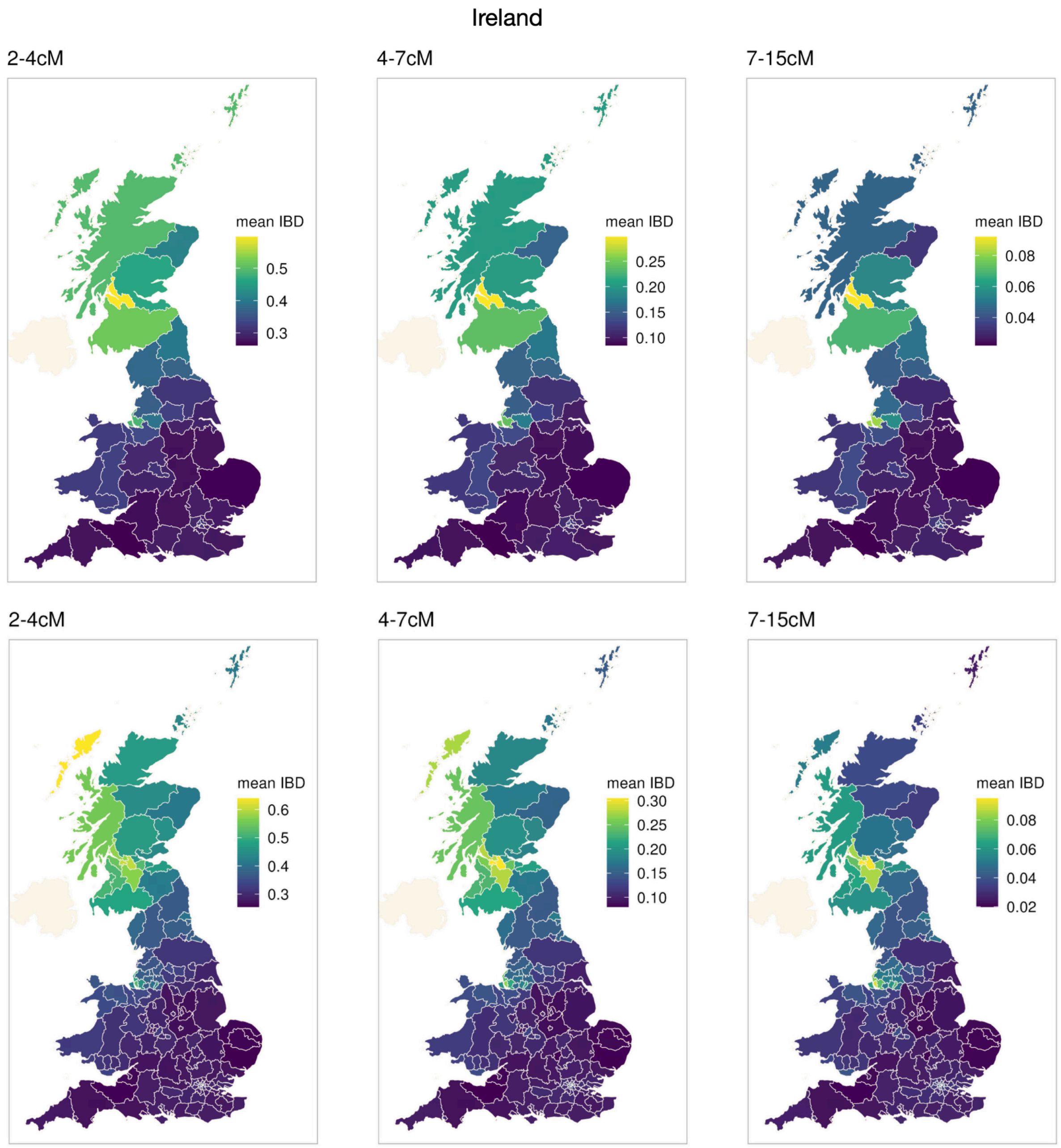


**Figure S34. Mean IBD sharing between the population of Ireland and regions of the UK aggregated across varying lengths of IBD segments (unit: cM) is shown at the UK NUTS Level 2 (top) and NUTS Level 3 (bottom).** Scotland and North West England have the highest overall IBD sharing with Ireland compared to other regions in England and Wales. At the NUTS level 2, West Scotland, particularly Glasgow in West Central Scotland, shows consistently high IBD sharing across all periods. In Merseyside, especially Liverpool, there is a notable increase in IBD sharing with Ireland for segments between 7–15 cM (MAP age=1555AD-1768AD, 50% HDI = 1325AD-1843AD).

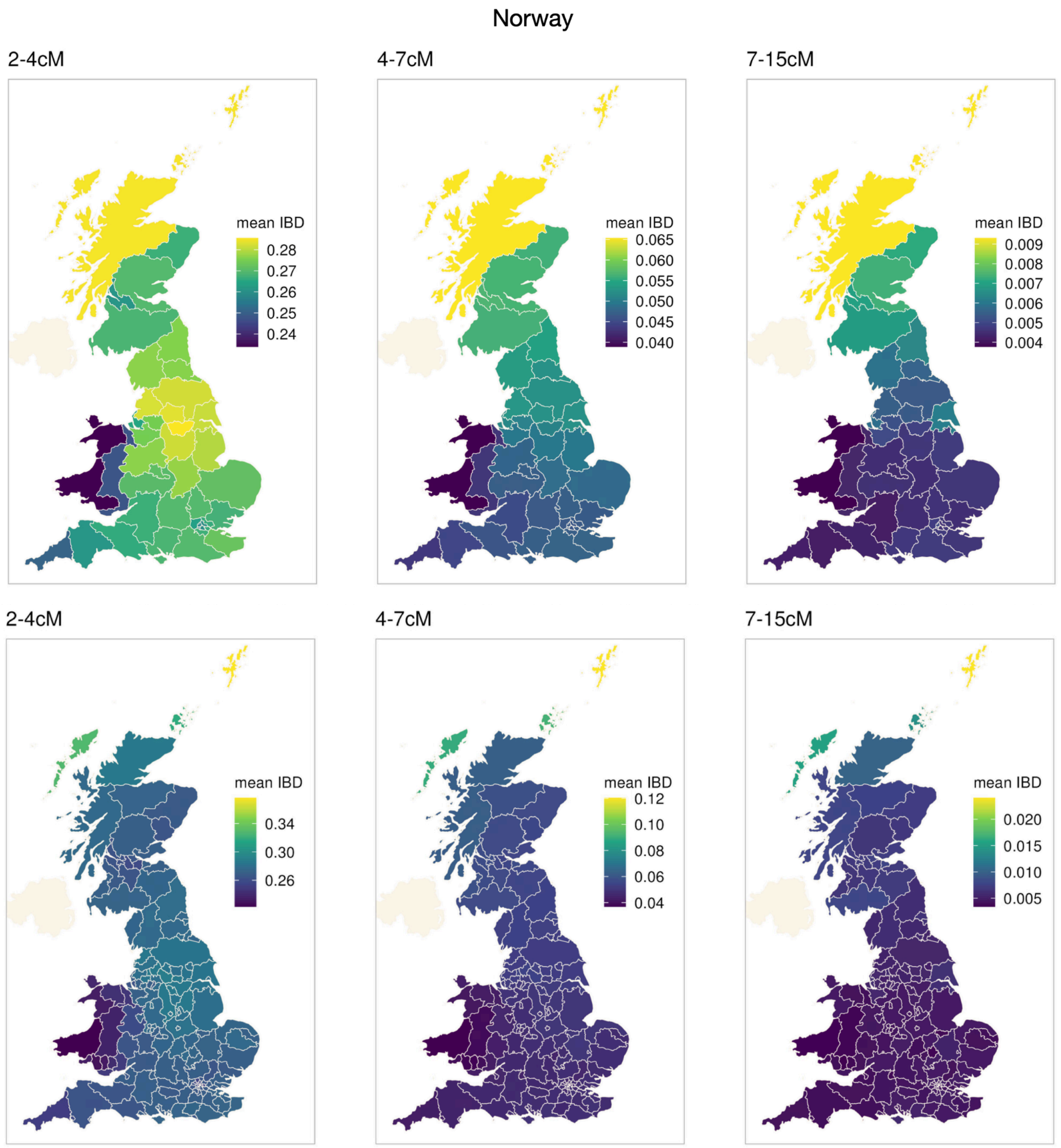


**Figure S35. Mean IBD sharing between the population of Norway and regions of the UK aggregated across varying lengths of IBD segments (unit: cM) is shown at the UK NUTS Level 2 (top) and NUTS Level 3 (bottom).** The Highlands and Islands of Scotland, especially Shetland, have the highest overall IBD sharing with Norway compared to England and Wales throughout all periods. We also noted regions around South Yorkshire are another hot spot of IBD sharing for segments between 2-4cM (MAP age = 555AD - 1255AD, 50% HDI =257 BC - 1549AD), which is similar to the IBD sharing pattern of the British with Danes.

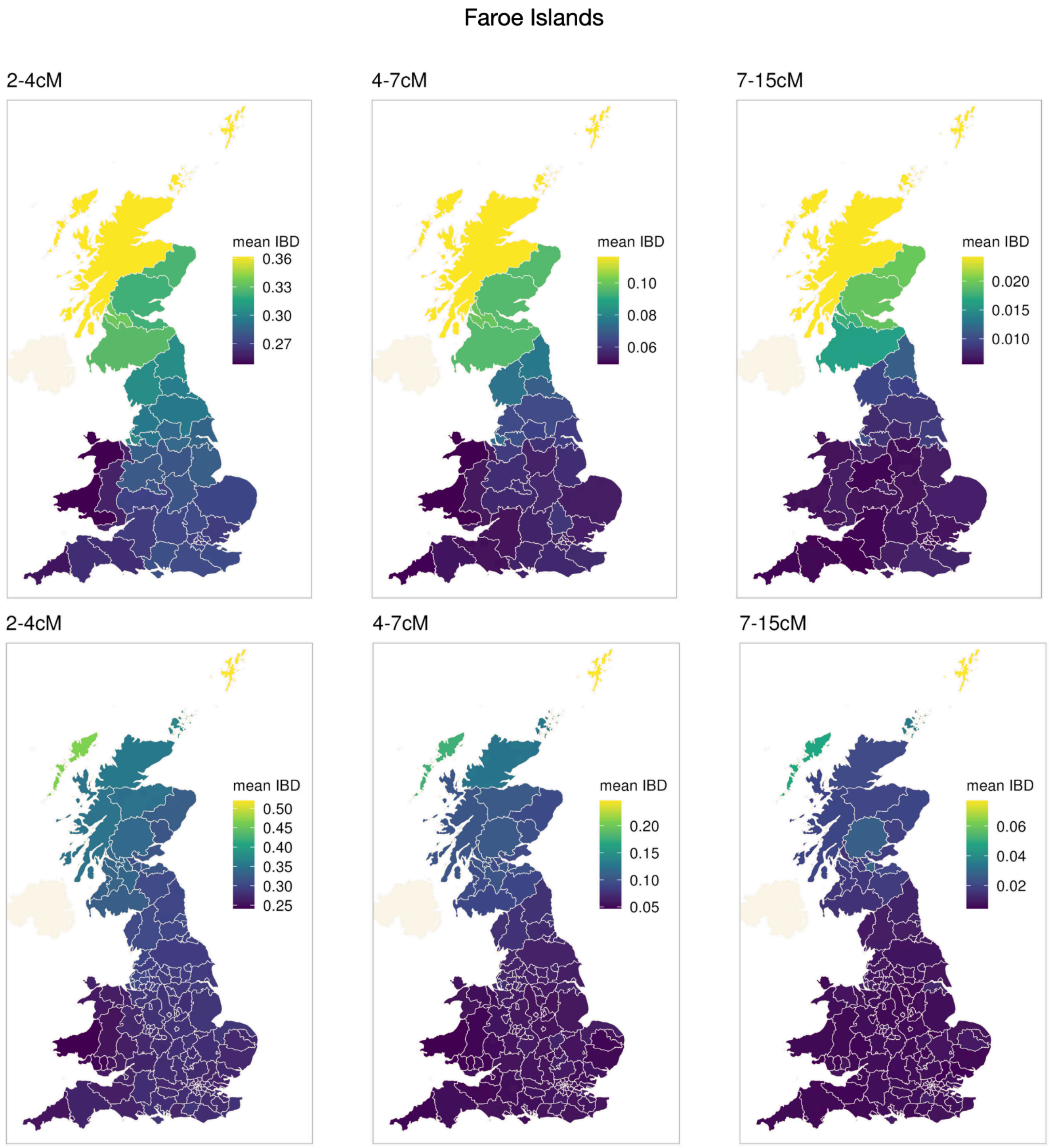


**Figure S36. Mean IBD sharing between the population of the Faroe Islands and regions of the UK aggregated across varying lengths of IBD segments (unit: cM) is shown at the UK NUTS Level 2 (top) and NUTS Level 3 (bottom).** The Highlands and Islands of Scotland, especially Shetland, have the highest overall IBD sharing with Faroe Islanders compared to England and Wales throughout all periods.

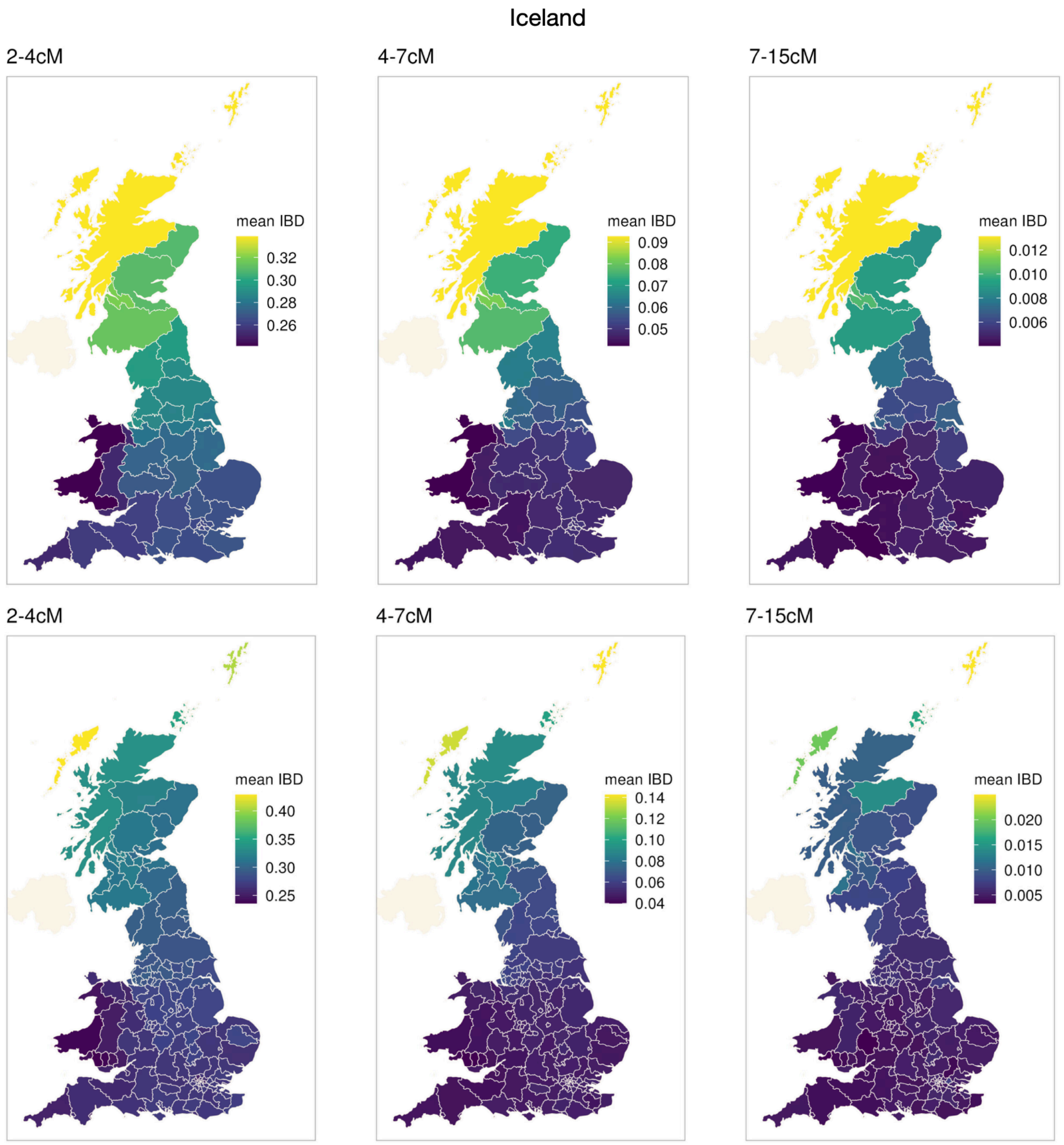


**Figure S37. Mean IBD sharing between the population of Iceland and regions of the UK aggregated across varying lengths of IBD segments (unit: cM) is shown at the UK NUTS Level 2 (top) and NUTS Level 3 (bottom).** The Highlands and Islands of Scotland have the highest overall IBD sharing compared to England and Wales throughout all periods, following a similar pattern to the Faroe Islanders.

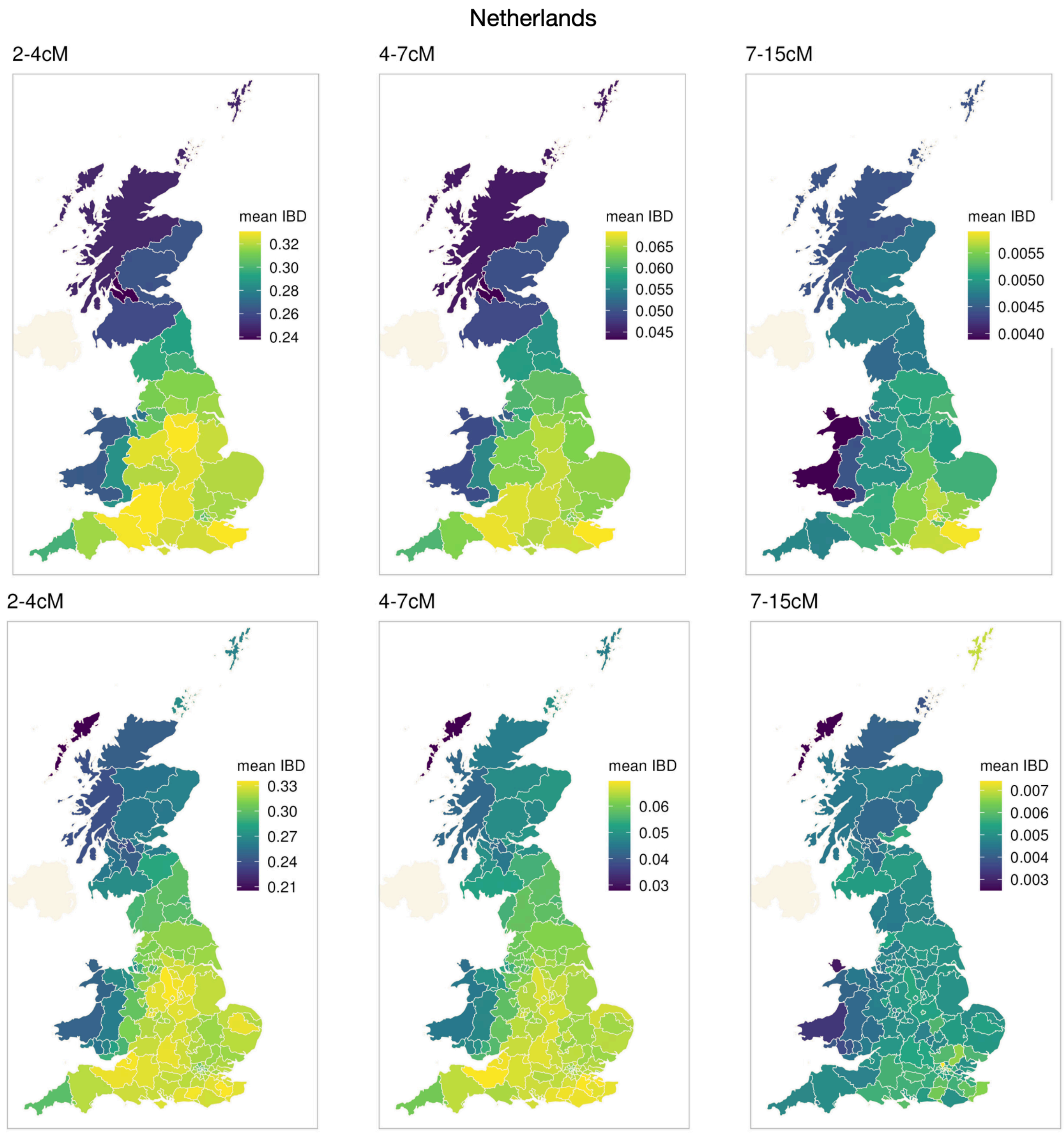


**Figure S38. Mean IBD sharing between populations of the Netherlands and regions of the UK aggregated across varying lengths of IBD segments (unit: cM) is shown at the UK NUTS Level 2 (top) and NUTS Level 3 (bottom).** Derbyshire has high IBD sharing with the Dutch for segments between 2-4cM (MAP age = 555AD - 1255AD, 50% credible interval = 257 BC - 1549AD). The South Coast of England (spanning from Kent and up to Somerset and Dorset), especially Kent (South East), also has high IBD sharing with the Dutch throughout all lengths of IBD segments. For long IBD segments (7-15cM, MAP age=1555AD-1768AD, 50% credible = 1325AD-1843AD), we also find increased IBD sharing between London and the Dutch.

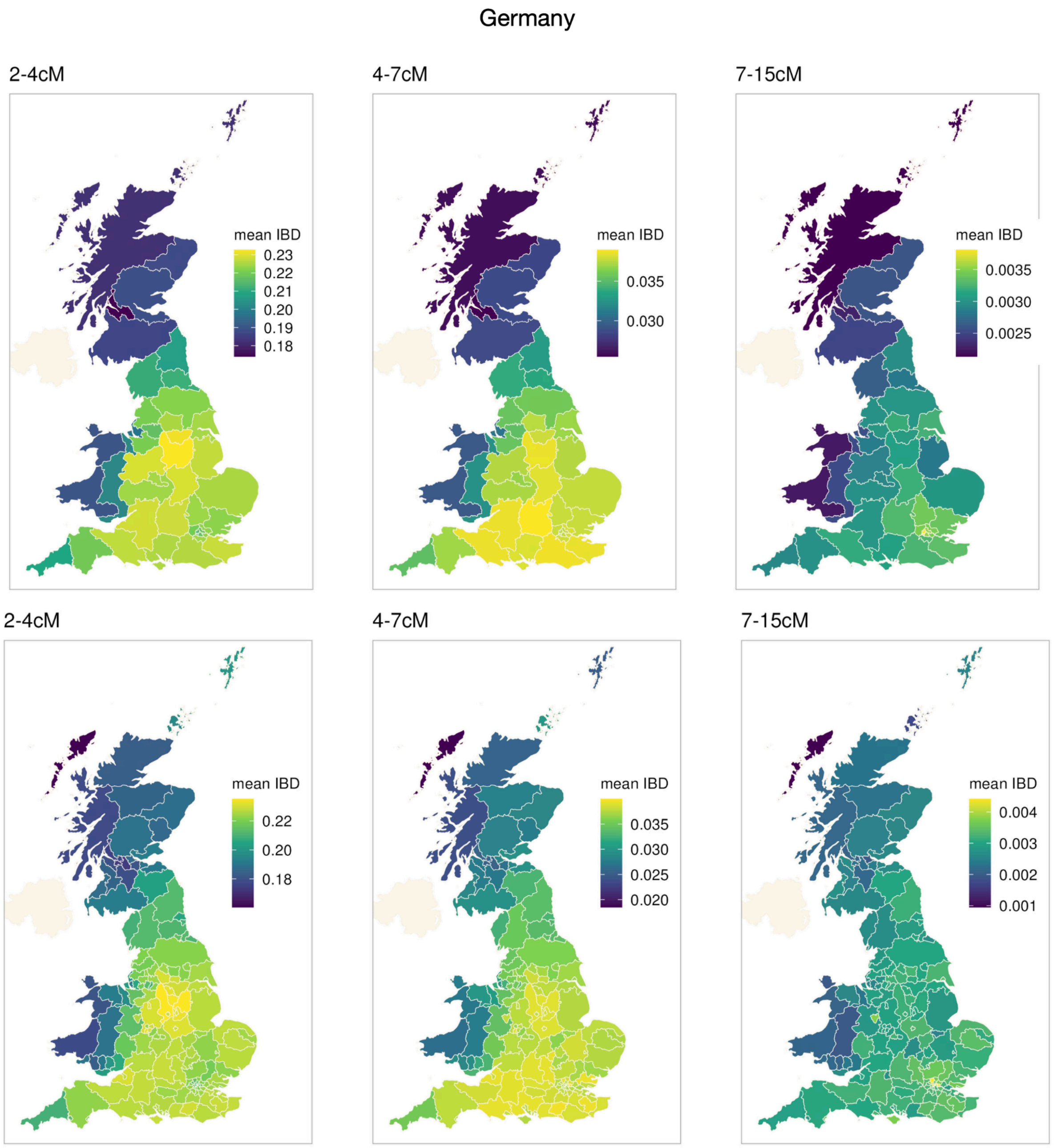


**Figure S39. Mean IBD sharing between the population of Germany and regions of the UK aggregated across varying lengths of IBD segments (unit: cM) is shown at the UK NUTS Level 2 (top) and NUTS Level 3 (bottom).** Derbyshire shows high IBD sharing with Germany for segments between 2–4 cM (MAP age = 555AD - 1255AD, 50% credible interval =257 BC - 1549AD). For segments between 4–15 cM (MAP age = 1255AD 1768AD, 50% credible interval = 849AD-1843AD), the hotspot shifts toward London and surrounding areas in the South East of England.

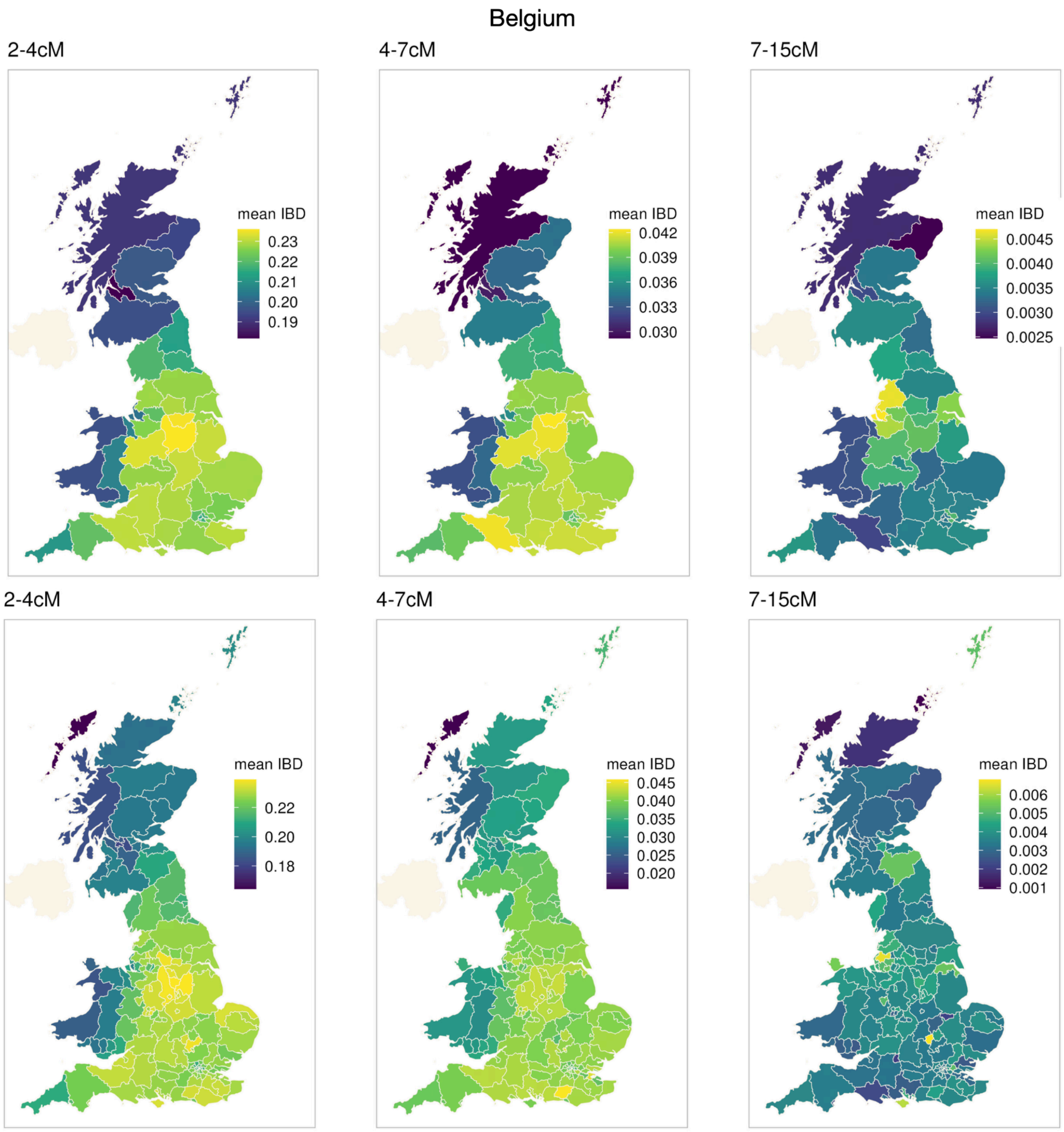


**Figure S40. Mean IBD sharing between the population of Belgium and regions of the UK aggregated across varying lengths of IBD segments (unit: cM) is shown at the UK NUTS Level 2 (top) and NUTS Level 3 (bottom).** Derbyshire has high IBD sharing with Belgium for segments between 2-4cM (MAP age = 555AD - 1255AD, 50% credible interval =257 BC - 1549AD). Between 4-7cM (MAP age=1255AD-1555AD, 50% credible interval = 849AD-1717AD), Belgium has increased IBD sharing with Dorset and Somerset (South West, England). Between 7-15cM (MAP age=1555AD-1768AD, 50% credible interval = 1325AD-1843AD), we also see increased IBD sharing with Lancashire and Merseyside (North West, England) and North Lincolnshire (East Midlands, England).

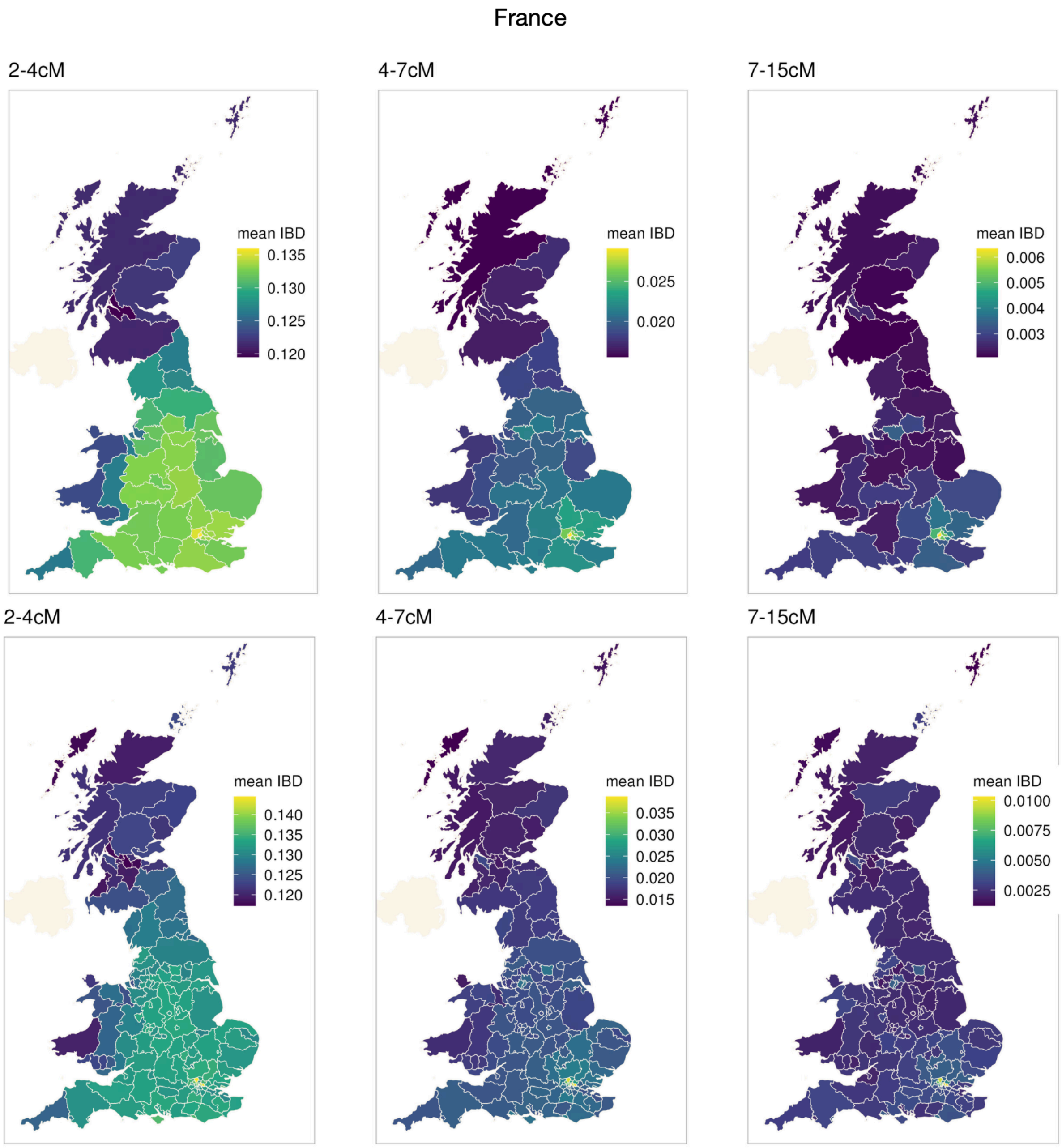


**Figure S41. Mean IBD sharing between the population of France and regions of the UK aggregated across varying lengths of IBD segments (unit: cM) is shown at the UK NUTS Level 2 (top) and NUTS Level 3 (bottom).** London has high IBD sharing with the French consistently throughout all periods.

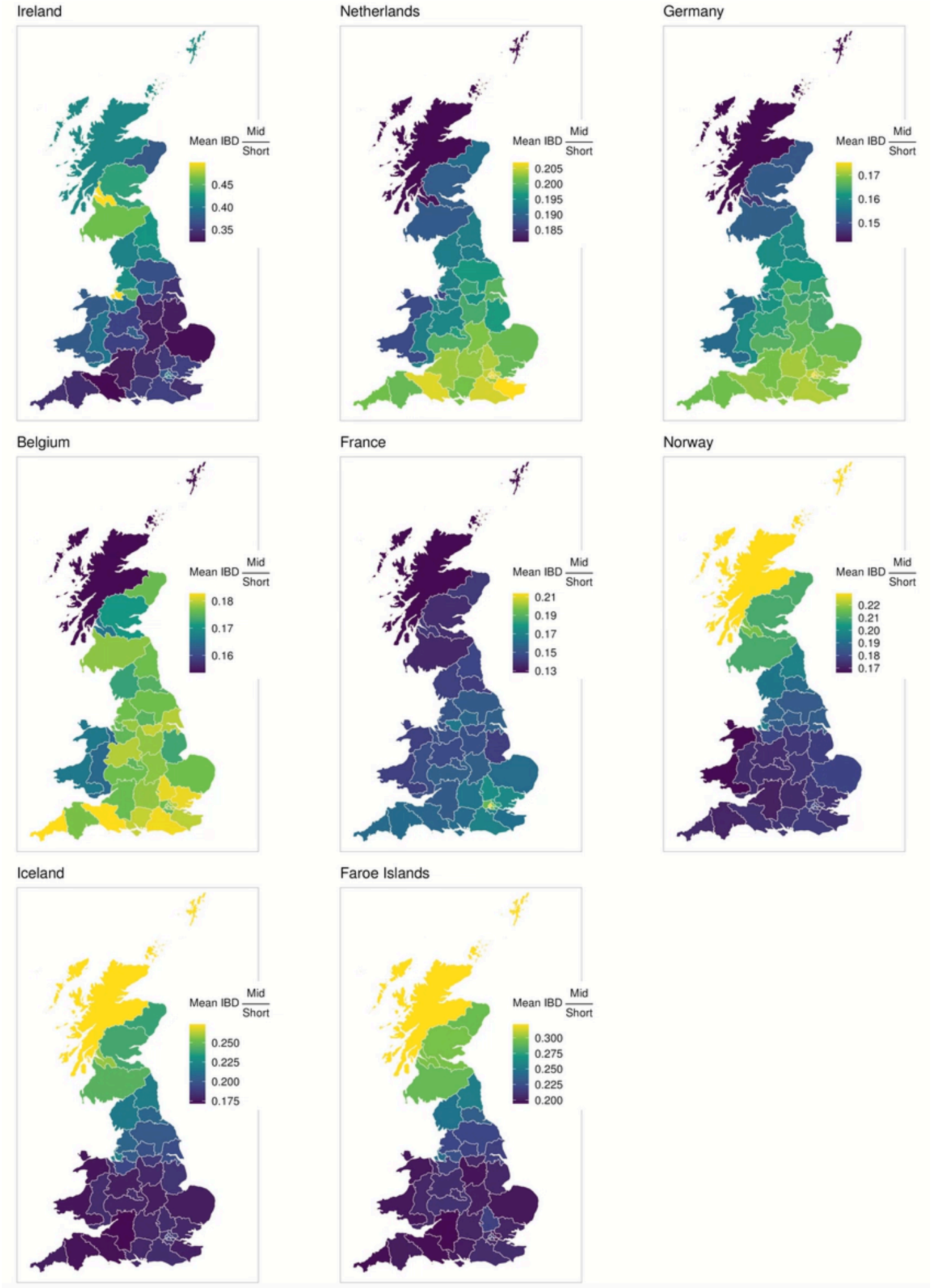


**Figure S42. Comparison of mean IBD sharing between British and the other neighbouring European Populations aggregated over medium-sized IBD segments (4-7cM) to short segments (2-4cM).** The ratio of mean IBD sharing aggregated over medium to short segments was plotted for each country across regions of the UK. Regions with a high ratio (shaded more yellow) indicate greater IBD sharing with that country's population for mid-range IBD segments (4-7cM), compared to short-range segments (2-4cM).

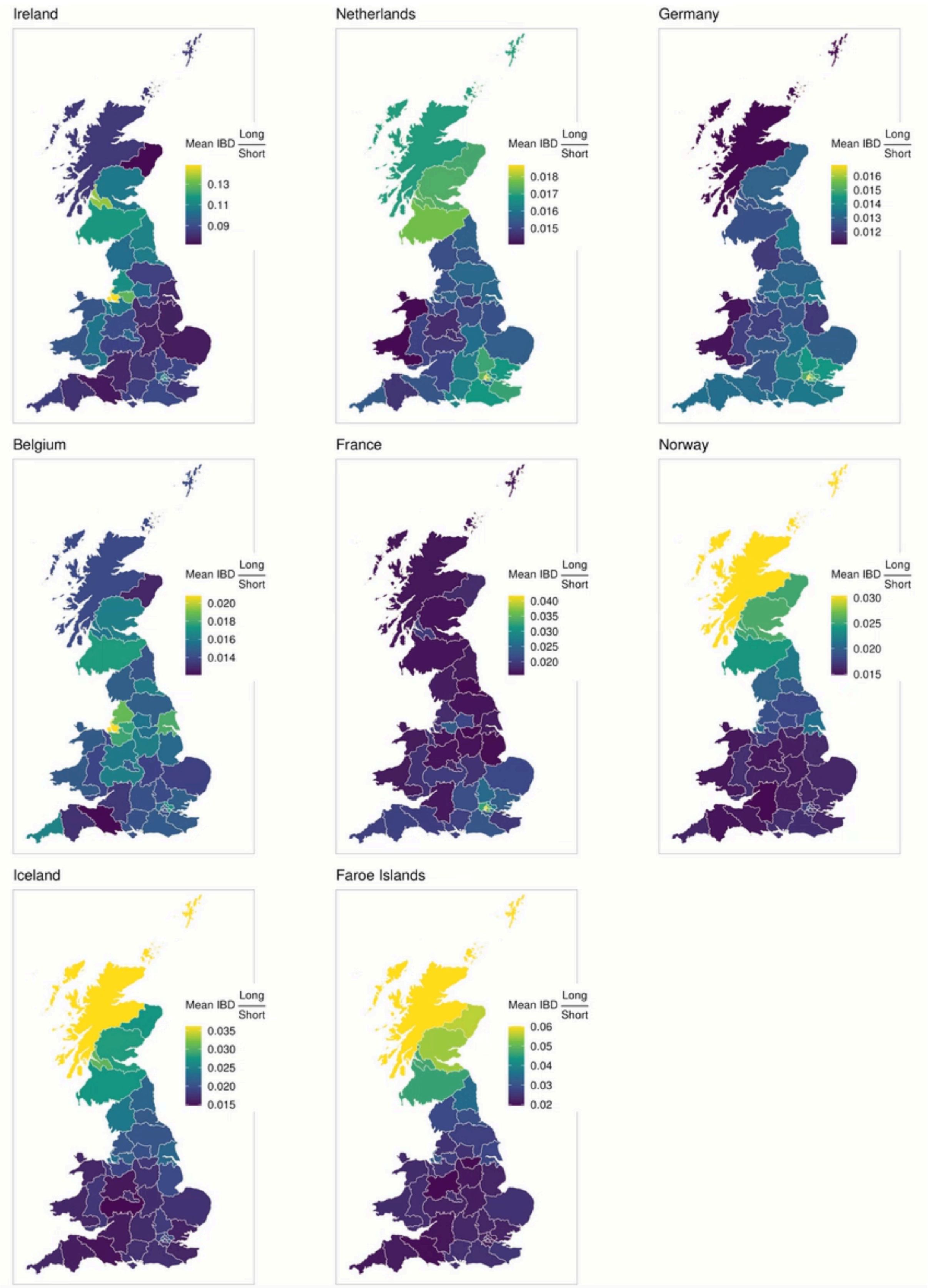


**Figure S43. Comparison of mean IBD sharing between British and the other neighbouring European Populations aggregated over long IBD segments (7-15cM) to short segments (2-4cM).** The ratio of mean IBD sharing aggregated over long to short segments was plotted for each country across regions of the UK. Regions with a high ratio (shaded more yellow) indicate greater IBD sharing with that country's population for long-range IBD segments (7-15cM), compared to short-range segments (2- 4cM).

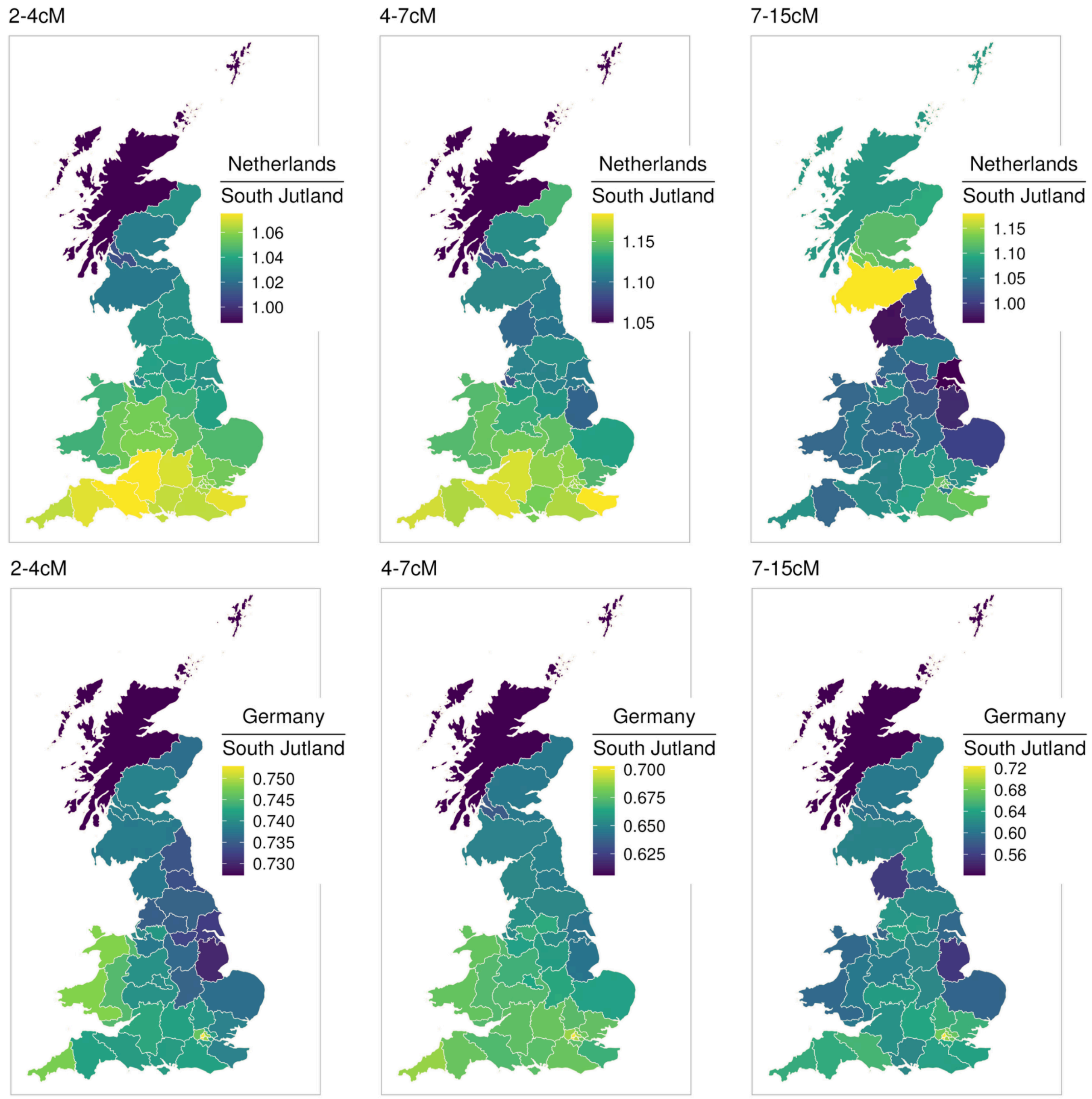


**Figure S44. Comparison of Britain's mean IBD Sharing with the Netherlands and Germany relative to South Jutland of Denmark.** Ratios of mean IBD sharing between Britain and the Netherlands (top) and Britain and Germany (bottom) relative to Britain and South Jutland were plotted across IBD segment length ranges. Ratios >1 indicate greater IBD sharing between Britain and the Netherlands or Germany compared to South Jutland; ratios <1 indicate the opposite.

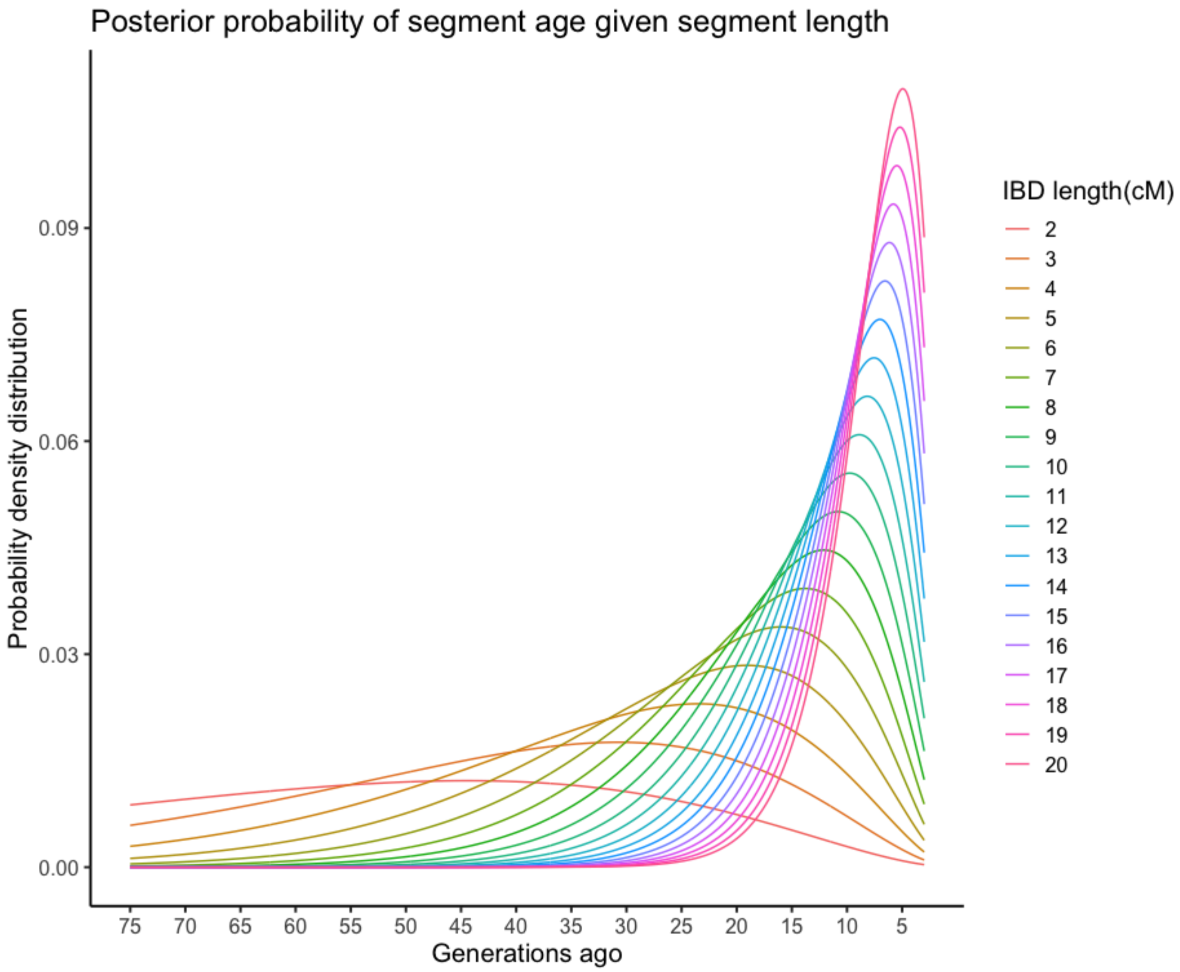


**Figure S45. Posterior density distribution of segment age given different segment lengths.**

### Supplementary Tables

**Table S1 - Mean IBD sharing among pairs of samples**

| Pair | Mean Total IBD sharing (cM) | Mean Total Number of IBD segments |
|---|---|---|
| Denmark-Denmark | 1.40 | 0.40 |
| Faroe Islands - Faroe Islands | 57.40 | 1.66 |
| England-England | 0.64 | 0.21 |
| Scotland-Scotland | 1.28 | 0.37 |
| Wales-Wales | 1.70 | 0.47 |
| Denmark-Faroe Islands | 0.95 | 0.27 |
| Denmark-England | 0.36 | 0.15 |
| Denmark-Scotland | 0.30 | 0.12 |
| Denmark-Wales | 0.32 | 0.13 |
| England-Scotland | 0.54 | 0.18 |
| Scotland-Wales | 0.43 | 0.16 |
| England-Wales | 0.60 | 0.20 |
| Faroe Islands - England | 0.37 | 0.14 |
| Faroe Islands - Scotland | 0.46 | 0.17 |
| Faroe Islands - Wales | 0.32 | 0.13 |

**Table S2 - Number of ancestor-offspring relationships extracted from the MGR-lite registry**

| Number of separating meiosis | Number of ancestor-offspring relationship pairs | Number of unique ancestors | Number of ancestor-offspring relationship pairs overlapped with genetic data |
|---|---|---|---|
| 1 | 9,394,359 | 4,450,371 | 40,057 |
| 2 | 8,972,463 | 2,417,374 | 16,091 |
| 3 | 3,510,568 | 854,864 | 181 |
| 4 | 172,067 | 50,914 | 0 |
| 5 | 1,076 | 445 | 0 |
| 6 | 4 | 2 | 0 |

**Table S3 - Number of Cousins extracted from the MGR-lite registry**

| Minimal number of separating meiosis | Number of pairs to be validated |
|---|---|
| 2 | 20,200 |
| 3 | 18,443 |
| 4 | 13,767 |
| 5 | 1,337 |
| 6 | 342 |

**Table S4. Mean total ROHs of Danish municipalities**

| Municipality | Mean totalROH EUR (cM) | Mean totalROH nonEUR (cM) | Sample size EUR | Sample size nonEUR |
|---|---|---|---|---|
| Aalborg | 1.96 | 4.10 | 4564 | 207 |
| Aarhus | 1.33 | 7.88 | 7284 | 464 |
| Albertslund | 1.39 | 72.38 | 303 | 127 |
| Allerød | 1.25 | 23.66 | 415 | 32 |
| Assens | 1.49 | 7.87 | 563 | 15 |
| Ballerup | 1.33 | 15.66 | 621 | 112 |
| Billund | 2.57 | 45.64 | 345 | 11 |
| Bornholm | 3.83 | 1.45 | 1703 | 86 |
| Brøndby | 1.82 | 51.90 | 423 | 111 |
| Brønderslev | 3.14 | 14.74 | 737 | 19 |
| Egedal | 1.60 | 11.95 | 539 | 56 |
| Esbjerg | 2.00 | 4.55 | 1823 | 115 |
| Faaborg-Midtfyn | 2.32 | 1.03 | 937 | 39 |
| Favrskov | 2.15 | 0.69 | 654 | 16 |
| Faxe | 2.73 | 3.92 | 1035 | 59 |
| Fredensborg | 0.77 | 37.27 | 560 | 142 |
| Fredericia | 1.77 | 4.59 | 757 | 44 |
| Frederiksberg | 1.69 | 9.51 | 8400 | 723 |
| Frederikshavn | 2.89 | 5.33 | 1290 | 51 |
| Frederikssund | 1.43 | 20.19 | 987 | 94 |
| Furesø | 0.40 | 41.10 | 430 | 84 |
| Gentofte | 1.70 | 4.01 | 3563 | 394 |
| Gladsaxe | 1.39 | 17.79 | 1803 | 205 |
| Glostrup | 1.37 | 16.11 | 2177 | 225 |
| Greve | 1.72 | 41.04 | 415 | 76 |
| Gribskov | 1.89 | 6.83 | 904 | 66 |
| Guldborgsund | 1.83 | 6.38 | 2645 | 183 |
| Halsnæs | 2.46 | 15.06 | 703 | 89 |

| | | | | |
|---|---|---|---|---|
| Hedensted | 2.77 | 0.25 | 444 | 12 |
| Helsingør | 1.46 | 28.16 | 1847 | 254 |
| Herlev | 0.73 | 39.13 | 452 | 70 |
| Herning | 2.82 | 23.27 | 1634 | 48 |
| Hillerød | 1.21 | 20.86 | 1650 | 170 |
| Hjørring | 3.06 | 1.00 | 1829 | 53 |
| Høje-Taastrup | 1.64 | 55.67 | 884 | 174 |
| Hørsholm | 1.16 | 12.09 | 1000 | 96 |
| Holbæk | 1.83 | 7.59 | 1825 | 119 |
| Holstebro | 2.93 | 18.80 | 1185 | 46 |
| Horsens | 1.57 | 9.13 | 1217 | 64 |
| Hvidovre | 1.57 | 47.18 | 720 | 162 |
| Ikast-Brande | 2.87 | 30.22 | 489 | 23 |
| Ishøj | 1.56 | 81.30 | 208 | 111 |
| Jammerbugt | 3.53 | 0.50 | 675 | 22 |
| Kalundborg | 2.91 | 18.35 | 1502 | 96 |
| Kerteminde | 0.70 | 0.17 | 299 | 12 |
| København | 1.91 | 17.46 | 31844 | 3944 |
| Køge | 1.20 | 23.47 | 1176 | 108 |
| Kolding | 1.98 | 0.44 | 1186 | 41 |
| Langeland | 3.54 | 1.00 | 370 | 13 |
| Lejre | 2.42 | 0.09 | 397 | 23 |
| Lemvig | 6.09 | 2.00 | 538 | 15 |
| Lolland | 1.84 | 4.97 | 1832 | 135 |
| Lyngby-Taarbæk | 2.10 | 15.15 | 1712 | 179 |
| Mariagerfjord | 2.40 | 8.27 | 902 | 37 |
| Middelfart | 1.94 | 1.35 | 592 | 23 |
| Morsø | 4.01 | 0.67 | 566 | 15 |
| Næstved | 1.68 | 8.73 | 2318 | 147 |
| Norddjurs | 2.87 | 1.63 | 1026 | 41 |
| Nordfyns | 2.89 | 1.07 | 470 | 15 |
| Nyborg | 2.25 | 2.19 | 534 | 27 |

| | | | | |
|---|---|---|---|---|
| Odder | 1.53 | 0.64 | 460 | 25 |
| Odense | 1.28 | 21.07 | 3103 | 246 |
| Odsherred | 3.65 | 8.83 | 777 | 29 |
| Randers | 1.85 | 4.90 | 1709 | 71 |
| Rebild | 2.45 | 1.19 | 491 | 16 |
| Ringkøbing-Skjern | 4.33 | 14.61 | 1092 | 28 |
| Ringsted | 1.06 | 37.11 | 801 | 65 |
| Rødovre | 1.08 | 6.67 | 1252 | 145 |
| Roskilde | 1.60 | 11.83 | 2398 | 176 |
| Rudersdal | 0.65 | 14.58 | 742 | 74 |
| Silkeborg | 1.91 | 5.89 | 1718 | 64 |
| Skanderborg | 1.73 | 0.33 | 772 | 30 |
| Skive | 4.05 | 3.13 | 1282 | 32 |
| Slagelse | 2.13 | 20.82 | 2356 | 155 |
| Solrød | 1.42 | 1.27 | 197 | 11 |
| Sorø | 2.91 | 0.32 | 678 | 28 |
| Stevns | 2.19 | 1.08 | 465 | 25 |
| Struer | 3.02 | 17.00 | 368 | 11 |
| Svendborg | 2.26 | 13.79 | 1279 | 62 |
| Syddjurs | 1.63 | 6.50 | 745 | 20 |
| Tårnby | 2.83 | 13.94 | 928 | 90 |
| Thisted | 3.42 | 0.67 | 1147 | 39 |
| Vallensbæk | 0.42 | 62.91 | 116 | 23 |
| Varde | 2.67 | 0.74 | 764 | 23 |
| Vejen | 3.18 | 1.62 | 501 | 13 |
| Vejle | 2.41 | 6.72 | 1581 | 65 |
| Vesthimmerlands | 2.59 | 1.04 | 802 | 25 |
| Viborg | 2.58 | 7.99 | 2071 | 81 |
| Vordingborg | 2.29 | 3.21 | 1366 | 96 |

Note: if a municipality (kommune) has less than 10 samples of either the EUR or non-EUR subset, we don't report it here.

**Table S5. The sample size of proxy populations from other European countries considered in the study**

| Country | Number of UKBB samples | Number of DNK samples | Total number of samples |
|---|---|---|---|
| Belgium | 74 | 38 | 112 |
| Finland | 148 | 146 | 294 |
| France | 454 | 120 | 574 |
| Germany | 818 | 1138 | 1956 |
| Iceland | 0 | 473 | 473 |
| Ireland | 3977 | 0 | 3977 |
| Netherlands | 349 | 216 | 565 |
| Norway | 104 | 1120 | 1224 |
| Poland | 558 | 1256 | 1814 |
| Sweden | 167 | 1056 | 1223 |

Note: The table displays the size of samples used as proxy populations for Britain/Denmark's neighbouring countries (after applying the Leiden community detection method). We reported the sample sizes recruited from the UKBB and the Danish (DNK) cohorts separately, as well as the combined total. For Iceland, samples are only available from the DNK cohorts. For Ireland, we only use samples collected from the UKBB.

**Table S6. Maximum a posteriori estimate of segment age and its 50% credible interval.**

| IBD length (cM) | MAP age (generation time) | MAP age (calendar time) | 50% credible interval (generation time) - lower bound | 50% credible interval (calendar time) - lower bound | 50% credible interval (generation time) - upper bound | 50% credible interval (calendar time) - upper bound |
|---|---|---|---|---|---|---|
| 2 | 50 | 555 AD | 29.0 | 1143 AD | 79 | 257 BC |
| 3 | 33.3 | 1022 AD | 20 | 1409 AD | 52.5 | 485 AD |
| 4 | 25.0 | 1255 AD | 15 | 1549 AD | 39.5 | 849 AD |
| 5 | 20.0 | 1395 AD | 12 | 1633 AD | 31.5 | 1073 AD |
| 6 | 16.7 | 1488 AD | 10 | 1675 AD | 26.5 | 1213 AD |
| 7 | 14.3 | 1555 AD | 9 | 1717 AD | 22.5 | 1325 AD |
| 8 | 12.5 | 1605 AD | 8 | 1745 AD | 20.0 | 1395 AD |
| 9 | 11.1 | 1644 AD | 7 | 1773 AD | 17.5 | 1465 AD |
| 10 | 10.0 | 1675 AD | 6 | 1787 AD | 15.5 | 1521 AD |
| 11 | 9.1 | 1700 AD | 6 | 1801 AD | 14.5 | 1549 AD |
| 12 | 8.3 | 1722 AD | 5 | 1815 AD | 13.0 | 1591 AD |
| 13 | 7.7 | 1740 AD | 5 | 1829 AD | 12.0 | 1619 AD |
| 14 | 7.1 | 1755 AD | 5 | 1829 AD | 11.5 | 1633 AD |
| 15 | 6.7 | 1768 AD | 4 | 1843 AD | 10.5 | 1661 AD |

Note:
The MAP estimates and 50% credible intervals (we used highest density intervals here) are derived assuming a constant and sufficiently large population size. Since we are talking about backwards in time, the lower bound indicates more recent generations, and the upper bound indicates more distant generations.

### Supplementary Note

#### Impact of complex demographic histories on the posterior distribution of segment age given segment length

In the main manuscript, we used the posterior distribution of segment age under a simplified demographic model assuming a constant population size. The prior distribution of segment age $f(g)$, is sensitive to these demographic assumptions. To evaluate how complex demographic histories could affect the age estimates, we performed ancestry simulations using msprime(Baumdicker et al. 2022; Kelleher, Etheridge, and McVean 2016; Nelson et al. 2020) to assess whether different demographic scenarios could qualitatively change the posterior distribution $f(g|l)$.

We simulated genealogical trees under the following demographic scenarios: (1) a single population with constant population size; (2) a single population with exponential growth at different rates; (3) a population bottleneck; (4) changes in migration rate between two populations. All the simulations were conducted on Chromosome 1 using the HapMap genetic map and the Discrete Time Wright-Fisher model, with an initial population size of 30,000 and a sample size of 200 individuals for single-population scenarios. In scenarios with migration, each population also has 200 samples. Simulations were conducted based on 2 haplotype copies per individual (ploidy =2). The lengths of the IBD segments shared across samples and the segment ages (i.e. corresponding ages of ancestors who transmitted the segments) were extracted from the simulated tree sequences. In particular, we compared two distributions shown in Figure S46: (1) $f(g|2 < l < 3)$, i.e. the posterior distribution of $g$

given IBD length = 2-3cM; (2) $f(g < TMRCA\,|2 < l < 3)$, i.e. the cumulative posterior distribution of $g$ being less than a given TMRCA, conditional on IBD length = 2-3cM.

For demographic scenarios with exponential growth, the scale of tested growth rates is chosen based on previous estimations for European populations reported in the literature(Tennessen et al. 2012; Fu et al. 2013; Gutenkunst et al. 2009). Compared to the model of constant population size, the mode of the posterior distribution $f(g|2 < l < 3)$ tends to shift toward older TMRCA values.

To test the impact of population bottlenecks, we introduced a reduction in population size between 10 and 20 generations ago, either to 50% or as low as 30% of the initial size. In these scenarios, the mode of the posterior distribution shifts toward more recent TMRCA values.

We also simulated the IBD sharing across two populations with episodes of mass migration history, implemented by adding a high symmetric migration rate (migration rate=0.5, i.e. 50% chance of having a parent from the other population during specific time windows, either a recent mass migration (10-15 generations ago) or an ancient mass migration (70-75 generations ago), while other time periods have a low background migration rate (rate=0.01). The mode of the posterior distribution shifts toward more recent generations in the recent mass migration scenario and toward older generations in the ancient migration scenario.

Given the complexity and inherent uncertainty in accurately estimating demographic parameters, we have chosen, for simplicity, to use the maximum a posteriori (MAP) estimate under a simple prior assuming constant population size, and we report the 50% credible interval. We caution that these estimates serve only as guidance and carry wide uncertainties.

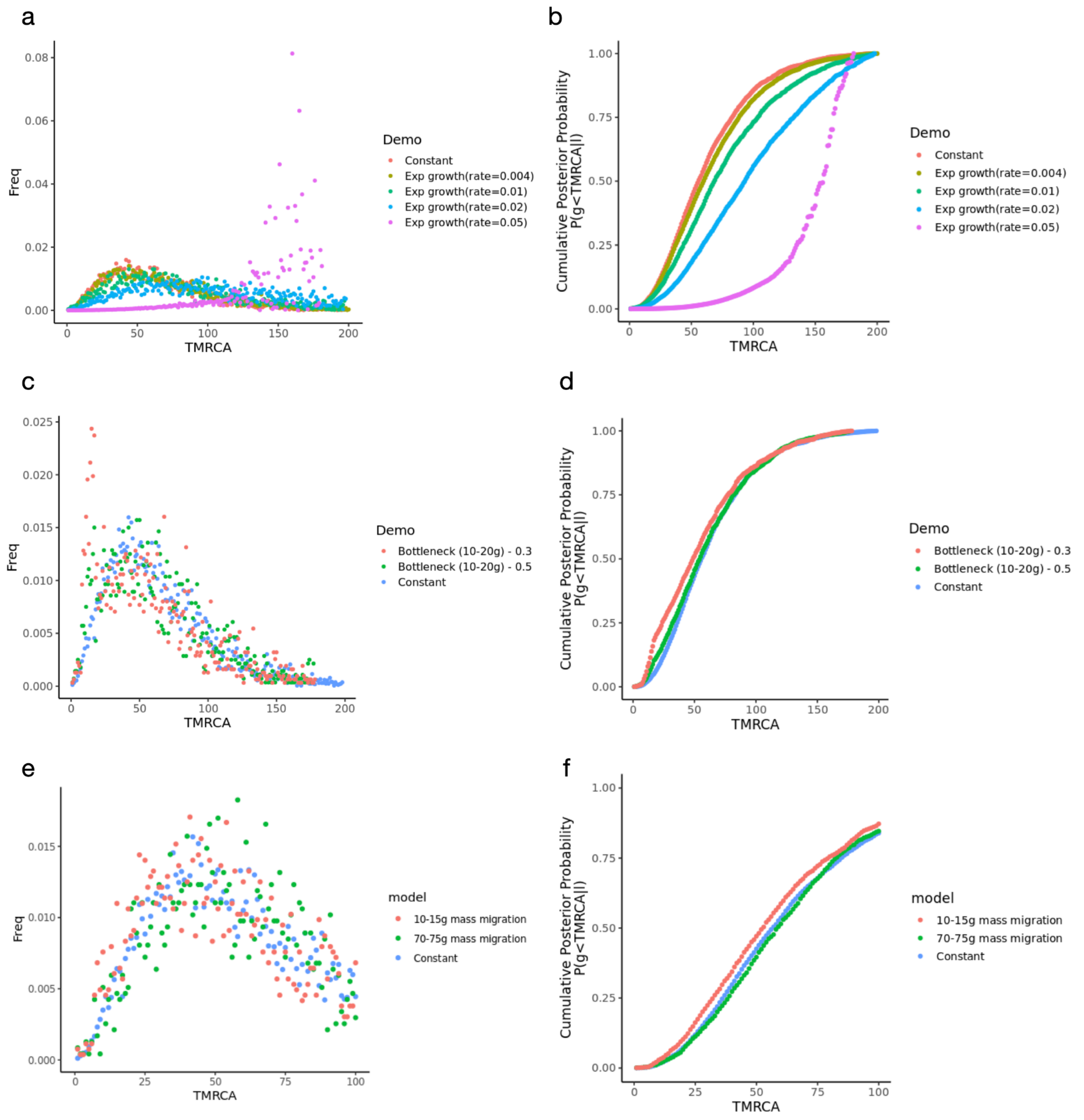


**Figure S46. Comparison of posterior distributions of segment age given IBD segment lengths (2–3 cM) under different demographic scenarios.** Each pair of panels shows results from the constant population size model (constant) and alternative demographic scenarios. The scenarios include: (1) single population with exponential growth at varying rates (a-b), (2) population bottleneck occurring 10–20 generations ago with varying severity (c-d), and (3) two-population models with symmetrical migration events at different time windows (recent mass migration 10–15 generations ago and ancient mass migration 70–75 generations ago) (e-f). Distributions shown in the left panel are the posterior distribution of segment age $f(g|2 < l < 3)$, and the right panel shows the cumulative posterior distribution

$f(g < TMRCA|2 < l < 3)$. Shifts in the mode of these distributions relative to the constant population model reflect the impact of demographic history on IBD-based age estimates.
